# Trinucleotide mRNA cap analog N6-benzylated at the site of posttranscriptional ^m6^Am mark facilitates mRNA purification and confers superior translational properties in vitro and in vivo

**DOI:** 10.1101/2023.11.10.566532

**Authors:** Marcin Warminski, Edyta Trepkowska, Miroslaw Smietanski, Pawel J. Sikorski, Marek R. Baranowski, Marcelina Bednarczyk, Hanna Kedzierska, Bartosz Majewski, Adam Mamot, Diana Papiernik, Agnieszka Popielec, Remigiusz A. Serwa, Brittany A. Shimanski, Piotr Sklepkiewicz, Marta Sklucka, Olga Sokolowska, Tomasz Spiewla, Diana Toczydlowska-Socha, Zofia Warminska, Karol Wolosewicz, Joanna Zuberek, Jeffrey S. Mugridge, Dominika Nowis, Jakub Golab, Jacek Jemielity, Joanna Kowalska

## Abstract

Eukaryotic mRNAs undergo co-transcriptional 5’-end modification with a 7-methylguanosine cap. In higher eukaryotes, the cap carries additional methylations, such as ^m6^A_m_ – a common epitranscriptomic mark unique to the mRNA 5’-end. This modification is regulated by the Pcif1 methyltransferase and the FTO demethylase, but its biological function is still unknown. Here, we designed and synthesized a trinucleotide FTO-resistant *N*6-benzyl analog of the ^m6^A_m_-cap – m^7^Gppp^Bn6^A_m_pG (termed *AvantCap*) and incorporated it into mRNA using T7 polymerase. mRNAs carrying ^Bn6^A_m_ showed several advantages over typical capped transcripts. The ^Bn6^A_m_ moiety was shown to act as an RP-HPLC purification handle, allowing separation of capped and uncapped RNA species, and to produce transcripts with lower dsRNA content than reference caps. In some cultured cells, ^Bn6^A_m_ mRNAs provided higher protein yields than mRNAs carrying A_m_ or ^m6^A_m_, although the effect was cell line-dependent. m^7^Gppp^Bn6^A_m_pG-capped mRNAs encoding reporter proteins administered intravenously to mice provided up to 6-fold higher protein outputs than reference mRNAs, while mRNAs encoding tumor antigens showed superior activity in therapeutic setting as anti-cancer vaccines. The biochemical characterization suggests several phenomena underlying the biological properties of *AvantCap*: (i) increased competitiveness of the mRNA 5’-end for eIF4E protein by reducing its propensity for unspecific interactions, (ii) direct involvement of eIF3 in alternative translation initiation, (iii) subtle differences in mRNA impurity profiles, or a combination of these effects. *AvantCapped-*mRNAs bearing the ^Bn6^A_m_ may pave the way for more potent mRNA-based vaccines and therapeutics and serve as molecular tools to unravel the role of the ^m6^A_m_ in mRNA.

## Introduction

Eukaryotic messenger RNAs (mRNAs) carry genetic information from the nucleus to the cytoplasm and serve as templates for protein biosynthesis in cells. As relatively short-lived and labile molecules, they are dynamically regulated on multiple levels, which plays a major role in controlling gene expression. Chemical modifications are utilized by both nature and researchers to modulate the biological properties of mRNA. One of the earliest discovered natural modifications of eukaryotic mRNA is the 7-methylguanosine 5’ cap, which in humans is accompanied by additional methylations at the 2’-*O* position of the first one or two transcribed nucleotides (Figure 1A).^1^ Although not yet fully understood, these methylations play a vital role as epigenetic marks by which the cell distinguishes between its own and foreign mRNAs during viral infection.^2, 3^ Analogous modifications are also introduced into exogenously delivered mRNA vaccines and therapeutics to make them resemble endogenous mRNA as much as possible.^4^ If adenosine is present as the first transcribed nucleotide (FTN) in mammalian mRNA, it can be additionally methylated at the *N*6-position to produce *N6*,2’-*O*-dimethyladenosine (^m6^A_m_).^5^ The methylation of adenine at the *N*6-position is a general regulatory mechanism in mRNA,^6^ but the biological effects of m^6^A presence strongly depend on its position in the mRNA body and the sequence context.^7^ The ^m6^Am in mRNA is only found at the 5’ end (as FTN) and has an as yet unclear biological function. It may serve as a basis for dynamic translation regulation mechanisms, relying on the addition and removal of the methyl group by m^6^A writers and erasers.^8, 9^ *N*-Methyltransferase Pcif1/CAPAM has been the only so far identified mRNA cap-specific ^m6^A writer, ^10-14^ while FTO has been identified as the ^m6^A eraser for both internal ^m6^A and ^m6^Am within the 5’ cap (Figure 1B).^15, 16^ However, the biological effects of m^6^Am and underlying molecular mechanisms are still under debate, since no cap-specific ^m6^Am reader has been identified. Interestingly, the *N*6-methylation of adenosine as FTN is common in all types of mammalian cells and has been found relatively abundant in several tissues: e.g., in mice the fraction of *N6-*methylated 5’ terminal A reaches around 70% in the liver, 90% in the heart, and 94% in the brain.^17, 18^ The large differences in ^m6^Am content observed in transcripts from different genes also support this modification’s putative regulatory role.^19, 20^

**Figure 1.**
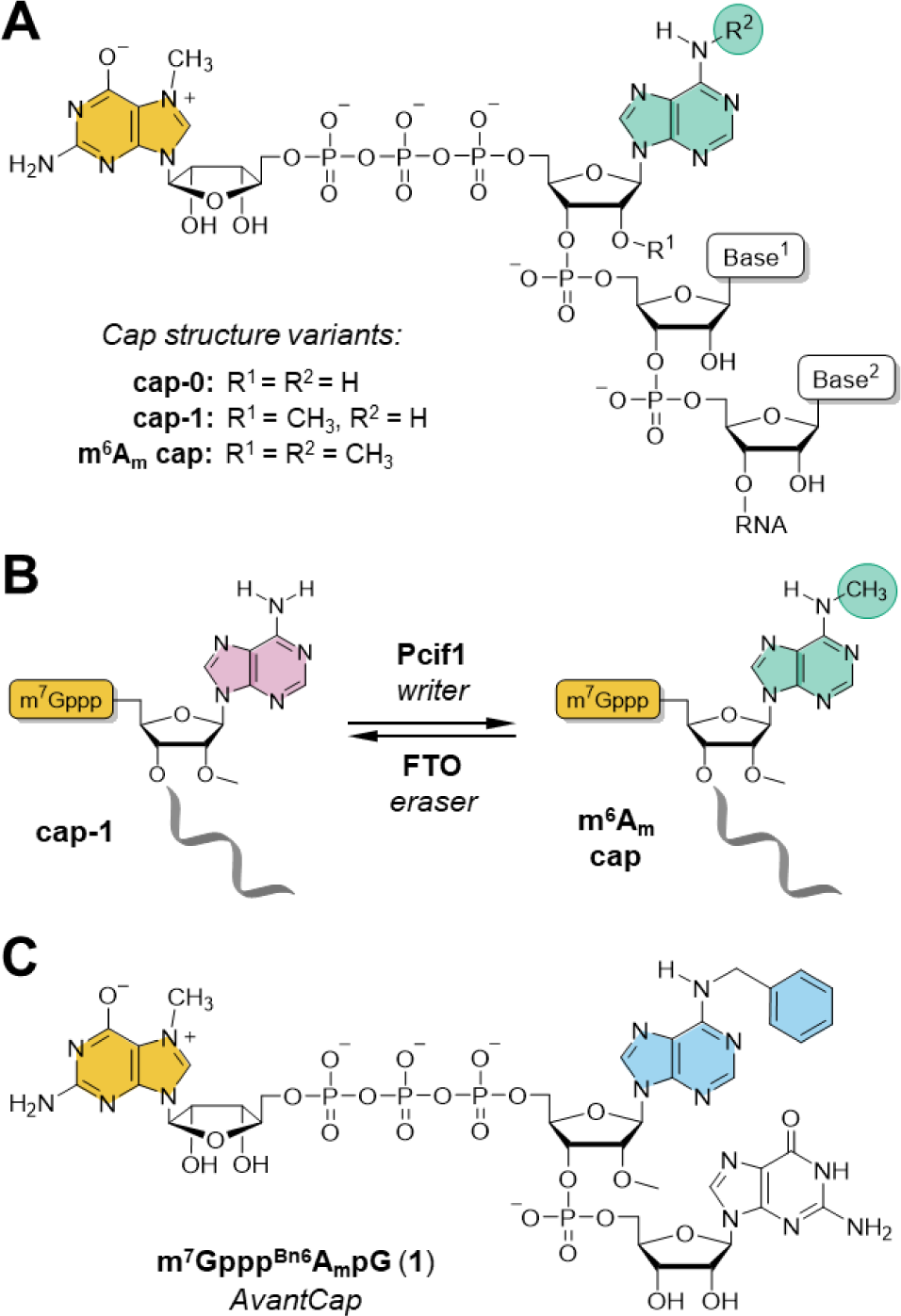
mRNA 5′ cap structure. (**A**) Natural variants of cap structures carrying adenosine adjacent to the 5’ cap; (**B**) Dynamic regulation of ^m6^Am presence in the cell; (**C**) Structure of m^7^Gppp^Bn6^AmpG (*AvantCap*).

Over the past few years, the molecular effect of the ^m6^Am modification on mRNA properties has been the subject of a lively scientific debate, with some conflicting data often resulting from the difficulty of separating the effects of 5′ cap-adjacent ^m6^Am and internal ^m6^A modifications. Most of the researchers agree that the level of cap-adjacent ^m6^Am positively correlates with mRNA translation rate,^9^ although that effect seems to be cell-line or tissue-dependent.^2, 21^ Direct comparison of bicistronic mRNAs capped with Am or ^m6^Am caps and containing IRES suggests that *N*6-methylation suppresses cap-dependent translation in rabbit reticulocyte lysate (RRL).^13^ To elucidate the biological role of ^m6^Am mark in a more complex system, Akichika et al., Boulias et al., and Sendinc et al. independently studied the PCIF1 knock-out cell lines.^10, 11, 13^ Under normal conditions, no difference in growth was observed between PCIF1 knock-out and wild-type cells, while under oxidative stress PCIF1-deficient cells showed defective growth,^10^ which suggests that ^m6^Am is important for survival under stress conditions. In line with this observation, Sun et al. reported elevated *N*6-methylation of the 5′-cap under heat shock and hypoxia conditions, particularly within the mRNAs coding for proteins engaged in stress-response mechanisms.^20^ Gene ontology analysis of mRNAs isolated from mouse liver revealed that ^m6^Am is clearly enriched in transcripts associated with mitochondrion and metabolic processes upon high-fat diet stress, linking the FTO activity with dynamic regulation of obesity.^22^ Studies in vivo showed that mice with mutations within the Pcif1 gene display reduced body weight, but their viability and fertility were unaffected. ^23^ A recent report has shown that *N6*-methylation of adenosine within the 5’ cap of viral mRNA attenuates the interferon-β–mediated suppression of viral infection,^24^ which suggests that this methylation may play a similar immunoregulatory role as 2’-*O*-methylations at the mRNA 5’ end.^2^ The methyltransferase activity of PCIF1 has also been shown to suppress HIV replication through enhancing the stability of host ^m6^Am modified transcripts, which is circumvented by the viral protein-mediated PCIF1 degradation.^25^ Other studies linked PCIF1 methyltransferase activity with susceptibility to SARS-CoV-2 and other coronavirus infections,^26^ as well as with the response to transforming growth factor beta (TGF-β) and to anti-PD-1 therapy in colorectal cancer cells.^27^

The ^m6^Am mark can be incorporated into in vitro transcribed (IVT) mRNA either by enzymatic posttranscriptional methylation^23, 28^ or co-transcriptional capping with properly designed priming nucleotide.^15, 20, 21, 29^ To enable more insight into the influence of ^m6^Am on the properties of mRNA in different biological settings, we have previously used a transcription-priming trinucleotide cap analog m^7^Gppp^m6^AmpG to evaluate translational properties of IVT mRNAs containing 5’ terminal ^m6^Am in different cell lines. ^21^ The presence of ^m6^Am did the not alter translation of reporter mRNA in mouse fibroblasts (3T3-L1), but increased translation efficiency in human cancer (HeLa) and mouse dendritic (JAWSII) cells (compared to Am). The observed translation up-regulation in specific cells, particularly in dendritic cells responsible for the generation of adaptive immunity, makes ^m6^Am an exciting candidate for improving the dynamically developing mRNA vaccine field.^30^ However, the potential reversibility of this modification makes it difficult to fully understand and harness its potential.

Synthetic modifications of mRNA 5′ cap have already proven to be an effective way to modulate translation efficiency of exogenously delivered transcripts.^31, 32^ However, unnatural modifications of the first transcribed nucleotide have been rarely explored so far.^19^ This is mostly because the typical protocols for the preparation of capped mRNA utilize dinucleotide cap analogs, which are not compatible with the majority of modifications at the FTN position.^33^ However, recently developed tri- and tetranucleotide capping reagents overcome these limitations.^2, 21, 34^ Here, we aimed to develop a capping reagent introducing an ^m6^Am mimic at the mRNA 5’ cap that would potentially retain the majority of its properties, but was resistant to demethylation by FTO. We envisaged that at the molecular level, methyl group can either stabilize complexes with proteins by hydrophobic interactions or destabilize them by steric hindrance. We hypothesized that both effects can be potentially enhanced by replacing ^m6^Am methyl group with a more bulky substituent such as benzyl, which has shown previously to be a good methyl mimic in terms of mRNA cap-protein interactions.^35^ Consequently, by replacing the *N*6-methyl group with benzyl in the m^7^Gppp^m6^AmpG cap structure we developed a novel type of mRNA capping reagent – m^7^Gppp^Bn6^AmpG (Figure 1C). After careful evaluation of m^7^Gppp^Bn6^AmpG in vitro, in cultured cells, and in vivo in mouse models, aided by biochemical and proteomic experiments, we found that ^Bn6^Am may indeed act as an FTO-resistant ^m6^Am mimic that enhances translational potential of IVT mRNA. Therefore, m^7^Gppp^Bn6^AmpG, which we termed *AvantCap*, is a highly promising reagent for modification of IVT mRNA with high potential to reveal the biological nuances of ^m6^Am functions as well as for therapeutic mRNA applications.

## RESULTS

### Chemical synthesis of AvantCap (**1**) – an mRNA cap analog containing N6-benzyl-2’-O- methyladenosine (^Bn6^Am)

The synthetic pathway leading to trinucleotide cap analog **1** included solid-phase synthesis of dinucleotide 5′-monophosphate **3a**, followed by its activation into *P*-imidazolide **4** and ZnCl2-mediated coupling reaction with 7-methylguanosine 5′-diphosphate (m^7^GDP) in solution (Scheme 1). The *N*6- benzyladenosine phosphoramidite for solid-phase synthesis was prepared by one-step alkylation of commercially available *N*6-phenoxyacetyl-2′-*O*-methyladenosine phosphoramidite in the presence of a base and phase-transfer catalyst.^17, 36^ The dinucleotide 5′-phosphate **3** was cleaved from the solid support and deprotected using standard protocols and then isolated by ion-exchange chromatography on DEAE Sephadex to give a triethylammonium salt suitable for further activation. The *P*-imidazolide **4** was prepared as described previously for mononucleotides,^37^ precipitated as a sodium salt and reacted with m^7^GDP in the presence of excess ZnCl2 to give trinucleotide cap analog **1**. Activation of dinucleotide **3a** instead of m^7^GDP appeared to be more efficient, particularly at larger scales (>50 µmol), and allowed to reduce the coupling time from 24–48 h to ca. 2 h. Compound **4** was relatively stable and was stored at -20–4°C for several months without signs of decomposition. The final product **1** was isolated by ion-exchange chromatography and additionally purified by RP-HPLC to give ammonium salts of **1** in good yield (70% starting from **3**). A typical solid-phase synthesis at 200 µmol scale required 1.2–1.5 equivalents of ^Bn6^Am phosphoramidite and yielded ca. 140 µmol of **3a**. We were able to upscale the procedure to obtain 3.15 g (ca. 2.5 mmol) of cap analog **1** starting from an equivalent of 5 mmol of solid-supported guanosine and 10.3 g (10.2 mmol) of 2′-*O-*methyladenosine phosphoramidite.

**Scheme 1.**
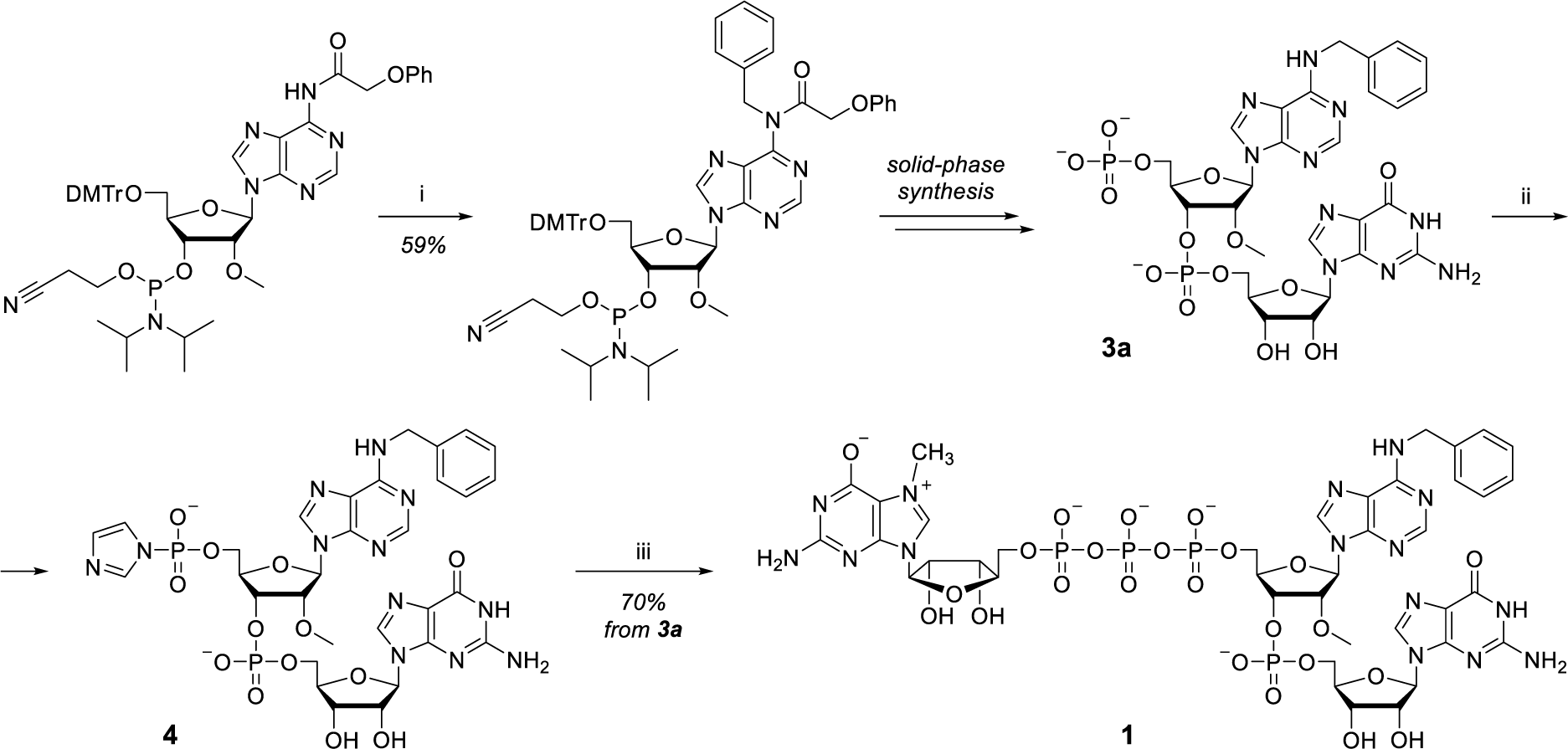
Chemical synthesis of m^7^Gppp^Bn6^AmpG (1). Reaction conditions: i) benzyl bromide, tetrabutylammonium bromide, 1M NaOHaq, CH2Cl2; ii) imidazole, 2,2′-dithiodipyridine, triphenylphosphine, triethylamine, DMF; iii) *N*7-methylguanosine 5′-diphosphate (m^7^GDP), ZnCl2, DMSO.

### Characterization of *AvantCap*

#### AvantCap initiates transcription by T7 polymerase to produce m^7^Gppp^Bn6^AmpG-capped RNA and facilitates mRNA purification by HPLC via hydrophobic effect

We have recently shown that *N*6-methylation of adenosine does not substantially impair the ability of trinucleotide cap analogs to prime in vitro transcription reaction (up to 80% of capping efficiency on short 35-nt RNAs obtained with m^7^Gppp^m6^AmpG *versus* 90% for those obtained with m^7^GpppAmpG).^21^ To assess how well is the more bulky *N*6-benzyl substituent accommodated by the T7 RNA polymerase, we performed an in vitro transcription (IVT) from DNA template containing T7 class III promoter (Φ6.5) sequence (Figure 2A) and coding for short RNAs under generic IVT conditions. The transcripts were trimmed by DNAzyme to reduce the heterogeneity of their 3’ ends,^38^ purified, and analyzed by gel electrophoresis to assess the ratio of capped and uncapped RNA (capping efficiency; for details see Experimental section). We found that the capping efficiency negatively correlates with the size of the *N*6-adenine substituent, but nonetheless m^7^Gppp^Bn6^AmpG was incorporated into 69% of the transcribed RNAs, compared to 97% for m^7^GpppAmpG and 82% for m^7^Gppp^m6^AmpG under the same conditions (Figure 2B). These observations can be explained by the fact that in order to pair with the thymidine of the DNA template, both ^m6^Am and ^Bn6^Am have to adopt the unfavorable anti conformation of the *N*6 substituent,^39^ which reduces the annealing rate.^40^

**Figure 2.**
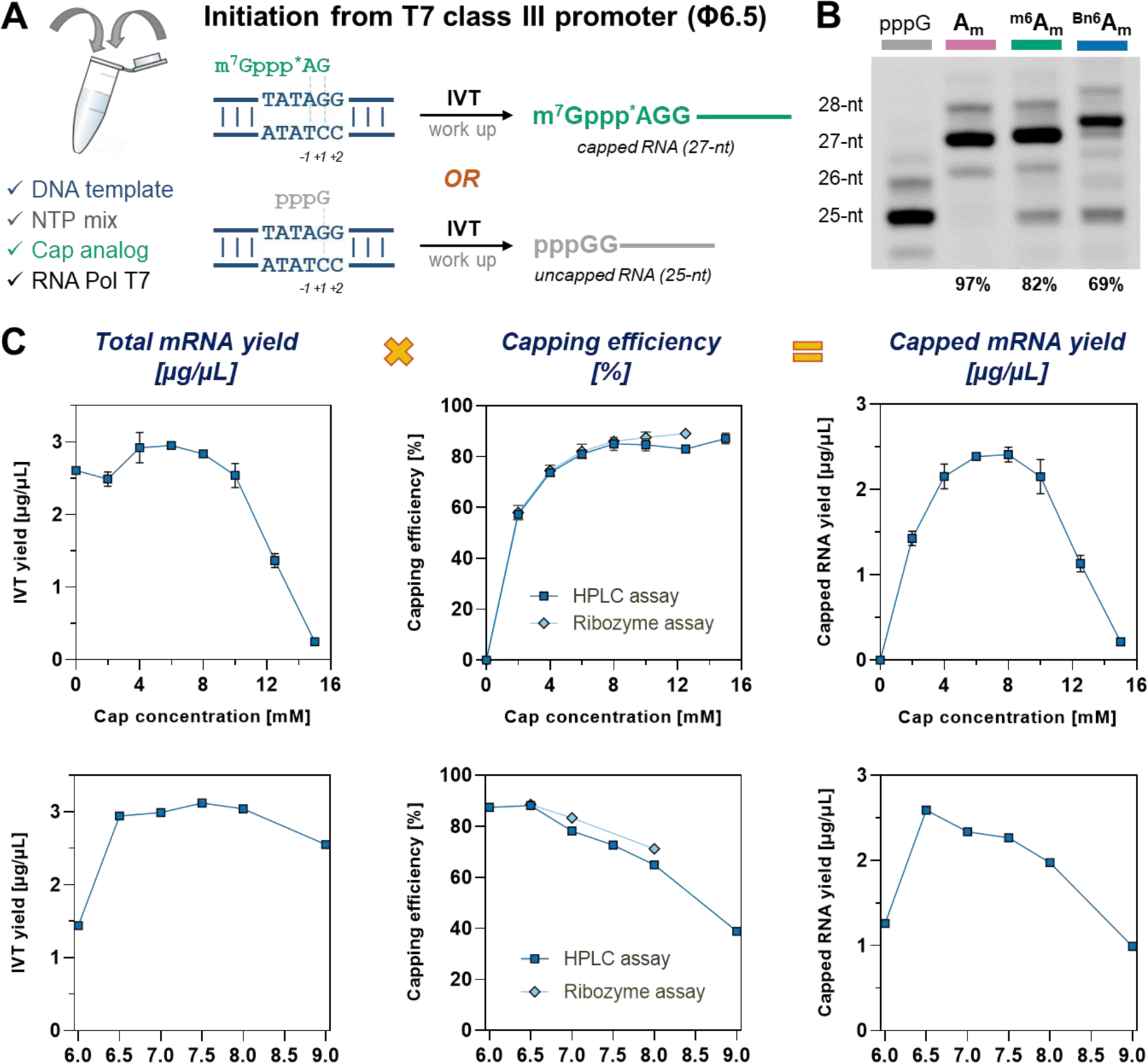
m^7^Gppp^Bn6^AmpG initiates transcription from T7 class III promoter (φ6.5) and GGG transcription start site. (**A**) Possible scenarios for initiation of transcription occurring in the presence of *AG-type trinucleotides (such as m^7^GpppApG) and the studied DNA template; *A denotes *N*6-modified adenosine residue (**B**) Capping efficiencies for short RNAs determined by gel electrophoresis. IVT reactions were performed in the presence of 1.25 µM template, 3 mM ATP, CTP, UTP and 0.75 mM GTP, 6 mM cap analog, at pH 7.9 (for details see Experimental section); (**C**) Capping efficiencies (determined by two methods) and IVT yields (determined spectrophotometrically after initial purification) as a function of *AvantCap* concentration and pH. IVT reactions were performed in the presence of 25 mM MgCl2, 40 ng/µL template, 5 mM ATP, CTP, UTP, 4 mM GTP and 10 mM cap analog (for optimizing pH) or various cap concentrations at pH 6.5.

We then moved on to full length mRNA (∼1000-nt) and attempted to optimize the IVT conditions to maximize both capping efficiency and RNA yield. Extensive optimization of the IVT reaction mix in terms of component concentration and buffer composition, revealed that cap concentration, pH of the buffer, and magnesium ions concentration are the most crucial factors affecting both capping efficiency and IVT yield (Figure 2C, Figure S1). Capping efficiencies reaching 90% were achieved under the optimized conditions (8–10 mM cap, pH 6.5, and 25 mM MgCl2) for model mRNA. The conditions provided IVT yields of 2.5 to 4.3 mg/mL and capping efficiencies from 80 to 90% for most of the studied mRNAs (Table S2).

When analyzing RNA integrity by HPLC we surprisingly found that m^7^Gppp^Bn6^AmpG-capped mRNA had significantly longer retention time than corresponding uncapped mRNA or mRNA carrying m^7^GpppAmpG or m^7^Gppp^m6^AmpG at the 5’ end (Figure 3A). This “hydrophobic effect” caused by the presence of benzyl group in mRNA was observed for different mRNAs up to 2000 nt in length, in each case allowing straightforward separation of capped and uncapped (5’-triphosphate) RNA (Figure 3B). A similar effect was observed before, but for much bulkier substituents, such as fluorescent tags ^41^ or photo-cleavable groups ^42^. This useful phenomenon can be harnessed for RP-HPLC purification of mRNAs and for direct assessment of capping efficiency without the typical processing of the mRNA sample, which relies on cleaving the 5’ terminal sequence by an enzyme, DNAzyme, or ribozyme followed by electrophoretic analysis. The capping efficiency values obtained by traditional ribozyme-based assay are in excellent agreement with values determined by the direct analysis of a small aliquot (0.5–1 µg) of mRNA by RP-HPLC (Figure 2C).

**Figure 3.**
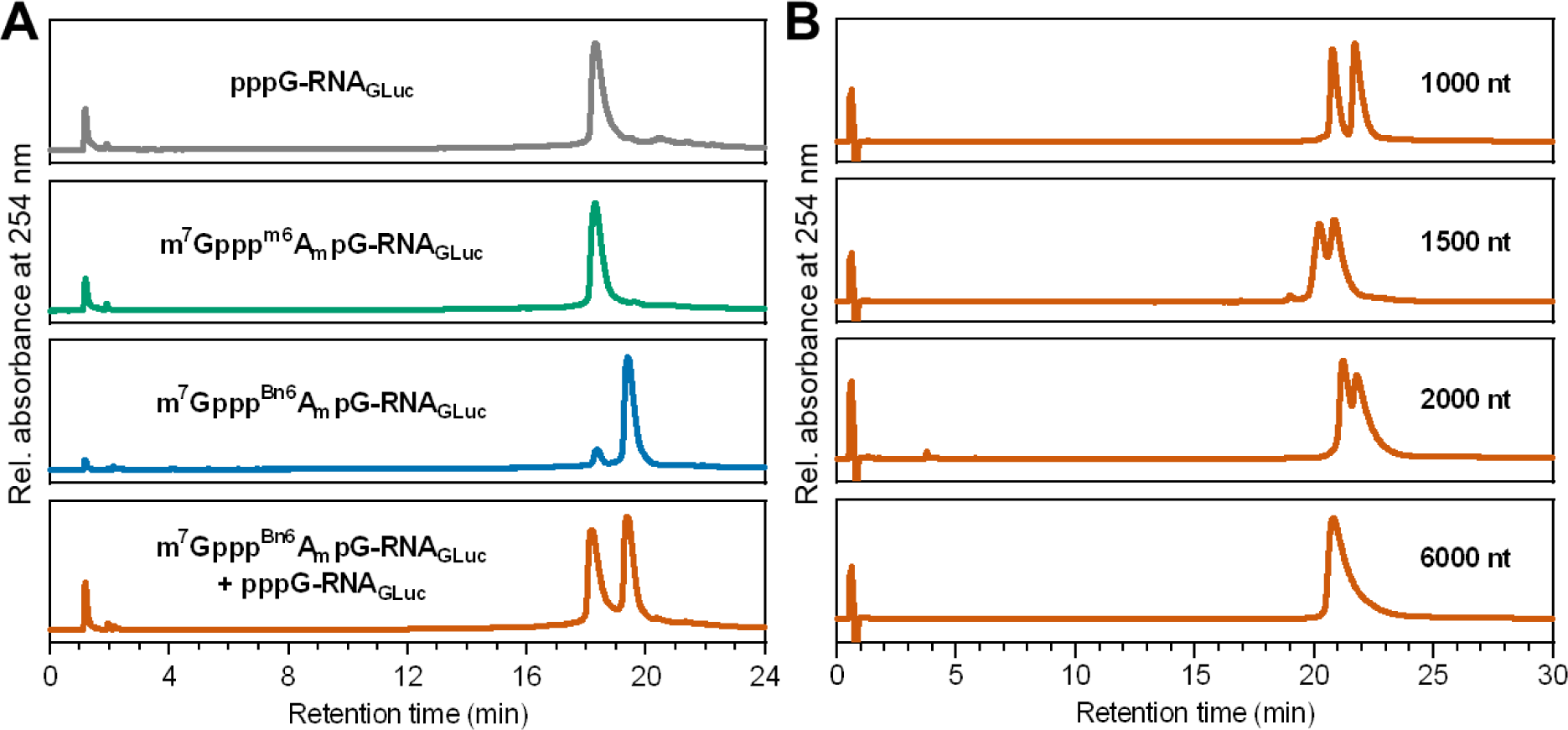
*N*6-Benzyl-2’-*O*-methyladenosine (^Bn6^Am) within the 5’ cap acts as an mRNA purification handle. (**A**) RP-HPLC analysis of *Gaussia* luciferase (Gluc) mRNA (956 nt) obtained by IVT carrying various 5’ terminal structures: uncapped mRNA (grey), m^7^Gppp^m6^AmpG-capped mRNA (green), m^7^Gppp^Bn6^AmpG-capped mRNA (blue), and 1:1 mixture of m^7^Gppp^Bn6^AmpG-capped mRNA and pppG-mRNA (orange) revealing that the presence of ^Bn6^Am delays mRNA retention and enables separation of capped mRNA (*R*t = 19.4 min) and uncapped mRNA (*R*t = 18.2 min). RP-HPLC conditions are given in the Experimental section. (**B**) RP-HPLC analyses of ∼1:1 mixtures of m^7^Gppp^Bn6^AmpG-capped transcripts of different lengths (1000, 1500, 2000, and 6000 nt) and corresponding uncapped mRNAs. RP-HPLC conditions are given in the Experimental section. ^Bn6^Am at the transcription start site increases mRNA translation in certain cell lines

Additionally, we have noticed that initially purified mRNAs co-transcriptionally capped with *AvantCap* contain reduced amount of dsRNA impurities in comparison to analogous transcripts with m^7^GpppAmpG (Figure S2). This feature was observed regardless from the applied purification method (affinity chromatography by oligo(dT)25 resin or purification using cellulose) or mRNA length and sequence composition. Therefore, we conclude that *AvantCap* can be harnessed to produce mRNAs with low dsRNA content.

After optimizing the IVT and purification protocols, we prepared a series of mRNAs encoding different reporter proteins (Table S1, Table S2) and tested them for translational activity in various cultured cell lines to gain first insights into biological activity of mRNAs carrying *AvantCap*. All mRNAs were purified by affinity chromatography followed by RP-HPLC and their purities, capping efficiencies, and homogeneities were verified (Table S2). Firefly luciferase (Fluc) encoding mRNAs capped with m^7^Gppp^Bn6^AmpG, m^7^Gppp^m6^AmpG, or m^7^GpppAmpG (Table S2) were transfected into colorectal cancer (CT26), human lung carcinoma (A549), and human kidney embryonic (HEK293T) cells and the luminescence dependent on Fluc protein expression was determined (Figure 4A). We found that the translational properties of mRNAs were cell culture dependent. In CT26 cells protein expression from mRNAs capped with m^7^Gppp^Bn6^AmpG was higher than those capped m^7^Gppp^m6^AmpG, and m^7^GpppAmpG-capped mRNA, while in HEK293T and A549 cells the protein levels were comparable. Next, we prepared a series of human erythropoietin (hEPO) encoding mRNAs capped with m^7^Gppp^Bn6^AmpG or m^7^GpppAmpG (Table S2) and transfected them into primary bone marrow (BM)-derived murine macrophages, primary bone marrow (BM)-derived dendritic cells or HEK293T cells. We observed increased protein expression in both primary murine cells but not in HEK293T cells (Figure 4B). Finally, we also prepared *Gaussia* luciferase (Gluc) encoding mRNAs capped with either m^7^Gppp^Bn6^AmpG or m^7^GpppAmpG, and compared their translation at different mRNA doses in human dendritic cells differentiated from monocytes of three healthy adult males (Figure 4C, Figure S5). In each case, we observed 2- to 10-fold higher expression of the reporter protein from mRNAs capped with m^7^Gppp^Bn6^AmpG.

**Figure 4.**
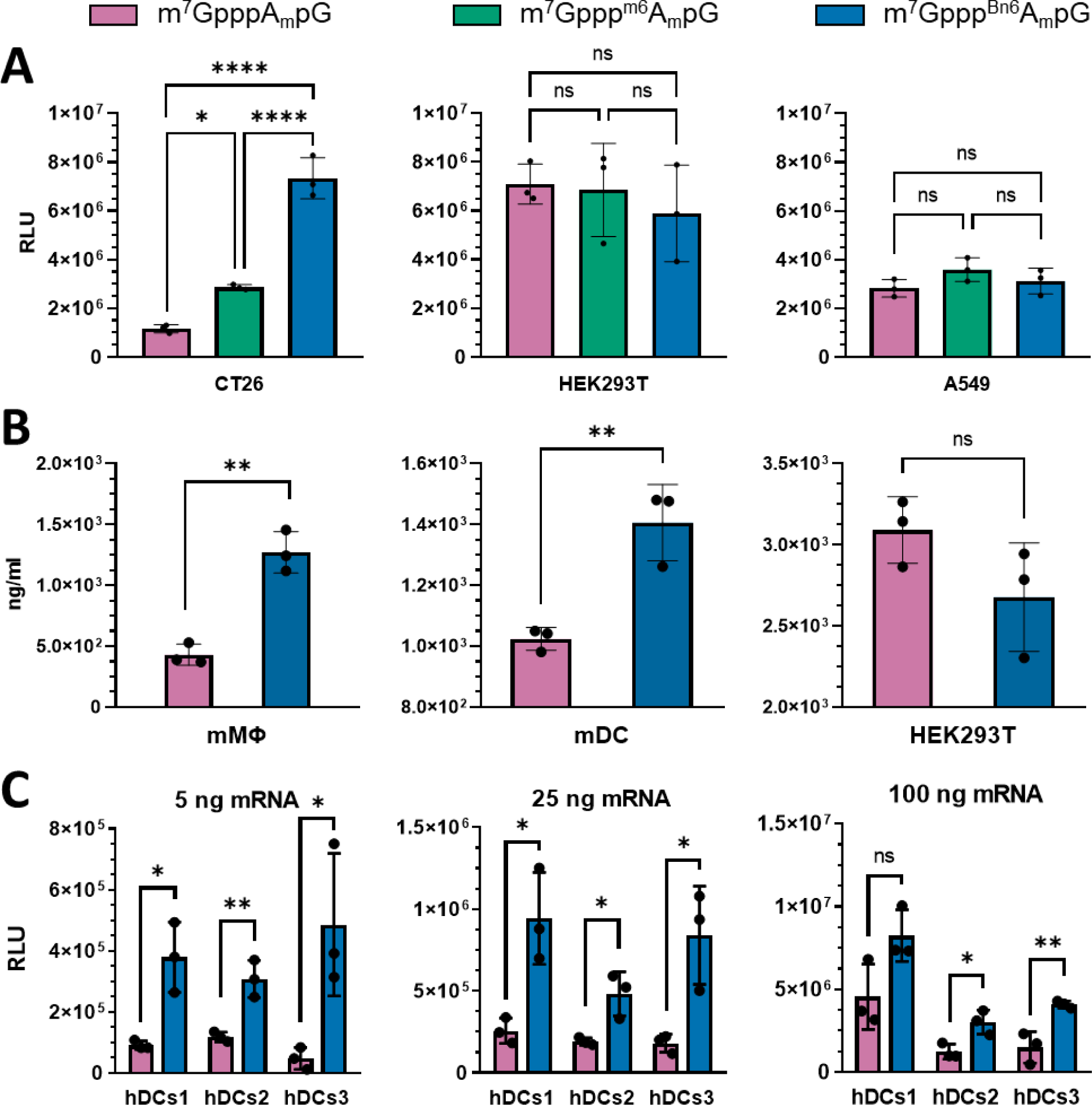
*N*6-benzyladenosine within mRNA 5’ cap increases protein expression in certain mammalian cells. (**A**) Firefly luciferase (Fluc)-dependent luminescence of murine colon carcinoma (CT26) and human embryonic kidney (HEK293T) and human Lung Carcinoma (A549) cells 6 h post transfection with 50 ng of Fluc mRNA; data show relative luminescence units (RLU) means ± SD, n=3, * P<0.05, **** P<0.001, ns – not significant, one-way ANOVA with Tukey’s multiple comparisons test. (**B**) Human erythropoietin (hEPO) concentration in culture medium of primary murine bone marrow (BM)-derived macrophages (mMΦ), murine BM-derived dendritic cells (mDC) and human embryonic kidney (HEK293T) cells 24 h post transfection with 200 ng of hEPO mRNA; data show hEPO concentrations (ng/mL) with means ± SD, n=3, **P<0.01, ns – not significant, two-tailed unpaired *t* test. (**C**) *Gaussia* luciferase (Gluc)-dependent luminescence of human monocyte-derived dendritic cells (hDCs) transfected with 5, 25 or 100 ng of Gluc mRNA; data show total protein production (sum of relative luminescence units [RLU] in daily measures from day 1 till 6 post transfection) with means ± SD, n=3, * P<0.05, **P<0.01, ns – not significant, two-tailed unpaired *t* test.

#### ^Bn6^Am at the transcription start site increases mRNA translation in vivo in mice

Next, we investigated how the presence of the benzyl modification affects mRNA expression in vivo. First, we used the wild-type Fluc reporter mRNAs capped with m^7^GpppAmpG or m^7^Gppp^Bn6^AmpG. mRNA purity and integrity was confirmed before each experiment (Figure S3, Table S2). Then, mRNAs were formulated into lipid nanoparticles (LNPs) using various ionizable lipids (GeneVoy-ILM^TM^, SM-102, or MC3) or complexed with commercially-available transfection reagent (TransIT®), and administered intravenously (i.v.) into mice. The luciferase activity was determined at multiple time points (Figure 5A,B; Figure S6). The absolute and the relative activities of differently capped mRNAs depended on the formulation type. The highest expression of a reporter gene was observed in mice treated with mRNAs formulated in SM-102 lipid, which also showed notable difference between m^7^GpppAmpG and m^7^Gppp^Bn6^AmpG-capped mRNAs (over 6-fold higher expression from ^Bn6^Am RNA at 4 h and 8 h time points and almost 3-fold at 24 h; Figure 5). Significantly higher expression was also observed when mRNAs were delivered with TransIT® (Figure S6), while for other formulations the differences were statistically significant for selected time points only (Figure 5). We have then used a different reporter protein that can be quantified using ELISA. To that end, we compared protein outputs from mRNAs encoding hEPO as a model of a therapeutically-relevant protein (Figure 5C). Again, purified mRNAs (Figure S4) formulated with SM-102 lipid provided the highest expression, but in this case the advantage of *AvantCap* over unmodified cap-1 was evident in all formulations at all time points. At 4 h post injection of SM-102 LNPs, the concentration of hEPO in mice blood serum was 6-fold higher in mice treated with ^Bn6^Am-capped mRNA than in those treated with m^7^GpppAmpG-RNA, and the ratio increased over time. Using a different potentially therapeutic mRNA, we detected over 5-fold higher α1-antitrypsin (hA1AT) concentrations in the sera of mice inoculated i.v. with TransIT®-formulated hA1AT-encoding mRNAs capped with m^7^Gppp^Bn6^AmpG as compared with m^7^GpppAmpG-capped mRNA (Figure S7D). hA1AT levels were also increased when HEK293T and A549 cells were transfected with mRNA capped with m^7^Gppp^Bn6^AmpG compared to m^7^GpppAmpG-capped mRNA (Figure S7A-C).

**Figure 5.**
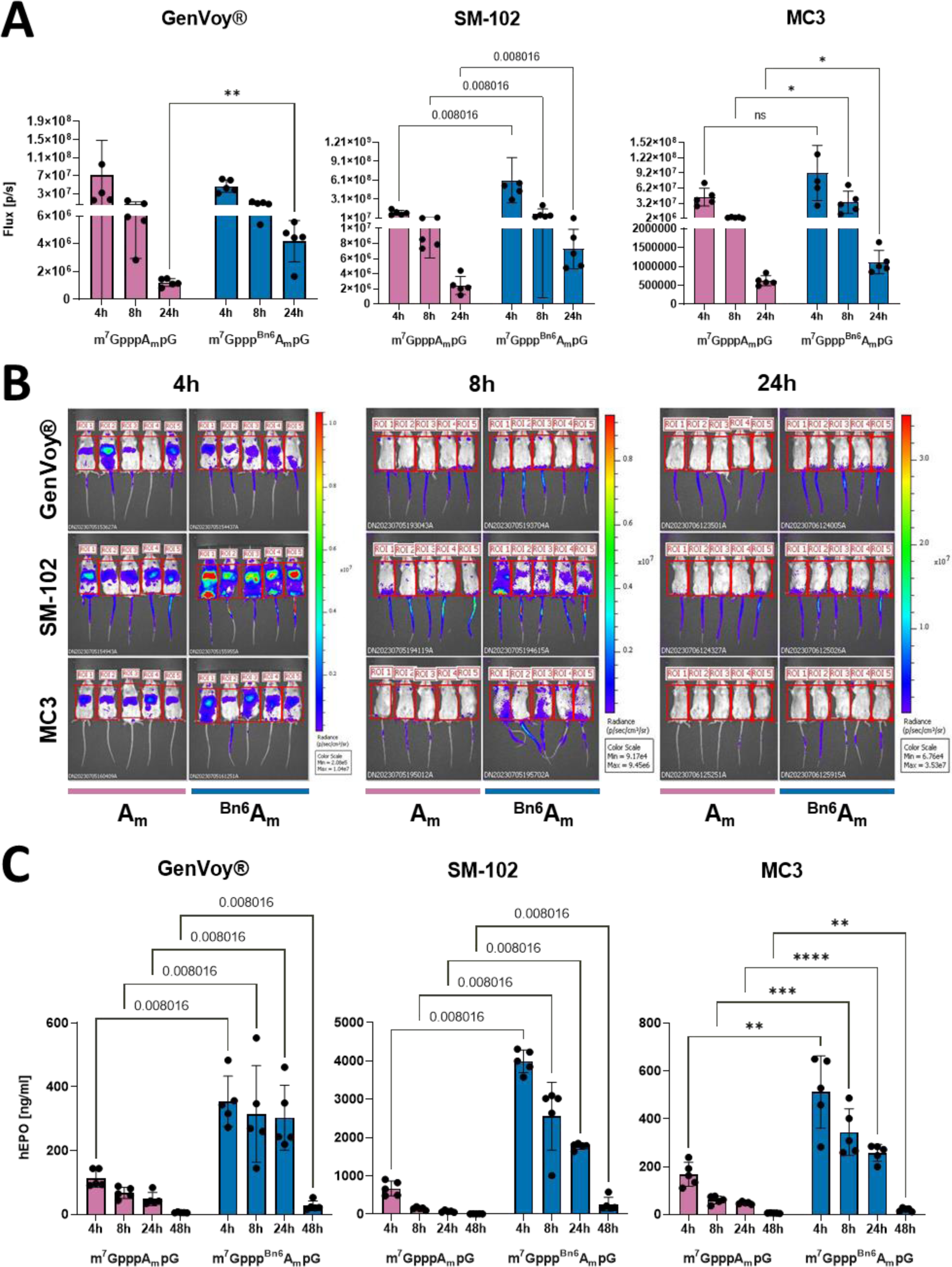
mRNA capped with m^7^Gppp^Bn6^AmpG (*AvantCap*) yields superior protein expression in vivo. (**A**) Intravital bioluminescence at 4 h, 8 h and 24 h in BALB/c mice injected intravenously (i.v.) with firefly luciferase (Fluc) encoding mRNAs, capped with m^7^GpppAmpG or m^7^Gppp^Bn6^AmpG, and formulated using different ionizable lipids (GenVoy-ILM^TM^, SM-102 and MC3). Flux [p/s] mean values ± SD, n=5, GenVoy-ILM^TM^ and MC3 data: * P<0.05, **P<0.01, multiple unpaired *t* test; SM-102 data: Mann-Whitney test, q values are shown. (**B**) Raw bioluminescence images of mice inoculated with Fluc encoding mRNA (scale for bioluminescent signals at the right). (**C**) Human erythropoietin (hEPO) serum concentrations 4 h, 8 h, 24 h and 48 h in C57BL/6 mice injected i.v. with hEPO encoding mRNAs, capped with m^7^GpppAmpG or m^7^Gppp^Bn6^AmpG, and formulated using different ionizable lipids (GenVoy-ILM^TM^, SM-102 and MC3). Data show mean values ± SD, n=5, GenVoy-ILM^TM^ and SM- 102 data: Mann-Whitney test, q values are shown; MC3 data: *P<0.05, **P<0.01, ***P<0.001, ****P<0.0001 multiple unpaired *t* test.

#### m^7^Gppp^Bn6^AmpG-capped mRNA shows superior therapeutic activity in cancer models

To verify if mRNAs capped with m^7^Gppp^Bn6^AmpG can exert therapeutic effects, we have carried out in vivo experiments in mice using two different therapeutically-relevant mRNAs. First, we used mRNAs encoding cancer associated antigens, a strategy used in anticancer vaccinations.^43^ Intravenous administration of mRNA encoding SIINFEKL peptide from ovalbumin (OVA) followed by adoptive transfer of OVA-recognizing OT-I T-cells isolated from the spleens of transgenic C57BL/6- Tg(TcraTcrb)1100Mjb/J mice, resulted in significant expansion of antigen-specific T-cells in recipient animals (Figure S8). Notably, an over 2-fold higher numbers of OT-I T-cells were observed in recipient mice when mRNA was capped with m^7^Gppp^Bn6^AmpG as compared with mice that received mRNA capped with m^7^GpppAmpG (Figure 6A). Encouraged by the results showing that mRNA encoding model antigen can induce expansion of antigen-specific T-cells, we investigated antitumor effects of mRNA encoding tumor antigens in two different tumor models. Lewis lung carcinoma (LLC) cells were stably transduced with OVA and inoculated into C57BL/6 mice. Mice were then treated i.v. with 100 ng of mRNA encoding irrelevant protein (Fluc) or OVA on days 7, 14 and 21. Inhibition of tumor growth was observed only in mice that received OVA-encoding mRNA, and statistically significant (vs controls) inhibition was observed only in mice that received mRNA capped with m^7^Gppp^Bn6^AmpG (Figure 6B). Similarly, significant inhibition of tumor growth was observed in BALB/c mice inoculated with CT26 colon adenocarcinoma cells stably transduced with a model human antigen (NY-ESO1) and treated with mRNA encoding NY-ESO1 and capped with m^7^Gppp^Bn6^AmpG (Figure 6C). Although no formal toxicology studies were performed, we have not noticed any gross adverse effects of mRNA (capped with either m^7^Gppp^Bn6^AmpG or m^7^GpppAmpG) administration such as weight loss, hunched posture, ruffled hair coat, lethargy, anorexia, diarrhea, or neurological impairment.

**Figure 6.**
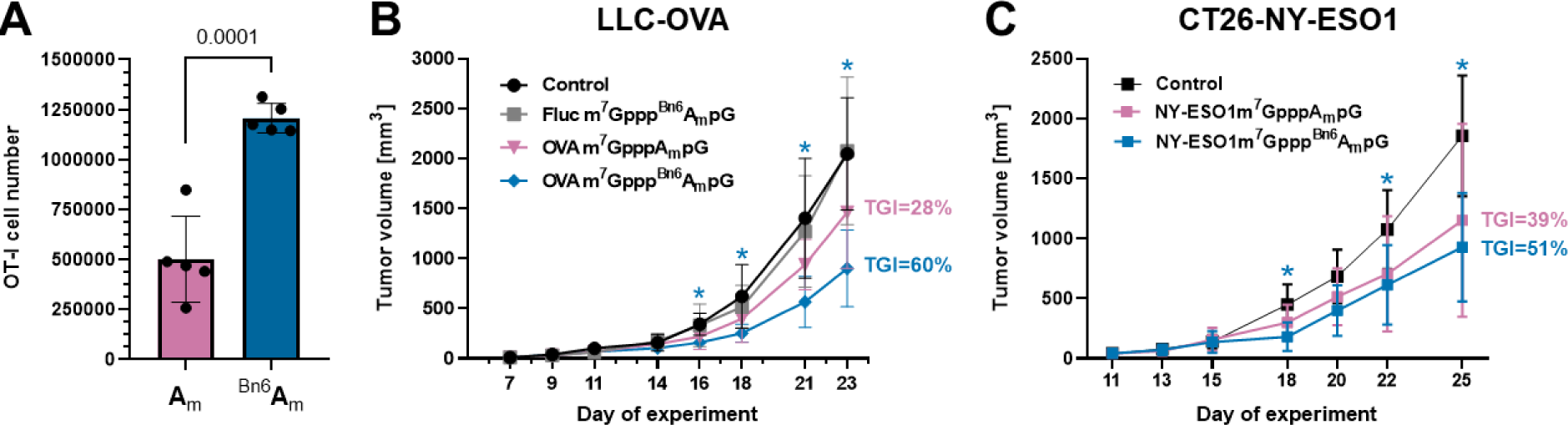
mRNA capped with m^7^Gppp^Bn6^AmpG (*AvantCap*) shows therapeutic activity in cancer models. (**A**) Quantitative OT-I T-cells proliferation in response to in vivo delivery of SIINFEKL antigen/peptide-encoding m^7^GpppAmpG or m^7^Gppp^Bn6^AmpG-capped mRNA. Mice were administered intravenously (i.v.) with 7.5 ng of mRNA in TransIT® formulation. T-cell numbers were normalized to CountBright Absolute Counting Beads (ThermoFisher Scientific). Data show OT-I T-cell cell numbers in the spleen after 72 h proliferation in vivo, data show means±SD, n=5, two-tailed unpaired *t* test. (**B**) C57BL/6 mice were inoculated with Lewis lung carcinoma (LLC) cells stably expressing a model antigen – ovalbumin (OVA) and weekly treated i.v. with 100 ng of OVA-encoding m^7^GpppAmpG or m^7^Gppp^Bn6^AmpG-capped mRNA formulated in TransIT®. Tumor volumes are shown as means±SD, n=8. Mixed-effects analysis with Dunnett’s multiple comparisons test, m^7^Gppp^Bn6^AmpG-capped mRNA vs control: day 16: P=0.0060; day 18: P=0.0410; day 21: P=0.0168; day 23: P=0.0080. Control mice received PBS. m^7^Gppp^Bn6^AmpG- capped mRNA encoding human hEPO served as a tumor antigen-irrelevant control. TGI – tumor growth inhibition compared with controls. (**C**) BALB/c mice were inoculated with CT26 murine adenocarcinoma (CT26) cells stably expressing a human tumor antigen (NY-ESO1) and weekly treated i.v. with 100 ng of NY-ESO1-encoding m^7^GpppAmpG or m^7^Gppp^Bn6^AmpG-capped mRNA formulated in TransIT®. Tumor volumes are shown as means±SD, n=8. 2way ANOVA with Dunnett’s multiple comparisons test, m^7^Gppp^Bn6^AmpG-capped mRNA vs control: day 18: P=0.0079; day 22: P=0.0360; day 25: P=0.0046. Control mice received PBS. TGI – tumor growth inhibition compared with controls.

### Biochemical consequences of incorporating *AvantCap*

To gain a deeper insight into the molecular mechanisms underlying the increased translation yield of mRNAs containing the ^Bn6^Am cap, we performed a series of biochemical and biophysical assays to verify which mRNA-related processes are affected by the modification. We tested the susceptibility of *AvantCap* to dealkylation by FTO and its affinity to several members of eIF4E (eukaryotic translation initiation factor 4E) family proteins. The ^Bn6^Am-capped mRNAs were characterized for translation in a cell-free system and for susceptibility to decapping by Dcp2, which contributes to overall mRNA stability.

#### ^Bn6^Am is not a substrate for FTO

Since the ^Bn6^Am cap was designed as an ^m6^Am cap analog, we first tested its susceptibility to *N*6-dealkylation by FTO – the only known ^m6^Am eraser – by monitoring the reaction progress by RP-HPLC with MS detection (Figure 7). While m^7^Gppp^m6^AmpG was demethylated with a half-life of about 50 minutes, only up to 10% of *AvantCap* was dealkylated even after 20 h of incubation with the enzyme. Also, using a 10-times higher concentration of the m^7^Gppp^Bn6^AmpG, we have not observed any significant dealkylation by FTO (Figure S9). The data suggests that the *AvantCap* can be considered as a stable analog of ^m6^Am cap under physiological conditions.

**Figure 7.**
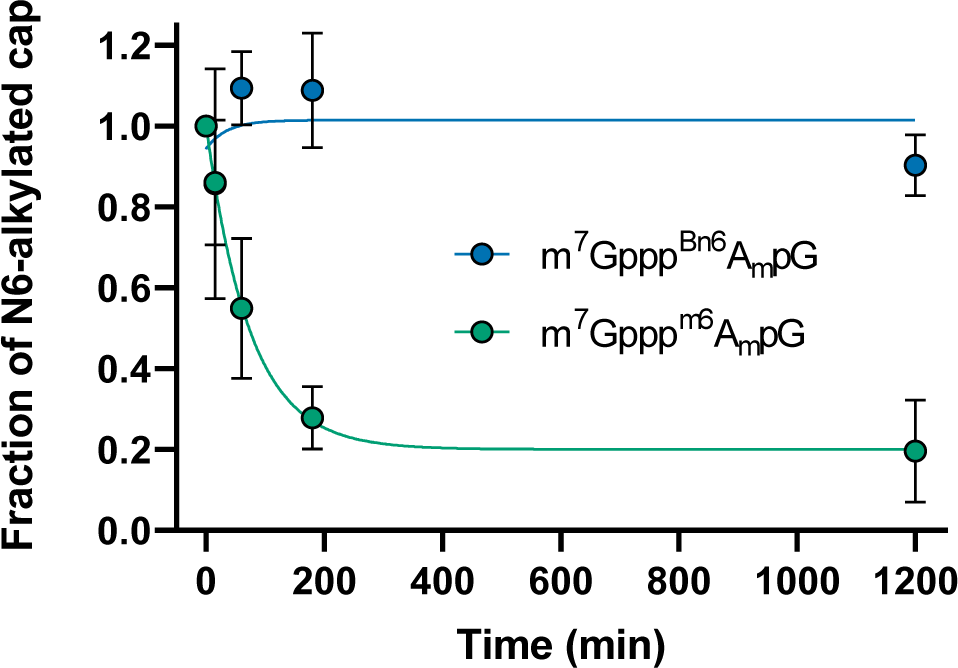
Cap structure containing ^Bn6^Am is resistant to removal by FTO. ^Bn6^Am- or ^m6^Am-containing cap analogs were incubated with FTO and the amount of *N6*-alkylated cap remaining in the mixture was assessed by RP-HPLC-MS. Reaction conditions: 20 µM cap and 2 µM FTO in 50 mM HEPES pH 7, containing 150 mM KCl, 75 µM Fe(II), 300 µM 2-oxoglutarate, 2 mM ascorbic acid. Similar data obtained for 200 µM cap are shown in Figure S9.

#### ^Bn6^Am has minor stabilizing effect on the interaction with translation initiation factor (eIF4E) and does not improve mRNA translation in a cell-free system

The affinity of a cap analog to eIF4E protein is often correlated with the translation efficiency of such capped mRNAs, therefore we quantified the binding interactions between trinucleotide cap structures and eIF4E using time-synchronized fluorescence quenching titration (FQT). To consider potential effects of ^Bn6^Am modification on both translation initiation and its inhibition, we analyzed interactions with three members of the eIF4E family (Table 1, Figure 8A): heIF4E1a, which is a part of the eIF4F complex responsible for ribosome recruitment, h4EHP, and heIF4E3, both of which lack the ability to bind eIF4G and thus act as translation suppressors. We reasoned that the increased translational capacity of mRNAs capped with m^7^Gppp^Bn6^AmpG may be explained by either stabilization of the interaction with eIF4E or destabilization of the interactions with h4EHP or eIF4E3. For all eIF4Es studied, we observed no significant effect for ^m6^Am compared to Am, and for ^Bn6^Am we observed only a minor (1.5–1.75-fold) increase in binding affinity.

**Figure 8.**
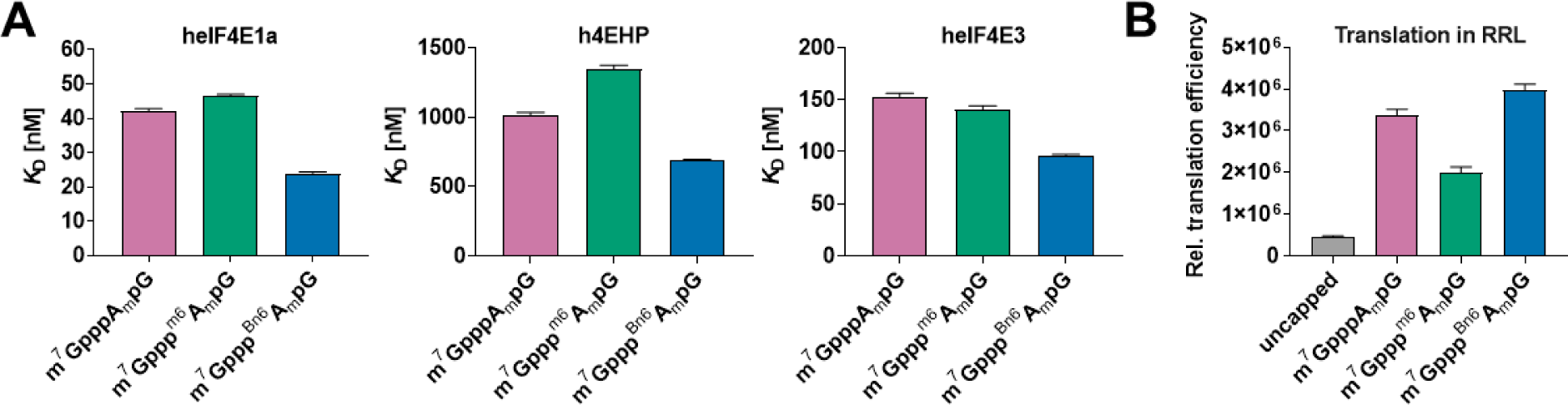
(**A**) Binding affinities of *AvantCap* (**1**) and reference compounds for human translation initiation factor 4E (eIF4E) isoforms 1a and 3 and human 4E homologous protein (4EHP). (**B**) Relative translation efficiencies of mRNA capped with m^7^Gppp^Bn6^AmpG (**1**) and reference analogs in the rabbit reticulocyte lysate (RRL).

**Table 1.** Binding affinities of m^7^Gppp^Bn6^AmpG (1) and reference compounds for human translation initiation factor 4E (eIF4E) isoforms 1a and 3 and human 4E homologous protein (4EHP).

| Cap Analogs | hElF4E1a |  | h4EHP (eIF4E2) |  | hElF4E3 |  |
| --- | --- | --- | --- | --- | --- | --- |
|  | *K<sub>D</sub>* (nM) | *K<sub>D</sub>*/*K<sub>D</sub>* cap-1 | *K<sub>D</sub>* (nM) | *K<sub>D</sub>*/*K<sub>D</sub>* cap-1 | *K<sub>D</sub>* (nM) | *K<sub>D</sub>*/*K<sub>D</sub>* cap-1 |
| m<sup>7</sup>GTP | 7.8 ± 0.3 | 0.19 ± 0.01 | 611 ± 19 | 0.60 ± 0.03 | 129 ± 14 | 0.84 ± 0.11 |
| m<sup>7</sup>GpppA<sub>m</sub>pG | 42.1 ± 0.7 | 1 | 1015 ± 19 | 1 | 153 ± 3 | 1 |
| m<sup>7</sup>Gppp<sup>m6</sup>A<sub>m</sub>pG | 46.4 ± 0.6 | 1.10 ± 0.03 | 1342 ± 32 | 1.32 ± 0.06 | 140 ± 4 | 0.92 ± 0.04 |
| m<sup>7</sup>Gppp<sup>Bn6</sup>A<sub>m</sub>pG | 23.9 ± 0.5 | 0.57 ± 0.02 | 687 ± 10 | 0.68 ± 0.02 | 95.7 ± 1.7 | 0.63 ± 0.02 |

We also compared the translation efficiencies of mRNAs capped with cap-1, ^m6^Am cap, and *AvantCap* in nuclease-treated rabbit reticulocyte lysates (RRL, Figure 8B). Consistent with the previous report,^13^ we observed a 40% reduction in protein production from mRNA with the ^m6^Am cap, which was completely reversed for the *N*6-benzyl modification. Still, this effect is unlikely to account for the up to 6-fold increase in translation yield observed in hDCs and in vivo.

#### AvantCap is susceptibile to decapping and does not affect mRNA stability in cultured cells

Methylation of the *N*6 position of cap-adjacent adenosine has been reported to interfere with decapping by Dcp2,^15^ but our previous in vitro studies suggested that the enzyme is insensitive to either 2′-*O* or *N*6 methylation of adenosine.^21^ We performed an in vitro decapping assay for short RNAs capped with m^7^Gppp^Bn6^AmpG, m^7^Gppp^m6^AmpG, and m^7^GpppAmpG and, surprisingly, found that RNAs carrying ^Bn6^Am modification were slightly more susceptible to decapping compared to both Am and ^m6^Am-capped transcripts (Figure 9). We have also observed no difference in mRNA stability in HEK293T cells electroporated with mRNA capped with m^7^Gppp^Bn6^AmpG, m^7^Gppp^m6^AmpG, or m^7^GpppAmpG as evidenced by RT-qPCR (Figure S10).

**Figure 9.**
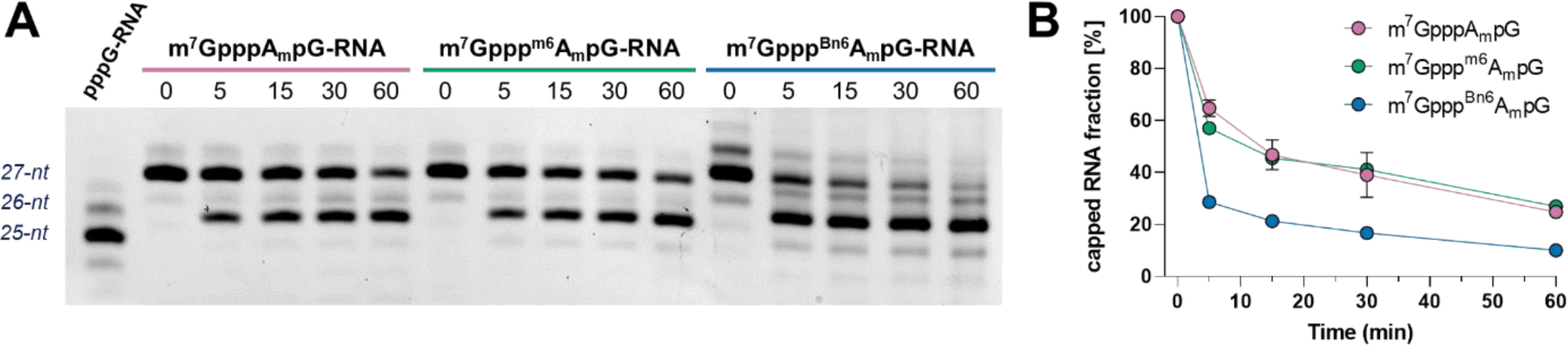
Susceptibility of short RNAs to decapping by PNRC2-hDcp1/Dcp2 complex in vitro. Short 27-nt capped RNAs (20 ng) were subjected to PNRC2-hDcp1/Dcp2 (13 nM) complex for 60 min at 37°C. Aliquots from different time points were resolved by polyacrylamide gel electrophoresis (PAGE), stained with SYBR Gold and analyzed by densitometry. (**A**) Representative PAGE gel from single experiment; (**B**) Results from triplicate experiments ± SEM. Error bars are not visible if smaller than data points. Individual data for all replicates is shown in Figure S11.

#### Protein fraction pulled-down from HEK293F cell extract using immobilized AvantCap is enriched in eIF4E and eIF3

Since none of the biochemical assays provided a satisfactory explanation for the mechanism underlying the increased translation of mRNAs capped with *AvantCap* in several cell lines and in vivo, we searched for the potential selective interactors of ^m6^Am and ^Bn6^Am caps in cell lysate using a pull-down assay (Figure 10A). To this end, we synthesized a series of trinucleotide cap analogs **2a-c** functionalized with an amine-terminated linker at the guanosine ribose and immobilized them on BrCN-activated Sepharose (Figure S12A). The resulting affinity resins **AR-1** (Am), **AR-2** (^m6^Am), and **AR-3** (^Bn6^Am; Figure 10B) were incubated with HEK293F cell extract in the presence of GTP to limit non-specific interactions. The pulled-down proteins were eluted with the corresponding trinucleotide cap analog (m^7^GpppAmpG for **AR-1**, m^7^Gppp^m6^AmpG for **AR-2,** and m^7^Gppp^Bn6^AmpG for **AR-3**), digested with trypsin, labeled with isobaric tags (TMT), and analyzed by shotgun proteomics. The list of identified proteins and the results of Student’s t-tests (two-sided, unpaired) performed to assess binding preferences of proteins to the resins are given in Table S3. Based on the statistical method applied, 29 protein groups were classified as preferentially binding to **AR-2** over **AR-1** and 36 as preferentially binding to **AR-3** over **AR-1** (Figure 10D); 17 of these were enriched in both **AR-2** and **AR-3** over **AR-1** (Figure 10C). Notably, the common group of proteins contained eIF4E, DcpS and eIF3 (11 of 13 subunits; the remaining two – eIF3J and eIF3K – were also enriched but did not pass our stringent statistical significance test) which were rather abundant in these eluates, as estimated by their iBAQ values (Figure S12B).^44^ The remaining four are present at lower levels and include RPA1, DNAJC10, R3HCC1L, and UPP1, none of which are directly related to mRNA translation. We also found 7 proteins whose concentrations were significantly reduced in both **AR-2** and **AR-3** eluates as compared to **AR-1**. These included NCBP1, which is known to stabilize the interaction of cap with NCBP2, and several other nuclear RNA-binding proteins: FUS, TIA1, TAF15, EWSR1, HNRNPH3, and HNRNPA2B1. Consistent with a previous report on snRNA,^14 m6^Am (and ^Bn6^Am but to a lesser extent) appeared to destabilize the interactions of cap with NCBP2. We observed no differences in the abundance of FTO among the samples, although it was generally present at low levels (Figure S12B).

**Figure 10.**
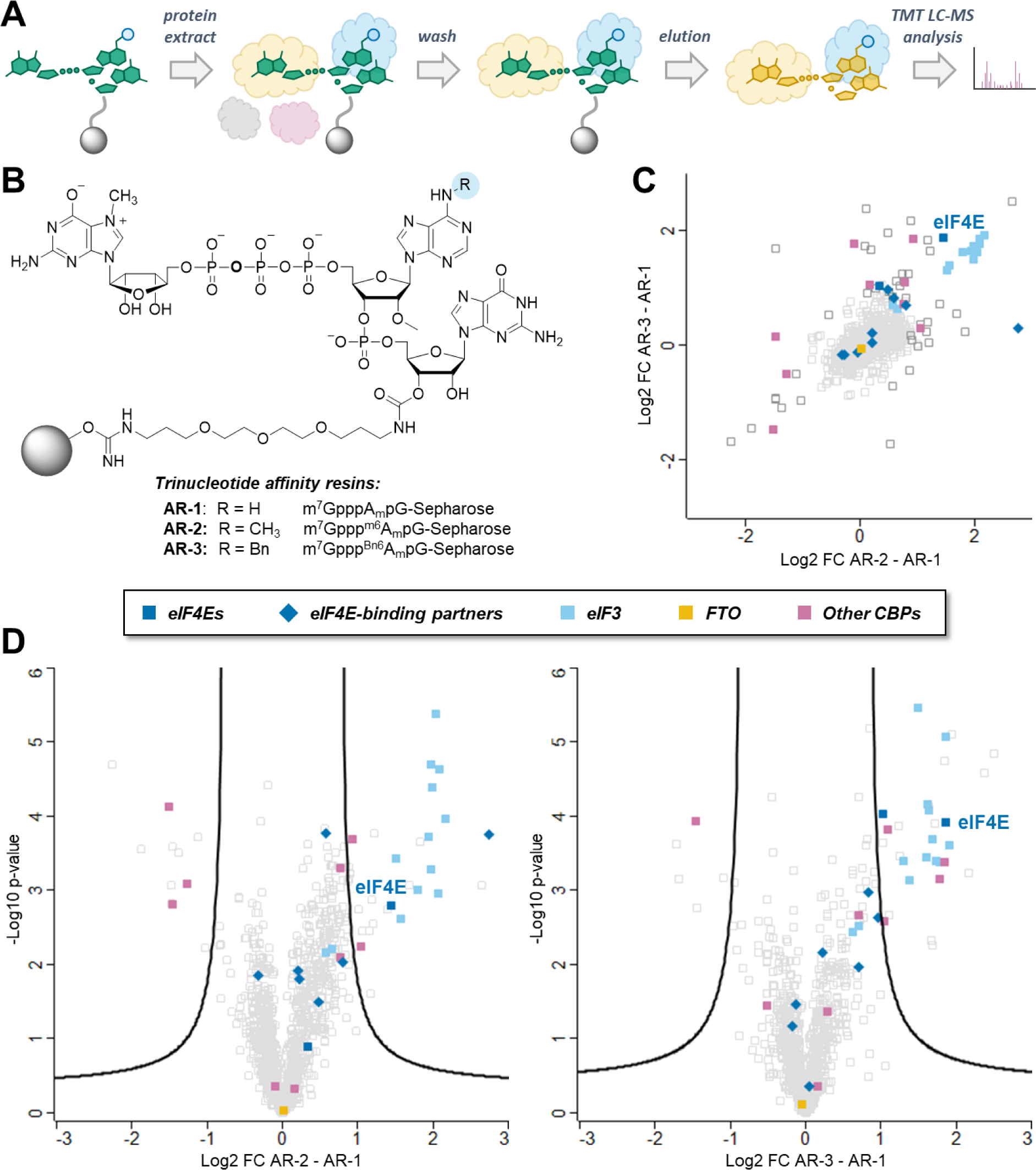
Pull-down assay with protein extract from HEK293F cells. (**A**) The workflow of the experiment. (**B**) Chemical structure of trinucleotide affinity resins. (**C**) Dot plot representations of correlation between log2 fold change **AR-3**/**AR-1** and log2 fold change **AR-2**/**AR-1**. (**D**) Volcano plots displaying the log2 fold change (log2 FC) against the t-test-derived −log10 statistical p-value (-Log p-value) for the protein groups in the eluates from trinucleotide cap affinity resins **AR-2**/**AR-1** (n = 3) and **AR-2**/**AR-1** (n = 3). Student’s t-tests (two-sided, unpaired) were performed to assess binding preferences of protein groups to **AR-2** and **AR-3** resins relative to **AR-1** resin.

One of the cap-dependent pathways of translation initiation in human cells relies on cap-binding activity of eIF3d, a subunit of the 800-kilodalton eIF3 complex. Based on the results of pull-down assay, we hypothesized that the increased mRNA translation might be caused by preferential engagement of alternative cap-dependent translation initiation that relies on cap-binding activity of eIF3d.^45^ To verify this hypothesis we pre-incubated hDCs with an mTOR inhibitor, INK128, which leads to dephosphorylation of the 4E-BP1 allowing its binding to and inhibiting eIF4E thereby creating conditions for alternative cap-dependent translation initiation.^46-48^ Transfection of hDCs with mRNA encoding hEPO led to higher protein concentrations, when mRNA was capped with m^7^Gppp^Bn6^AmpG (Figure S13A left). Preincubation with INK128 strongly attenuated mRNA translation for all tested mRNAs, but this effect was more pronounced, when mRNA was capped with m^7^Gppp^m6^AmpG or m^7^GpppAmpG as compared with m^7^Gppp^Bn6^AmpG (Figure S13A right). eIF3d makes specific contacts with the cap and these interactions are essential for the assembly of translation initiation complexes on eIF3-specialized mRNAs such as the one for the cell proliferation regulator c-JUN.^46^ c-JUN mRNA contains an inhibitory RNA element that blocks eIF4E recruitment, thus enforcing alternative cap recognition by eIF3d. We have prepared mRNAs encoding Fluc with variants of c-JUN 5’UTRs: (i) C-JUN – 300 nt of the wild-type c-JUN, and (ii) ΔeIF3 – C-JUN 5’UTR without stem loop responsible for eIF3 binding (nucleotide position 181–214) (Figure S13B). Transfection of these mRNA constructs into hDCs revealed that while the luminescence with normal 5’-UTRs (human beta globin) is 20% higher when mRNA is capped with m^7^Gppp^Bn6^AmpG (as compared with mRNA capped with m^7^Gppp^m6^AmpG), the difference is almost 4-fold higher, when C-JUN is used as 5’-UTR (Figure S13C). Deletion of stem loop responsible for eIF3 binding abrogates translational advantage of mRNA capped with m^7^Gppp^Bn6^AmpG (Figure S13C). These results suggest that the advantage of using m^7^Gppp^Bn6^AmpG may be at least to some extent due to the eIF3-depentent translation initiation and interaction of eIF3d with *AvantCap*, albeit this requires further experimental verification.

## DISCUSSION

The success of mRNA-based therapeutic approaches has been facilitated in large part by the use of chemical modifications to manipulate biological properties. The prime example is the substitution of uridines with (1-methyl)pseudouridines in the RNA body to reduce the undesired reactogenicity of mRNA,^49^ but some chemical modifications of mRNA ends have also proven to be beneficial in this context.^32^ Here, we report *AvantCap* – a novel synthetically easily accessible trinucleotide cap analog containing an *N*6-benzyl-2’-*O*-methyladenylate moiety (Bn^6^Am) that confers superior properties to in vitro transcribed capped mRNAs with potential benefits for mRNA therapeutics (Figure 1, Scheme 1). The design of this analog was inspired by the naturally occurring 5’-terminal ^m6^Am mark present in many endogenous mammalian mRNAs of yet not fully understood function. Here, by replacing the *N6*-methyl in m^7^Gppp^m6^Am with a benzyl group, we obtained a mimic that is more hydrophobic, resistant to demethylation by FTO, efficiently incorporated into RNA 5’ end, and yields mRNAs that are more translationally active in many of the tested biological settings (Figures 2–7). The presence of the ^Bn6^Am modification results in a hydrophobic effect,^41^ which is useful for RNA purification on RP-HPLC columns as it significantly increases RNA retention (Figures 2, 3). For RNAs up to 2000 nt, this effect is strong enough to allow separation of capped and uncapped RNAs, even on a preparative scale. We speculate that even for longer RNAs, for which no clear separation is observed, enrichment of capped RNA species is possible. In addition, mRNAs produced in the presence of *AvantCap* contains less dsRNA impurities than mRNAs obtained in the presence of m^7^GpppAmpG (Figure S2). The purity of mRNAs, including dsRNA and uncapped RNA content, has recently been identified as one of the most important factors for their therapeutic applications, particularly in anti-cancer and protein replacement approaches.^50^ Therefore, we believe that *AvantCap* can provide a significant improvement in this area.

We investigated the biological properties of mRNAs capped with *AvantCap* in various biological settings as recent studies have revealed that translational properties of IVT transcribed mRNAs may be cell-type dependent. Indeed, we have found that in several mammalian cell lines such as HEK293 or A549 mRNAs carrying *AvantCap* have comparable properties to mRNAs carrying unmodified cap 1 (Figure 4). However, in other cell types such as CT26, macrophages or dendritic cells mRNAs carrying the ^Bn6^Am mark afford significantly higher protein outputs. This effect is also observed in vivo, for all reporter proteins tested, albeit its magnitude is varying depending on the mRNA and formulation type (Figure 5). Among three different LNP-types, the formulation based on lipid SM-102, provided best differentiation between differently capped mRNAs and showed up to 6-fold higher expression of mRNAs carrying *AvantCap*, compared to mRNAs carrying cap 1 structure. Even more pronounced superior activity of mRNAs carrying *AvantCap* was observed for cationic-lipid transfection reagent (TransIT®) (Figure S6). It is unclear why different lipid-formulations result in such differences in biological activity of mRNA, but it may be related to different cellular signaling pathways being activated by different delivery methods and/or different cell-types being targeted in vivo. We also show that mRNAs carrying *AvantCap* show superior activity in therapeutic models of anti-cancer vaccines (Figure 6, Figure S8)

Trying to uncover the rationale behind the biological effects of *AvantCap*, we thoroughly characterized its biochemical properties in vitro. To that end, we analyzed binding affinity to eIF4E and translational properties in a cell-free system (Table 1, Figure 8), susceptibility to enzymatic decapping (Figure 9), and dealkylation by FTO (Figure 7). These data indicate that m^7^Gppp^Bn6^AmpG is an FTO-, but not Dcp2- resistant mimic of m^7^Gppp^m6^AmpG, with slightly increased affinity to eIF4E. To further compare biochemical properties of m^7^Gppp^Bn6^AmpG, m^7^Gppp^m6^AmpG, and m^7^GpppAmpG we synthesized a series of trinucleotide affinity resins and performed a pull-down assay with cellular protein extracts (Figure 10). Two possible mechanisms emerge from this assay: (i) direct involvement of eIF3 in the differentiation between Am and ^m6^Am/^Bn6^Am caps or (ii) enhancement of selectivity towards eIF4E in a complex cellular environment upon *N*6 modification of the cap, by reducing its affinity for off-target proteins. The first hypothesis seems attractive in the context of recent reports on alternative cap-dependent translation initiation using the eIF4G2 (DAP5)/eIF3d pathway activated under stress conditions.^48^ Cell culture experiments under conditions prohibitory to conventional translation pathway, did not exclude the possibility of participation of *AvantCapped*-mRNAs in alternative translation mechanisms, but it requires further investigation. The second possibility is analogous to the mechanism of evading the innate immune response by forming the cap-1 structure, which is not recognized by IFIT proteins or RIG-I-like receptors. It is also possible that the observed differences in biological activity arise from subtly different impurity profiles for differently capped mRNAs that differentially activate cellular signaling pathways and the innate immune system. However, we have put significant effort into preparing high quality mRNA by employing two step purification process, including RP-HPLC purification, and verifying quality of the final mRNA products (Table S2, Figures S3,S4). Overall, the superior biological activity of *AvantCap* combined with straightforward and scalable synthesis, make this analog an attractive opportunity for advanced applications of IVT mRNA.

## Supporting Information

All experimental details, supporting Tables S1-S3, Figures S1-S13, NMR and HRMS Spectra for chemical compounds.

## Supporting information

S1-Experimental section

Graphical summary

## Acknowledgments

We thank Pawel Turowski (Explorna) for technical assistance in cell culture experiments. Financial support from the National Science Centre (2018/31/B/ST5/03821 to J.K, 2019/33/B/ST4/01843 to J.J.) is gratefully acknowledged. The research was partially carried out in the frames of the project co-financed by the European Union from the European Regional Development Fund under the Smart Growth Operational Program. Project implemented as part of the National Centre for Research and Development call: Fast Track 6/1.1.1/2019

## Conflict of interest

J.J., J.K., P.J.S., and M.W. are inventors of a patent related to *AvantCap*. Some of the authors are shareholders of Explorna Therapuetics.

## References

1. Furuichi, Y.; Muthukrishnan, S.; Shatkin, A. J. 5’-Terminal m-7G(5’)ppp(5’)G-m-p in vivo: identification in reovirus genome RNA. Proceedings of the National Academy of Sciences 1975, 72 (2), 742-745. Bélanger, F.; Stepinski, J.; Darzynkiewicz, E.; Pelletier, J. Characterization of hMTr1, a Human Cap1 2′-O-Ribose Methyltransferase*. Journal of Biological Chemistry 2010, 285 (43), 33037-33044. DOI: 10.1074/jbc.M110.155283. Langberg, S. R.; Moss, B. Post-transcriptional modifications of mRNA. Purification and characterization of cap I and cap II RNA (nucleoside-2’-)-methyltransferases from HeLa cells. Journal of Biological Chemistry 1981, 256 (19), 10054–10060. DOI: 10.1016/S0021-9258(19)68740-5.

2. Drazkowska, K.; Tomecki, R.; Warminski, M.; Baran, N.; Cysewski, D.; Depaix, A.; Kasprzyk, R.; Kowalska, J.; Jemielity, J.; Sikorski, Pawel J. 2′-O-Methylation of the second transcribed nucleotide within the mRNA 5′ cap impacts the protein production level in a cell-specific manner and contributes to RNA immune evasion. Nucleic Acids Research 2022, 50 (16), 9051–9071. DOI: 10.1093/nar/gkac722 (acccessed 11/17/2022).

3. Werner, M.; Purta, E.; Kaminska, K. H.; Cymerman, I. A.; Campbell, D. A.; Mittra, B.; Zamudio, J. R.; Sturm, N. R.; Jaworski, J.; Bujnicki, J. M. 2′-O-ribose methylation of cap2 in human: function and evolution in a horizontally mobile family. Nucleic Acids Research 2011, 39 (11), 4756–4768. DOI: 10.1093/nar/gkr038. Despic, V.; Jaffrey, S. R. mRNA ageing shapes the Cap2 methylome in mammalian mRNA. Nature 2023, 614 (7947), 358-366. DOI: 10.1038/s41586-022-05668-z. Smietanski, M.; Werner, M.; Purta, E.; Kaminska, K. H.; Stepinski, J.; Darzynkiewicz, E.; Nowotny, M.; Bujnicki, J. M. Structural analysis of human 2′-O-ribose methyltransferases involved in mRNA cap structure formation. Nature Communications 2014, 5, 3004, Article. DOI: 10.1038/ncomms4004 https://www.nature.com/articles/ncomms4004#supplementary-information.

4. Sahin, U.; Muik, A.; Derhovanessian, E.; Vogler, I.; Kranz, L. M.; Vormehr, M.; Baum, A.; Pascal, K.; Quandt, J.; Maurus, D.;, et al. COVID-19 vaccine BNT162b1 elicits human antibody and TH1 T cell responses. Nature 2020, 586 (7830), 594–599. DOI: 10.1038/s41586-020-2814-7. Corbett, K. S.; Edwards, D. K.; Leist, S. R.; Abiona, O. M.; Boyoglu-Barnum, S.; Gillespie, R. A.; Himansu, S.; Schäfer, A.; Ziwawo, C. T.; DiPiazza, A. T.;, et al. SARS-CoV-2 mRNA vaccine design enabled by prototype pathogen preparedness. Nature 2020, *586* (7830), 567-571. DOI: 10.1038/s41586-020-2622-0.

5. Wei, C.-M.; Gershowitz, A.; Moss, B. N6, O2′-dimethyladenosine a novel methylated ribonucleoside next to the 5′ terminal of animal cell and virus mRNAs. Nature 1975, 257, 251. DOI: 10.1038/257251a0. Keith, J. M.; Ensinger, M. J.; Moss, B. HeLa cell RNA (2‘-O-methyladenosine-N6-)-methyltransferase specific for the capped 5‘-end of messenger RNA. Journal of Biological Chemistry 1978, 253 (14), 5033-5039. DOI: 10.1016/S0021-9258(17)34652-5.

6. Meyer, K. D.; Jaffrey, S. R. Rethinking m6A Readers, Writers, and Erasers. Annual Review of Cell and Developmental Biology 2017, 33 (1), 319–342. DOI: 10.1146/annurev-cellbio-100616-060758 (acccessed 2023/02/22). Shi, H.; Wei, J.; He, C. Where, When, and How: Context-Dependent Functions of RNA Methylation Writers, Readers, and Erasers. Molecular Cell 2019, 74 (4), 640-650. DOI: 10.1016/j.molcel.2019.04.025.

7. Dominissini, D.; Moshitch-Moshkovitz, S.; Schwartz, S.; Salmon-Divon, M.; Ungar, L.; Osenberg, S.; Cesarkas, K.; Jacob-Hirsch, J.; Amariglio, N.; Kupiec, M.;, et al. Topology of the human and mouse m6A RNA methylomes revealed by m6A-seq. Nature 2012, 485 (7397), 201–206. DOI: 10.1038/nature11112. Meyer, Kate D.; Saletore, Y.; Zumbo, P.; Elemento, O.; Mason, Christopher E.; Jaffrey, Samie R. Comprehensive Analysis of mRNA Methylation Reveals Enrichment in 3′ UTRs and near Stop Codons. Cell 2012, 149 (7), 1635-1646. DOI: 10.1016/j.cell.2012.05.003.

8. Anreiter, I.; Tian, Y. W.; Soller, M. The cap epitranscriptome: Early directions to a complex life as mRNA. BioEssays 2023, 45 (3), 2200198, 10.1002/bies.202200198. DOI: 10.1002/bies.202200198 (acccessed 2023/02/22).

9. Cesaro, B.; Tarullo, M.; Fatica, A. Regulation of Gene Expression by m6Am RNA Modification. In International Journal of Molecular Sciences, 2023; Vol. 24.

10. Akichika, S.; Hirano, S.; Shichino, Y.; Suzuki, T.; Nishimasu, H.; Ishitani, R.; Sugita, A.; Hirose, Y.; Iwasaki, S.; Nureki, O.;, et al. Cap-specific terminal N6-methylation of RNA by an RNA polymerase II–associated methyltransferase. Science 2019, 363 (6423), eaav0080. DOI: 10.1126/science.aav0080.

11. Boulias, K.; Toczydłowska-Socha, D.; Hawley, B. R.; Liberman, N.; Takashima, K.; Zaccara, S.; Guez, T.; Vasseur, J.-J.; Debart, F.; Aravind, L.;, et al. Identification of the m6Am Methyltransferase PCIF1 Reveals the Location and Functions of m6Am in the Transcriptome. Molecular Cell 2019, 75 (3), 631–643.e638. DOI: 10.1016/j.molcel.2019.06.006.

12. Sun, H.; Zhang, M.; Li, K.; Bai, D.; Yi, C. Cap-specific, terminal N6-methylation by a mammalian m6Am methyltransferase. Cell Research 2019, 29 (1), 80–82. DOI: 10.1038/s41422-018-0117-4.

13. Sendinc, E.; Valle-Garcia, D.; Dhall, A.; Chen, H.; Henriques, T.; Navarrete-Perea, J.; Sheng, W.; Gygi, S. P.; Adelman, K.; Shi, Y. PCIF1 Catalyzes m6Am mRNA Methylation to Regulate Gene Expression. Molecular Cell 2019, 75 (3), 620–630.e629. DOI: 10.1016/j.molcel.2019.05.030.

14. Koh, C. W. Q.; Goh, Y. T.; Goh, W. S. S. Atlas of quantitative single-base-resolution N6-methyl-adenine methylomes. Nature Communications 2019, 10 (1), 5636. DOI: 10.1038/s41467-019-13561-z.

15. Mauer, J.; Luo, X.; Blanjoie, A.; Jiao, X.; Grozhik, A. V.; Patil, D. P.; Linder, B.; Pickering, B. F.; Vasseur, J.-J.; Chen, Q.;, et al. Reversible methylation of m6Am in the 5′ cap controls mRNA stability. Nature 2017, 541 (7637), 371–375, Article. DOI: 10.1038/nature21022 http://www.nature.com/nature/journal/v541/n7637/abs/nature21022.html#supplementary-information.

16. Wei, J.; Liu, F.; Lu, Z.; Fei, Q.; Ai, Y.; He, P. C.; Shi, H.; Cui, X.; Su, R.; Klungland, A.;, et al. Differential m6A, m6Am, and m1A Demethylation Mediated by FTO in the Cell Nucleus and Cytoplasm. Molecular Cell 2018, 71 (6), 973-985.e975. DOI: 10.1016/j.molcel.2018.08.011.

17. Kruse, S.; Zhong, S.; Bodi, Z.; Button, J.; Alcocer, M. J. C.; Hayes, C. J.; Fray, R. A novel synthesis and detection method for cap-associated adenosine modifications in mouse mRNA. Scientific Reports 2011, 1, 126, Article. DOI: 10.1038/srep00126 https://www.nature.com/articles/srep00126#supplementary-information.

18. Galloway, A.; Atrih, A.; Grzela, R.; Darzynkiewicz, E.; Ferguson, M. A. J.; Cowling, V. H. CAP-MAP: cap analysis protocol with minimal analyte processing, a rapid and sensitive approach to analysing mRNA cap structures. Open Biology 2020, 10 (2), 190306. DOI: 10.1098/rsob.190306 (acccessed 2023/02/22). Wang, J.; Alvin Chew, B. L.; Lai, Y.; Dong, H.; Xu, L.; Balamkundu, S.; Cai, W. M.; Cui, L.; Liu, C. F.; Fu, X.-Y.;, et al. Quantifying the RNA cap epitranscriptome reveals novel caps in cellular and viral RNA. Nucleic Acids Research 2019, 47 (20), e130-e130. DOI: 10.1093/nar/gkz751 (acccessed 2/23/2023).

19. Muthmann, N.; Albers, M.; Rentmeister, A. CAPturAM, a Chemo-Enzymatic Strategy for Selective Enrichment and Detection of Physiological CAPAM-Targets. Angewandte Chemie International Edition 2023, 62 (4), e202211957, 10.1002/anie.202211957. DOI: 10.1002/anie.202211957 (acccessed 2023/02/22).

20. Sun, H.; Li, K.; Zhang, X.; Liu, J. e.; Zhang, M.; Meng, H.; Yi, C. m6Am-seq reveals the dynamic m6Am methylation in the human transcriptome. Nature Communications 2021, 12 (1), 4778. DOI: 10.1038/s41467-021-25105-5.

21. Sikorski, P. J.; Warminski, M.; Kubacka, D.; Ratajczak, T.; Nowis, D.; Kowalska, J.; Jemielity, J. The identity and methylation status of the first transcribed nucleotide in eukaryotic mRNA 5′ cap modulates protein expression in living cells. Nucleic Acids Research 2020, 48 (4), 1607–1626. DOI: 10.1093/nar/gkaa032 (acccessed 1/5/2021).

22. Ben-Haim, M. S.; Pinto, Y.; Moshitch-Moshkovitz, S.; Hershkovitz, V.; Kol, N.; Diamant-Levi, T.; Beeri, M. S.; Amariglio, N.; Cohen, H. Y.; Rechavi, G. Dynamic regulation of N6,2′-O-dimethyladenosine (m6Am) in obesity. Nature Communications 2021, 12 (1), 7185. DOI: 10.1038/s41467-021-27421-2.

23. Pandey, R. R.; Delfino, E.; Homolka, D.; Roithova, A.; Chen, K.-M.; Li, L.; Franco, G.; Vågbø, C. B.; Taillebourg, E.; Fauvarque, M.-O.;, et al. The Mammalian Cap-Specific m6Am RNA Methyltransferase PCIF1 Regulates Transcript Levels in Mouse Tissues. Cell Reports 2020, 32 (7), 108038. DOI: 10.1016/j.celrep.2020.108038.

24. Tartell, M. A.; Boulias, K.; Hoffmann, G. B.; Bloyet, L.-M.; Greer, E. L.; Whelan, S. P. J. Methylation of viral mRNA cap structures by PCIF1 attenuates the antiviral activity of interferon-β. Proceedings of the National Academy of Sciences 2021, 118 (29), e2025769118. DOI: 10.1073/pnas.2025769118.

25. Zhang, Q.; Kang, Y.; Wang, S.; Gonzalez, G. M.; Li, W.; Hui, H.; Wang, Y.; Rana, T. M. HIV reprograms host m6Am RNA methylome by viral Vpr protein-mediated degradation of PCIF1. Nature Communications 2021, 12 (1), 5543. DOI: 10.1038/s41467-021-25683-4.

26. Wang, L.; Wang, S.; Wu, L.; Li, W.; Bray, W.; Clark, A. E.; Gonzalez, G. M.; Wang, Y.; Carlin, A. F.; Rana, T. M. PCIF1-mediated deposition of 5′-cap N6,2′-O-dimethyladenosine in ACE2 and TMPRSS2 mRNA regulates susceptibility to SARS-CoV-2 infection. Proceedings of the National Academy of Sciences 2023, 120 (5), e2210361120. DOI: 10.1073/pnas.2210361120 (acccessed 2023/02/23).

27. Wang, L.; Wu, L.; Zhu, Z.; Zhang, Q.; Li, W.; Gonzalez, G. M.; Wang, Y.; Rana, T. M. Role of PCIF1-mediated 5′-cap N6-methyladeonsine mRNA methylation in colorectal cancer and anti-PD-1 immunotherapy. The EMBO Journal 2023, 42 (2), e111673, 10.15252/embj.2022111673. DOI: 10.15252/embj.2022111673 (acccessed 2023/02/23).

28. van Dülmen, M.; Muthmann, N.; Rentmeister, A. Chemo-Enzymatic Modification of the 5′ Cap Maintains Translation and Increases Immunogenic Properties of mRNA. Angewandte Chemie International Edition 2021, 60 (24), 13280–13286, 10.1002/anie.202100352. DOI: 10.1002/anie.202100352 (acccessed 2021/07/01).

29. Ishikawa, M.; Murai, R.; Hagiwara, H.; Hoshino, T.; Suyama, K. Preparation of eukaryotic mRNA having differently methylated adenosine at the 5′-terminus and the effect of the methyl group in translation. Nucleic Acids Symposium Series 2009, 53 (1), 129–130. DOI: 10.1093/nass/nrp065 (acccessed 2/24/2023).

30. Sahin, U.; Karikó, K.; Türeci, Ö. mRNA-based therapeutics — developing a new class of drugs. Nature Reviews Drug Discovery 2014, 13 (10), 759–780. DOI: 10.1038/nrd4278.

31. Warminski, M.; Sikorski, P. J.; Kowalska, J.; Jemielity, J. Applications of Phosphate Modification and Labeling to Study (m)RNA Caps. Topics in Current Chemistry 2017, 375 (1), 16. DOI: 10.1007/s41061-017-0106-y. Bollu, A.; Peters, A.; Rentmeister, A. Chemo-Enzymatic Modification of the 5′ Cap To Study mRNAs. Accounts of Chemical Research 2022, 55 (9), 1249-1261. DOI: 10.1021/acs.accounts.2c00059.

32. Warminski, M.; Mamot, A.; Depaix, A.; Kowalska, J.; Jemielity, J. Chemical Modifications of mRNA Ends for Therapeutic Applications. Accounts of Chemical Research 2023, 56 (20), 2814–2826. DOI: 10.1021/acs.accounts.3c00442.

33. Pasquinelli, A. E.; Dahlberg, J. E.; Lund, E. Reverse 5’ caps in RNAs made in vitro by phage RNA polymerases. RNA 1995, 1 (9), 957–967.

34. Henderson, J. M.; Ujita, A.; Hill, E.; Yousif-Rosales, S.; Smith, C.; Ko, N.; McReynolds, T.; Cabral, C. R.; Escamilla-Powers, J. R.; Houston, M. E. Cap 1 Messenger RNA Synthesis with Co-transcriptional CleanCap® Analog by In Vitro Transcription. Current Protocols 2021, 1 (2), e39, 10.1002/cpz1.39. DOI: 10.1002/cpz1.39 (acccessed 2021/07/02). Senthilvelan, A.; Vonderfecht, T.; Shanmugasundaram, M.; Potter, J.; Kore, A. R. Click-iT trinucleotide cap analog: Synthesis, mRNA translation, and detection. Bioorganic & Medicinal Chemistry 2023, 77, 117128. DOI: 10.1016/j.bmc.2022.117128. Depaix, A.; Grudzien-Nogalska, E.; Fedorczyk, B.; Kiledjian, M.; Jemielity, J.; Kowalska, J. Preparation of RNAs with non-canonical 5′ ends using novel di- and trinucleotide reagents for co-transcriptional capping. Frontiers in Molecular Biosciences 2022, 9, Original Research. Kozarski, M.; Drazkowska, K.; Bednarczyk, M.; Warminski, M.; Jemielity, J.; Kowalska, J. Towards superior mRNA caps accessible by click chemistry: synthesis and translational properties of triazole-bearing oligonucleotide cap analogs. RSC Advances 2023, 13 (19), 12809-12824, 10.1039/D3RA00026E. DOI: 10.1039/D3RA00026E.

35. Brown, C. J.; McNae, I.; Fischer, P. M.; Walkinshaw, M. D. Crystallographic and Mass Spectrometric Characterisation of eIF4E with N7-alkylated Cap Derivatives. Journal of Molecular Biology 2007, 372 (1), 7–15. DOI: 10.1016/j.jmb.2007.06.033. Grudzien, E. W. A.; Stepinski, J.; Jankowska-Anyszka, M.; Stolarski, R.; Darzynkiewicz, E.; Rhoads, R. E. Novel cap analogs for in vitro synthesis of mRNAs with high translational efficiency. RNA 2004, 10 (9), 1479-1487. Jia, Y.; Chiu, T.-L.; Amin, E. A.; Polunovsky, V.; Bitterman, P. B.; Wagner, C. R. Design, synthesis and evaluation of analogs of initiation factor 4E (eIF4E) cap-binding antagonist Bn7-GMP. European Journal of Medicinal Chemistry 2010, 45 (4), 1304–1313. DOI: 10.1016/j.ejmech.2009.11.054. Wojcik, R.; Baranowski, M. R.; Markiewicz, L.; Kubacka, D.; Bednarczyk, M.; Baran, N.; Wojtczak, A.; Sikorski, P. J.; Zuberek, J.; Kowalska, J.;, et al. Novel N7-Arylmethyl Substituted Dinucleotide mRNA 5′ cap Analogs: Synthesis and Evaluation as Modulators of Translation. In Pharmaceutics, 2021; Vol. 13. Grzela, R.; Piecyk, K.; Stankiewicz-Drogon, A.; Pietrow, P.; Lukaszewicz, M.; Kurpiejewski, K.; Darzynkiewicz, E.; Jankowska-Anyszka, M. N2 modified dinucleotide cap analogs as a potent tool for mRNA engineering. RNA 2023, 29 (2), 200–216. Cornelissen, N. V.; Mineikaitė, R.; Erguven, M.; Muthmann, N.; Peters, A.; Bartels, A.; Rentmeister, A. Post-synthetic benzylation of the mRNA 5′ cap via enzymatic cascade reactions. Chemical Science 2023, 14 (39), 10962–10970, 10.1039/D3SC03822J. DOI: 10.1039/D3SC03822J.

36. Ziemkiewicz, K.; Warminski, M.; Wojcik, R.; Kowalska, J.; Jemielity, J. Quick Access to Nucleobase-Modified Phosphoramidites for the Synthesis of Oligoribonucleotides Containing Post-Transcriptional Modifications and Epitranscriptomic Marks. The Journal of Organic Chemistry 2022, 87 (15), 10333–10348. DOI: 10.1021/acs.joc.2c01390.

37. Lohrmann, R.; Orgel, L. E. Preferential formation of (2’–5’)-linked internucleotide bonds in non-enzymatic reactions. Tetrahedron 1978, 34 (7), 853–855. DOI: 10.1016/0040-4020(78)88129-0.

38. Coleman, T. M.; Wang, G.; Huang, F. Superior 5′ homogeneity of RNA from ATP-initiated transcription under the T7 ϕ2.5 promoter. Nucleic Acids Research 2004, 32 (1), e14–e14. DOI: 10.1093/nar/gnh007 (acccessed 7/12/2023).

39. Roost, C.; Lynch, S. R.; Batista, P. J.; Qu, K.; Chang, H. Y.; Kool, E. T. Structure and Thermodynamics of N6- Methyladenosine in RNA: A Spring-Loaded Base Modification. Journal of the American Chemical Society 2015, 137 (5), 2107–2115. DOI: 10.1021/ja513080v. Kierzek, E.; Kierzek, R. The thermodynamic stability of RNA duplexes and hairpins containing N(6)-alkyladenosines and 2-methylthio-N(6)-alkyladenosines. Nucleic Acids Research 2003, 31 (15), 4472–4480. PMC.

40. Shi, H.; Liu, B.; Nussbaumer, F.; Rangadurai, A.; Kreutz, C.; Al-Hashimi, H. M. NMR Chemical Exchange Measurements Reveal That N6-Methyladenosine Slows RNA Annealing. Journal of the American Chemical Society 2019, 141 (51), 19988–19993. DOI: 10.1021/jacs.9b10939.

41. Mamot, A.; Sikorski, P. J.; Siekierska, A.; de Witte, P.; Kowalska, J.; Jemielity, J. Ethylenediamine derivatives efficiently react with oxidized RNA 3′ ends providing access to mono and dually labelled RNA probes for enzymatic assays and in vivo translation. Nucleic Acids Research 2022, 50 (1), e3–e3. DOI: 10.1093/nar/gkab867 (acccessed 11/18/2022). Depaix, A.; Mlynarska-Cieslak, A.; Warminski, M.; Sikorski, P. J.; Jemielity, J.; Kowalska, J. RNA Ligation for Mono and Dually Labeled RNAs. Chemistry – A European Journal 2021, 27 (47), 12190–12197, 10.1002/chem.202101909. DOI: 10.1002/chem.202101909 (acccessed 2021/11/25).

42. Inagaki, M.; Abe, N.; Li, Z.; Nakashima, Y.; Acharyya, S.; Ogawa, K.; Kawaguchi, D.; Hiraoka, H.; Banno, A.; Meng, Z.;, et al. Cap analogs with a hydrophobic photocleavable tag enable facile purification of fully capped mRNA with various cap structures. Nature Communications 2023, 14 (1), 2657. DOI: 10.1038/s41467-023-38244-8.

43. Liu, C.; Shi, Q.; Huang, X.; Koo, S.; Kong, N.; Tao, W. mRNA-based cancer therapeutics. Nature Reviews Cancer 2023, 23 (8), 526–543. DOI: 10.1038/s41568-023-00586-2. Coulie, P. G.; Van den Eynde, B. J.; van der Bruggen, P.; Boon, T. Tumour antigens recognized by T lymphocytes: at the core of cancer immunotherapy. Nature Reviews Cancer 2014, 14 (2), 135–146. DOI: 10.1038/nrc3670.

44. Schwanhäusser, B.; Busse, D.; Li, N.; Dittmar, G.; Schuchhardt, J.; Wolf, J.; Chen, W.; Selbach, M. Global quantification of mammalian gene expression control. Nature 2011, 473 (7347), 337–342. DOI: 10.1038/nature10098.

45. Lamper, A. M.; Fleming, R. H.; Ladd, K. M.; Lee, A. S. Y. A phosphorylation-regulated eIF3d translation switch mediates cellular adaptation to metabolic stress. Science 2020, 370 (6518), 853–856. DOI: 10.1126/science.abb0993 (acccessed 2023/10/13).

46. Lee, A. S. Y.; Kranzusch, P. J.; Doudna, J. A.; Cate, J. H. D. eIF3d is an mRNA cap-binding protein that is required for specialized translation initiation. Nature 2016, 536 (7614), 96–99. DOI: 10.1038/nature18954.

47. de la Parra, C.; Ernlund, A.; Alard, A.; Ruggles, K.; Ueberheide, B.; Schneider, R. J. A widespread alternate form of cap-dependent mRNA translation initiation. Nature Communications 2018, 9 (1), 3068. DOI: 10.1038/s41467-018-05539-0.

48. Volta, V.; Pérez-Baos, S.; de la Parra, C.; Katsara, O.; Ernlund, A.; Dornbaum, S.; Schneider, R. J. A DAP5/eIF3d alternate mRNA translation mechanism promotes differentiation and immune suppression by human regulatory T cells. Nature Communications 2021, 12 (1), 6979. DOI: 10.1038/s41467-021-27087-w. Shin, S.; Han, M.-J.; Jedrychowski, M. P.; Zhang, Z.; Shokat, K. M.; Plas, D. R.; Dephoure, N.; Yoon, S.-O. mTOR inhibition reprograms cellular proteostasis by regulating eIF3D-mediated selective mRNA translation and promotes cell phenotype switching. Cell Reports 2023, 42 (8), 112868. DOI: 10.1016/j.celrep.2023.112868.

49. Karikó, K.; Muramatsu, H.; Welsh, F. A.; Ludwig, J.; Kato, H.; Akira, S.; Weissman, D. Incorporation of Pseudouridine Into mRNA Yields Superior Nonimmunogenic Vector With Increased Translational Capacity and Biological Stability. Molecular Therapy 2008, 16 (11), 1833–1840. DOI: 10.1038/mt.2008.200. Andries, O.; Mc Cafferty, S.; De Smedt, S. C.; Weiss, R.; Sanders, N. N.; Kitada, T. N1-methylpseudouridine-incorporated mRNA outperforms pseudouridine-incorporated mRNA by providing enhanced protein expression and reduced immunogenicity in mammalian cell lines and mice. Journal of Controlled Release 2015, 217, 337–344. DOI: 10.1016/j.jconrel.2015.08.051.

50. Weissman, D.; Pardi, N.; Muramatsu, H.; Karikó, K. HPLC Purification of In Vitro Transcribed Long RNA. In *Synthetic Messenger RNA and Cell Metabolism Modulation: Methods and Protocols*, Rabinovich, P. M. Ed.; Humana Press, 2013; pp 43–54. Feng, X.; Su, Z.; Cheng, Y.; Ma, G.; Zhang, S. Messenger RNA chromatographic purification: advances and challenges. Journal of Chromatography A 2023, 1707, 464321. DOI: 10.1016/j.chroma.2023.464321. Guimaraes, G. J.; Kim, J.; Bartlett, M. G. Characterization of mRNA therapeutics. Mass Spectrometry Reviews 2023, *n/a* (n/a). DOI: 10.1002/mas.21856 (acccessed 2023/09/01).

