## Supplementary material for "Trinucleotide mRNA cap analog N6-benzylated at the site of posttranscriptional ^m6^Am mark facilitates mRNA purification and confers superior translational properties in vitro and in vivo": S1-Experimental section

[#]*Division of Biophysics, Institute of Experimental Physics, Faculty of Physics, University of Warsaw, 02-089 Warsaw, Poland;* [§]*Centre of New Technologies, University of Warsaw, 02-089 Warsaw, Poland;* [$]*Explorna Therapeutics sp. z o.o. Zwirki i Wigury 93, 02-89 Warsaw, Poland;* [«]*Laboratory of Epitranscriptomics, Department of Environmental Microbiology and Biotechnology, Institute of Microbiology, Faculty of Biology, Biological and Chemical Research Centre, University of Warsaw, 02-089 Warsaw, Poland;* [&]*Department of Chemistry & Biochemistry, University of Delaware, Newark DE 19716;* [♠]*Laboratory of Experimental Medicine, Faculty of Medicine, Medical University of Warsaw, 02-097 Warsaw, Poland;* [♣]*Proteomics Core Facility, IMol Polish Academy of Sciences, 02-247 Warsaw, Poland;* *correspondence to:

### 1. Chemical syntheses

#### 1.1. Reagents

Solvents, chemical reagents, and starting materials were acquired from commercial sources and used without further purification. 5′-*O*-DMTr-2′-*O*-Me-*N*6-phenoxyacetyladenosine 3′-O-phosphoramidite was purchased from Biosearch Technologies or ChemGenes. $^{m6}A_m$ and $^{Bn6}A_m$ phosphoramidites were synthesized as described earlier.(Ziemkiewicz, Warminski, Wojcik, Kowalska, & Jemielity, 2022) Solid support for oligonucleotide syntheses was purchased from GE Healthcare. DNA synthesis grade acetonitrile (<10 ppm of water) was used for the coupling reaction and for washing the solid support.

#### 1.2. Analytical and preparative methods for chemical synthesis of cap analogs

Analytical RP-HPLC was performed on a Gemini® 3 μm NX-C18 LC column (110 Å, 150 × 4.6 mm, 3 μm, flow rate 1.0 mL/min) with linear gradient elution with 0.05 M ammonium acetate buffer pH 5.9 (buffer A) and $1{:}1_{v/v}$ methanol/buffer A (buffer B): Method A: 0-100% in 15 min; Method B: 0-100% in 7.5 min. NMR spectra were recorded at 25°C with a Bruker Avance III HD spectrometer at 500.24 MHz ($^{1}$H NMR) and 202.49 MHz ($^{31}$P NMR) using 5 mm PABBO BB/19F-1H/D Z-GRD probe. The raw NMR data were processed using MestReNova v 12.0.2-20910 Software. The $^{1}$H NMR chemical shifts were calibrated to $CHCl_3$ (7.260 ppm) or $D_2O$ (4.790). For the calibration of $^{31}$P NMR chemical shifts, $H_3PO_4$ was used as an external standard. Signals were assigned based on correlation spectroscopy (COSY) and heteronuclear single quantum coherence ($^{1}$H-$^{13}$C HSQC, $^{1}$H-$^{31}$P HSQC) spectra. High-resolution mass spectra (HRMS) were recorded with a LTQ Orbitrap Velos (ThermoFisher Scientific) spectrometer.

All the compounds were purified by ion-exchange chromatography on DEAE Sephadex A25 ($HCO_3^-$ form) using the Biotage® Selekt Flash Purification System. After loading the column with the reaction mixture and washing it with deionized water, the products were eluted using a linear gradient of triethylammonium bicarbonate (TEAB) in water: 0–0.9 M for dinucleotides (**3a-c**) and 0–1.2 M for trinucleotides (**1, 2a-c**). The fractions containing the desired product were combined, concentrated under reduced pressure, and evaporated to dryness with repeated additions of 96% and then acetonitrile to give a white solid. The trinucleotide cap **1** was further purified by preparative RP-HPLC using a Gemini® 5 μm NX-C18 LC columns (110 Å, 150 × 10 mm, flow rate 5.0 mL/min or 110 Å, 250 × 21.2 mm, flow rate 30 mL/min) with a linear gradient of acetonitrile in 0.05 M ammonium acetate buffer (pH 5.9) and UV detection at 254 nm. After repeated freeze-drying of the collected fractions, the products were isolated as ammonium salts forming white solids. The reaction yields were calculated based on the optical density (mOD = absorbance of the solution × volume in mL) of combined fractions measured in 0.1M phosphate buffer pH 7.0 at 260 nm, using the following extinction coefficients: 34.4 $mM^{-1}cm^{-1}$ for $m^7Gppp^{Bn6}A_mpG$, 32.0 $mM^{-1}cm^{-1}$ for $m^7GpppA_mpG$ and $m^7Gppp^{m6}A_mpG$, 27.1 $mM^{-1}cm^{-1}$ for all $p^*A_mpG$ dinucleotides.

#### 1.3. Synthesis of dinucleotide 5′ phosphates 3a-c ($p^*A_mpG$)

The dinucleotides 3a-c were synthesized according to the protocols described previously.(Sikorski et al., 2020; Ziemkiewicz et al., 2022)

*General procedure:* Synthesis of dinucleotides was performed manually on a 5′-*O*-DMT-2′-*O*-TBDMS-$rG^{iBu}$ 3′-lcaa PrimerSupport 5G (308 μmol/g) solid support (GE Healthcare) placed in a 10 mL syringe equipped with frit filter. In the coupling step, the syringe was closed with a plunger and the resin was gently shaken with a solution of appropriate adenosine phosphoramidite ($A^{Bz}$, $^{m6}A_m^{Pac}$, or $^{Bn6}A_m^{Pac}$,(Ziemkiewicz et al., 2022) 1.2–2.0 equivalents) or biscyanoethyl phosphoramidite (3.0 equivalents) in dry acetonitrile (1.0 mL) and 0.30 M 5-(benzylthio)-1-*H*-tetrazole in acetonitrile (1.5 mL). A solution of 3% (w/v) trichloroacetic acid in dichloromethane was used as a detritylation reagent, and 0.05 M iodine in pyridine/water (9:1) for oxidation. After the last cycle of synthesis, dinucleotides, still attached to the solid support, were washed with 20% (v/v) diethylamine in acetonitrile to remove 2-cyanoethyl protecting groups. Finally, the solid support was washed with acetonitrile and dried under reduced pressure. The product was cleaved from the solid support and deprotected with AMA (methylamine/ammonium hydroxide 1:1 (v/v)) at 37°C for 3 h, evaporated to dryness, and redissolved in DMSO (200 μL). The TBDMS groups were removed using triethylammonium trihydrofluoride (TEA·3HF; 250 µL) and triethylamine (430 µL) at 65°C for 3 h, and then the mixture was cooled down and diluted with 0.05 M $NaHCO_3$ in water (25 mL). The product was isolated by ion-exchange chromatography on DEAE Sephadex (gradient elution 0–0.9 M TEAB) to afford after evaporation triethylammonium salt of dinucleotide p*$A_m$pG. The synthesis scales, yields, HPLC and HRMS data for particular dinucleotides are summarized in **Błąd! Nie można odnaleźć źródła odwołania.**.

**Table 1.** Summary of the synthesis scales, yields, and HRMS data for synthesized dinucleotides.

| Abbreviation | | Synthesis scale [µmol] [a] | Yield [µmol] | m/z calcd. | m/z found |
|---|---|---|---|---|---|
| **3a** | p$A_m$pG | 200 | 142 | 705.11889 | 705.11981 |
| **3b** | $p^{m6}A_m$pG | 200 | 146 | 719.13454 | 719.13537 |
| **3c** | $p^{Bn6}A_m$pG | 200 | 143 | 795.16584 | 795.16658 |

[a] calculated as a product of solid support weight and loading;

##### 1.4. Synthesis of trinucleotide cap analog 1 ($m^7Gppp^{Bn6}A_m$pG, *AvantCap*)

*Procedure A (small scale):* The triethylammonium salt of $p^{Bn6}A_m$pG (843 mOD, 31.1 µmol) and $m^7$GDP-Im (29.3 mg, 46.7 µmol) were suspended in DMSO (620 µL), dry $ZnCl_2$ (67.7 mg, 498 µmol) was added and the mixture was stirred at room temperature for 40 h. The reaction was quenched by adding a solution of $Na_2$EDTA (20 mg/mL; 20 equivalents) and $NaHCO_3$ (10 mg/mL) in water, and the product was isolated by ion-exchange chromatography on DEAE Sephadex (gradient elution 0–1.2 M TEAB) and further purified by RP-HPLC to give, after repeated freeze-drying from water, an ammonium salt of $m^7Gppp^{Bn6}A_m$pG (600 mOD, 18.7 µmol, 60%) as a white solid.

*Procedure B (larger scale):* The triethylammonium salt of $p^{Bn6}A_m$pG (95570 mOD, 3.53 mmol), imidazole (3.84 g, 56.4 mmol), 2,2′-dithiodipyridine (4.65 g, 21.2 µmol) and triethylamine (2.96 mL, 21.2 µmol) were dissolved in DMF (70 mL), then triphenylphosphine (5.54 g, 21.2 µmol) was added and the mixture was stirred at room temperature for 1 h. The product was precipitated by adding a solution of $NaClO_4$ (4.32 g, 35.3 mmol) in cold acetonitrile (700 mL), centrifuged, washed (by resuspension, centrifugation, and decantation) 3 times with cold acetonitrile, and dried under reduced pressure. The solid was suspended in DMF (67 mL), $m^7$GDP (2.28 g, 4.06 mmol) and $ZnCl_2$ (5.52 g, 40.6 mmol) were added, and the mixture was stirred at room temperature for 1 h. The reaction was quenched by adding a solution of $Na_2$EDTA (20 mg/mL; 20 equivalents) and $NaHCO_3$ (10 mg/mL) in water, and the product was isolated by ion-exchange chromatography on DEAE Sephadex (gradient elution 0–1.2 M TEAB) and further purified by RP-HPLC to give, after repeated freeze-drying from water, an ammonium salt of $m^7Gppp^{Bn6}A_m$pG (86430 mOD, 2.51 mmol, 50%) as a white solid.

$m^7Gppp^{Bn6}A_m$pG (**1**): **RP-HPLC** (gradient elution 0-50% MeOH in $CH_3COONH_4$ pH 5.9 in 15 min): $R_t$ = 15.534 min; **HRMS ESI(-)**: *m/z* 1234.19524 (calcd. for $C_{39}H_{48}N_{15}O_{24}P_4^-$ [MH]⁻ 1234.19526); **$^1$H NMR** (500 MHz, $D_2O$) δ = 9.02 (s, 1H), 8.41 (s, 1H), 8.13 (s, 1H), 7.98 (s, 1H), 7.44 – 7.28 (m, 5H), 6.01 (d, *J* = 5.8 Hz, 1H), 5.86 (d, *J* = 3.8 Hz, 1H), 5.84 (d, *J* = 5.8 Hz, 1H), 4.93 (dt, *J* = 8.1, 3.8 Hz, 1H), 4.83 – 4.80 (overlapped with $H_2O$, m, 2H), 4.57 (dd, *J* = 4.9, 3.9 Hz, 1H), 4.54 – 4.49 (m, 2H), 4.47 – 4.42 (m, 2H), 4.39 – 4.15 (m, 9H), 4.01 (s, 3H), 3.42 (s, 3H). **$^{31}$P NMR** (203 MHz, $D_2O$) δ = 0.07 (s, 1P), -10.54 (d, *J* = 18.6 Hz, 1P), -10.65 (d, *J* = 18.3 Hz, 1P), -22.03 (t, *J* = 18.4 Hz, 1P).

##### 1.5. Synthesis of functionalized trinucleotide cap analogs 2a-c ($m^7$Gppp*$A_m$pG-L13$_N$)

*General procedure:* The triethylammonium salt of p*$A_m$pG (**3a-c**) was dissolved in DMSO (to 0.05M solution) and 1,1′-carbonyldiimidazole (CDI, 10 equivalents) was added. The mixture was stirred at room temperature for 2 h and then water (16 equivalents) was added to hydrolyze an excess of CDI. After 20 min, 4,7,10-trioxatridecane-1,13-diamine (3 equivalents) and 1,8-Diazabicyclo[5.4.0]undec-7-ene (DBU, 1 equivalent) were added and the mixture was stirred at room temperature until all the dinucleotide 2′,3′-*O*,*O*-carbonate was consumed (as evidenced by RP-HPLC). The crude product – a regioisomeric mixture of 2′-*O*/3′-*O*-carbamoyl dinucleotide *P*-imidazolide (Im-p*$A_m$pG-L13$_N$) – was isolated as a sodium salt by the addition of a $NaClO_4$ (6 equivalents) solution in cold acetonitrile (10 times volume of DMSO). The white precipitate was centrifuged, washed with cold acetonitrile three times and dried under reduced pressure. The crude Im-p*$A_m$pG-L13$_N$, $m^7$GDP (2 equivalents) and $ZnCl_2$ (16 equivalents) were suspended in DMSO (to

0.025 M concentration of Im-p*$A_m$pG-$L13_N$) and the mixture was stirred at room temperature for 3–7 days. The reaction was quenched by adding a solution of $Na_2$EDTA (20 mg/mL; 20 equivalents) and $NaHCO_3$ (10 mg/mL) in water, and the product was isolated by ion-exchange chromatography on DEAE Sephadex (gradient elution 0–1.2 M TEAB) to afford – after evaporation – triethylammonium salt of $m^7$Gppp*$A_m$pG-$L13_N$. The products were contaminated with $m^7$G mononucleotide, but since it lacks primary amine groups, it does not interfere with subsequent immobilization on the Sepharose. The synthesis scales, yields, HPLC and HRMS data for particular cap analogs are summarized in **Błąd! Nie można odnaleźć źródła odwołania.**.

**Table 2.** Summary of the synthesis scales, yields, HPLC, and HRMS data for functionalized trinucleotide cap analogs.

| Abbreviation | | Synthesis scale [µmol] [a] | Yield [µmol] | RP-HPLC $R_t$ [min] [b] | m/z calcd. | m/z found [c] |
|---|---|---|---|---|---|---|
| **2a** | $m^7$Gppp$A_m$pG-$L13_N$ | 13.5 | 4.75 | 10.842*<br>11.845* | 1390.30626 | 1390.30808<br>1390.30788 |
| **2b** | $m^7$Gppp$^{m6}$$A_m$pG-$L13_N$ | 10.0 | 3.22 | 6.909**<br>7.408** | 1404.32191 | 1404.32163<br>1404.32154 |
| **2c** | $m^7$Gppp$^{Bn6}$$A_m$pG-$L13_N$ | 19.1 | 12.6 | 9.460**<br>9.684** | 1480.35321 | 1480.35484<br>739.67388$ |

[a] based on the amount of p*$A_m$pG used for the reaction; [b] HPLC conditions: * Method A, ** Method B, see *General information*; [c] The 2′-*O*/3′-*O* regioisomers were characterized separately; $ z=2

##### 1.6. Synthesis of affinity resins AR-1, AR-2, and AR-3

The affinity resins were synthesized based on the previously published protocol (Szczepaniak, Zuberek, Darzynkiewicz, Kufel, & Jemielity, 2012). Briefly, a suspension of Sepharose™ CL-4B (3 mL of settled resin) in water (6 mL total volume) was cooled down in an ice-water bath and a solution of BrCN (386 mg) in acetonitrile (386 µL) was added. The mixture was stirred in an ice-water bath and the pH was kept at the level of 11 by addition of 1 M NaOH (ca. 200 µL every 1–5 min). After 2 h the slurry was filtered on a Buchner funnel (not letting the resin get dry) and washed with cold water (200 mL), and then with 0.1 M carbonate buffer pH 9 (200 mL). The resin was transferred into a vial and suspended in carbonate buffer pH 9 (3 mL). A solution of trinucleotide cap analog **2a, 2b**, or **2c** (32.0 mOD, 1.0 µmol) in water (100 µL) was added, and the mixture was stirred gently at 4°C overnight. The slurry was filtered, washed with water (150 mL) and 20% ethanol (150 mL). The flow-through was analyzed by RP-HPLC to estimate the resin loading (usually 0.9–1.0 µmol/mL of the settled resin). The resins were stored in 20% ethanol (3 mL – total volume of ca. 6 mL) at 4°C.

#### 2. mRNA synthesis, purification, and quality control

##### 2.1. Preparation of DNA plasmid vectors

Luciferase gene (Fluc) from firefly (Lampyridae) was purchased from Invitrogen (ThermoFisher Scientific) with restriction sites for AdeI and BamHI endonucleases, and cloned into pJet1.2 (ThermoFisher Scientific) plasmid vector using sticky-end cloning method. The ordered plasmid with Fluc sequence (2 µg) was digested (15 min, 37°C) with AdeI (ThermoFisher Scientific) and BamHI (ThermoFisher Scientific) restriction enzymes and 10× Fast Digest Buffer (ThermoFisher Scientific). The digested DNA was then separated on an 1% agarose gel and the strand containing the Fluc sequence was excised and purified using a commercial DNA purification kit (Macherey-Nagel) according to the protocol. The pJet1.2 plasmid vector (2 µg) was linearized (16 h, 37°C) with AarI restriction enzyme (ThermoFisher Scientific), 10× AarI buffer (ThermoFisher Scientific) and 50× Oligo (ThermoFisher Scientific) and purified using a commercial DNA purification kit (Macherey-Nagel). Then the pJET1.2 solution including 100 ng of DNA was mixed with the insert (Fluc sequence) at a molar ratio of 1:3 (vector- insert), added 10× Ligase buffer (ThermoFisher Scientific) and T4 DNA Ligase (ThermoFisher Scientific), incubated (1 h, 25°C), and then reaction mix (20 µl) cooled to 4°C and incubated an additional 1 h. Half of the ligation mixture (10 µl) was mixed with pre-melted on ice commercially available chemo competent bacteria (50 µl) Top10 (ThermoFisher Scientific) and incubated on ice for 30 min. The mixture was transferred to 42°C (heat shock) and cooled on ice for 2 min and 500 µl of SOC outgrowth medium (ThermoFisher Scientific) was added and incubated at 37°C for 1 h with shaking (300 RPM). The transformation mixture (200 µl) was spread on LB-agar plate (Roth) with 100 mg/mL Ampicillin (Roth) and incubated (37°C, 16 h). Then, single colonies were selected and inoculated with liquid LB medium (Roth) in a volume of 5 mL and supplemented with Ampicillin (Roth) at a concentration of 100 mg/mL and the cultures were incubated (37°C, 16 h). The bacterial cultures were then centrifuged (4000g, 10 min) and the plasmids were purified using the commercial GeneJET Plasmid Miniprep Kit (ThermoFisher Scientific). Concentration was measured using a Nanodrop 2000c spectrophotometer (ThermoFisher Scientific) and plasmids were sent for DNA sequencing using the Sanger method (Genomed). The pJet1.2_FLuc plasmid contained a short form of the poly(A) tail ~30 adenine nucleotide. In order to obtain a DNA template with a poly(A) tail consist of 90 adenine nucleotides, a several-step procedure was carried out to insert an adenine oligonucleotide into the 3′ end of Fluc sequence. The insert was designed as a double-stranded DNA oligonucleotide and added to the pJet plasmid vector that encodes Fluc using a blunt end cloning method. Solutions of two DNA oligonucleotides (Genomed) with sequence: $A_{60}$ (coding strand); $T_{60}$ (template strand) were mixed in a 1:1 ratio (final 50 µM of each DNA strand) and an enzyme reaction was set up to phosphorylate the 5' ends of the oligonucleotides by adding T4 Polynucleotide Kinase (NEB) and 10× buffer for T4 Polynucleotide Kinase (NEB). The reaction was carried out for 30 minutes at 37°C. The strands were then hybridized by heating to 95°C and slowly cooling to 25°C for 2 h, step gradient ~2°C/~3 min. The insert DNA was purified using a commercial DNA purification kit (Macherey-Nagel) according to the protocol. The circular plasmid pJET1.2_Fluc that encodes firefly luciferase (2 µg) was linearized (16 h, 37°C) with AarI restriction enzyme (ThermoFisher Scientific), 10× AarI buffer (ThermoFisher Scientific) and 50× Oligo (ThermoFisher Scientific) and purified using a commercial DNA purification kit (Macherey-Nagel). The enzymatic reaction was then prepared with DNA Polymerase I, Large Fragment (Klenow), 10× buffer 3 (NEB), and 10 mM NTP (ThermoFisher Scientific) to remove 3′ overhangs and filling in 5′ overhangs to form blunt ends (15 min, 25°C) and again purified the vector using a commercial DNA purification kit (Macherey-Nagel). Finally, dephosphorylation reaction was performed using FastAP Thermosensitive Alkaline Phosphatase (ThermoFisher Scientific) and 10× Fast AP buffer (ThermoFisher Scientific) by incubating (10 min, 37°C) and purifying the final vector using a commercial DNA purification kit (Macherey-Nagel). The final vector pJET1.2_Fluc solution including 100 ng of DNA was mixed with the insert at a molar ratio of 1:3 (vector- insert), added 10× Ligase buffer (ThermoFisher Scientific), PEG 2000 (ThermoFisher Scientific), ATP (ThermoFisher Scientific) and T4 DNA Ligase (ThermoFisher Scientific) and incubated (1 h, 25°C) and then reaction mix (20 µl) cooled to 4°C and incubated an additional 1 h. Then transformation and clonal selection as well as DNA sequencing were carried out analogous to the procedure described above.

Human erythropoietin gene (hEPO), *Gaussia* luciferase gene (Gluc) and ovalbumin (OVA) were purchased from Invitrogen (ThermoFisher Scientific) with restriction sites for AdeI and BamHI endonucleases, and was cloned (analogously to the method for Fluc) into pJet1.2 (ThermoFisher Scientific) plasmid vector using sticky-end cloning method. Then, an analogous procedure to that described above was carried out to extend the poly(A) tail for the pJet1.2_EPO, pJet1.2_Gluc and pJet1.2_OVA plasmids to the final poly(A) tail length of 90 adenine nucleotides. α1-antitrypsin gene (hA1AT) and NY-ESO1 gene were purchased from Invitrogen (ThermoFisher Scientific) in pUC57 plasmid vector. Then, an analogous procedure to that described above was carried out to extend the poly(A) tail for the pUC57_A1AT and pUC57_NY-ESO1 plasmids to the final poly(A) tail length of 90 adenine nucleotides.

##### 2.2. Capping efficiency of short RNAs

Short RNAs were obtained by IVT using T7 class II promoter Φ2.5 (initiated by ATP – TAATACGACTCACTATT**A**) or class III promoter Φ6.5 (initiated by GTP – TAATACGACTCACTATA**G**). The typical IVT reaction (100 µL) was incubated for 4 h at 37°C and contained: 3 mM each of UTP/GTP/CTP for Φ2.5 or UTP/ATP/CTP for Φ6.5, 0.75 mM of ATP or GTP (NTPs were purchased from ThermoFisher Scientific Scientfic), respectively, and 6 mM of trinucleotide cap analog ($m^7GpppA_mpG$, $m^7Gppp^{m6}A_mpG$ or $m^7Gppp^{Bn6}A_mpG$), 1.25 µM of annealed oligonucleotides as DNA template, 1 U/µL RiboLock RNase Inhibitor (ThermoFisher Scientific), 20 mM $MgCl_2$, RNA Polymerase Buffer (ThermoFisher Scientific) and T7 RNA polymerase (0.3 mg/mL, in-house prepared). The preparation of uncapped RNAs was performed as before, but with 3 mM of ATP/GTP and without the presence of cap analog. After 4 hours sample was treated with DNase I (5 U, ThermoFisher Scientific) and incubated for another 30 min at 37°C. The reaction mixture was stopped by addition of equimolar amount of $Na_2EDTA$ water solution (EDTA to $Mg^{2+}$) and purified using Monarch® RNA Cleanup Kit (NEB). Such prepared RNA sample was then purified using HPLC with Phenomenex® Clarity 3 μM Oligo-RP column with a linear gradient of buffer B (0.2 M triethylammonium acetate pH 7.0 and acetonitrile, v/v) from 10% to 33.3% in buffer A (0.1 M triethylammonium acetate pH 7.0) in 35 min at 1 mL/min. Collected fractions of RNA were then precipitated with 0.33 M NaOAc pH 5.2 and 2.5 volumes of 80% ethanol. Moreover, to obtain homogenous 3´ end of synthetized RNAs, obtained 35-nt long uncapped and 36- or 37-nt long capped transcripts (1 μM) were incubated with 1 μM DNAzyme 10-23 (TGATCGGCTAGGCTAGCTACAACGAGGCTGGCCGC) in 50 mM Tris pH 8.0 and 50 mM $MgCl_2$ for 1 hour at 37°C, purified using HPLC and precipitated as above. The quality of obtained short RNAs and capping efficiency were checked on 15% acrylamide / 7 M urea / 1× TBE gel with SYBR Gold (Invitrogen) staining and the concentration was determined spectrophotometrically using NanoDrop 2000c. The PAGE band intensities corresponding to capped and uncapped RNAs were quantified densitometrically using 1D Gel Image Analysis Software TotalLab CLIQS.

##### 2.3. Optimization of in vitro transcription conditions for improvement of capping efficiency with $m^7Gppp^{Bn6}A_mpG$

The conditions of the IVT have been optimized to achieve ~90% capping efficiency with $m^7Gppp^{Bn6}A_mpG$. For these studies DNA template encoding ~1000 nt long transcript (NY-ESO1) was chosen to ensure sufficient separation of capped and uncapped mRNA using analytical RP-HPLC.

In the first set of optimization experiments capping efficiency was determined for mRNA obtained in IVT at various pH, ranging from 6.0 to 9.0. 20 μL of each prepared IVT mix contained following components at indicated final concentration: transcription buffer prepared in two sets of variants: 1) containing 40 mM Bis-Tris at the pH ranging from 6.0 to 7.0, and 2) containing 40 mM Tris-HCl at the pH ranging from 7.0 to 9.0, 2 mM spermidine, 1 U/μL RNase inhibitor (RiboLock, ThermoFisher Scientific), 5 mM ATP, CTP and UTP, 4 mM GTP, 10 mM DTT, 25 mM $MgCl_2$, 0.002 U/μL inorganic pyrophosphatase (ThermoFisher Scientific), 10 mM $m^7Gppp^{Bn6}A_mpG$, 40 ng/μL linearized plasmid as the DNA template and 0.125 μg/μL T7 RNA polymerase (in-house prepared). All components were gently mixed by pipetting and incubated for one hour at 37°C. To remove DNA template, IVT mix was incubated for next 30 min at 37°C with 0.025 U/μL DNase I (ThermoFisher Scientific). The enzymes in the IVT mix were inactivated by addition of one volume of 52 mM EDTA, and full-length mRNAs were isolated from abortive transcripts using oligo dT chromatography (POROS™ Oligo $(dT)_{25}$ Affinity Resin, ThermoFisher Scientific) according to manufacturer's protocol. In short, crude IVT mix was diluted with high salt buffer (500 mM NaCl, 10 mM Tris-HCl pH 7.5, 1 mM EDTA) and loaded on the accordingly equilibrated $oligo(dT)_{25}$ resin. After binding step (10 min, RT), resin was washed with high and low salt buffer (200 mM NaCl, 10 mM Tris-HCl pH 7.5, 1 mM EDTA), respectively. mRNA was eluted with RNase-free water heated to 65°C. At this step IVT efficiency was estimated by measuring the volume and the absorbance of eluted mRNA at 260 nm (NanoDrop $One^C$, ThermoFisher Scientific) and calculated as mRNA yield obtained from 1 μL of starting IVT volume.

Capping efficiency was determined using RP-HPLC method based on delayed retention of mRNA containing $m^7Gppp^{Bn6}A_mpG$ in a linear gradient of organic solvent. Agilent Infinity 1260 II instrument and RNASep™ Prep (ADS Biotec) was used for analysis of 1 μg of each mRNA sample initially purified by $oligo(dT)_{25}$. Sufficient separation of capped and uncapped mRNA was obtained for a linear gradient of acetonitrile (10-14.5% in 22 min at 0.9 mL/min flow rate) in 0.1 M triethylammonium acetate (TEAA) pH 7.0 at 55°C. Capping efficiency was determined based on the area of automatically integrated peaks (using Agilent OpenLAB CDS ChemStation Edition C.01.10), referring each of the peak to capped and uncapped mRNA. To confirm results obtained using RP-HPLC, capping efficiency was also determined by ribozyme-based assay (see *2.7 Capping efficiency of full-length mRNAs*).

Second set of experiments was performed for the buffers at pH in the range of 6.0-7.0 with 0.25 intervals. mRNA yield was estimated as described previously, and for RP-HPLC analysis bioZen™ 2.6 μm Oligo LC column (Phenomenex®) was used at the linear gradient of acetonitrile (10-20% in 50 min at 0.5 mL/min flow rate) in 0.1 M TEAA pH 7.0 at 55°C.

#### 2.4. In vitro transcription protocols

Depending on the application, mRNA was *in vitro* transcribed and purified following one of optimized IVT protocols:

**Protocol A**

IVT mix contained following components at indicated final concentration: 40 mM Tris-HCl pH 7.9, 2 mM spermidine, 1 U/µL RNase inhibitor (RiboLock, ThermoFisher Scientific), 2 mM ATP, CTP and UTP, 1 mM GTP, 1 mM DTT, 10 mM $MgCl_2$, 0.002 U/µL inorganic pyrophosphatase (ThermoFisher Scientific), 2 mM $m^7GpppA_mpG$ or $m^7Gppp^{Bn6}A_mpG$, 50 ng/µL linearized plasmid as the DNA template and 0.125 µg/µL T7 RNA polymerase (*in-house* prepared). All components were gently mixed by pipetting and incubated for 2 h at 37°C. After DNA template removal (incubation with 0.025 U DNase I/µL IVT for 30 min at 37°C) the crude mRNA sample was initially purified using Monarch® RNA Cleanup Kit (New England Biolabs).

Next, uncapped transcripts were enzymatically removed from the sample by two-step reaction. First, initially purified mRNA was treated with 5´ polyphosphatase (Lucigen), which hydrolizes 5´ triphosphate to 5´ monophosphate. Typical reaction mix contained: 30 µg of mRNA, 2 U/µL RNA 5´ polyphosphatase, 1× reaction buffer (50 mM HEPES-KOH pH 7.5, 100 mM NaCl, 1 mM EDTA, 0.1% β-mercaptoethanol, 0.01% Triton X-100) and 1 U/µL RNase inhibitor (RiboLock, ThermoFisher Scientific). After 1 h incubation at 37°C, each sample was purified using Monarch® RNA Cleanup Kit (New England Biolabs). In the second step uncapped mRNA molecules with 5´ monophosphate were degraded by XRN-1 5´→3´ exonuclease (New England Biolabs). The reaction mix contained: 30 µg of mRNA, 0.5 U/µL XRN-1, 1× reaction buffer (50 mM Tris-HCl pH 7.9, 100 mM NaCl, 10 mM $MgCl_2$, 1 mM DTT) and 1 U/µL RNase inhibitor (RiboLock, ThermoFisher Scientific). After 2 h incubation at 37°C, each sample was purified using Monarch® RNA Cleanup Kit (New England Biolabs).

To remove contaminants, such as dsRNA and short mRNA species, second purification step was applied. Each mRNA sample was purified by RP-HPLC on Shimadzu Nexera Prep using RNASep™ Prep column (ADS Biotec). mRNA was eluted with the linear gradient (10-14.5%) of acetonitrile in 0.1 M TEAA pH 7.0 at 55°C. Eluate quality was analyzed on 1× TBE 1.2% agarose gel, and the fractions containing mRNA of the highest purity were combined, precipitated overnight at -20°C with 0.3 M NaOAc pH 5.2 and one volume of isopropanol, and washed with 80% ethanol. Prior further analysis, mRNA pellets were resuspended in RNase-free water.

**Protocol B**

IVT mix contained following components at indicated final concentration: 40 mM Tris-HCl pH 7.9, 2 mM spermidine, 1 U/µL RNase inhibitor (RiboLock, ThermoFisher Scientific), 5 mM ATP, CTP and UTP, 4 mM GTP, 10 mM DTT, 25 mM $MgCl_2$, 0.002 U/µL inorganic pyrophosphatase (ThermoFisher Scientific), 10 mM $m^7GpppA_mpG$ or $m^7Gppp^{Bn6}A_mpG$, 40 ng/µL linearized plasmid as the DNA template and 0.125 µg/µL T7 RNA polymerase (*in-house* prepared). All components were gently mixed by pipetting and incubated for 1 h at 37°C. After DNA template removal (incubation with 0.025 U DNase I/µL IVT for 30 min at 37°C), enzymes in IVT mix were inactivated by addition of one volume of 52 mM EDTA, and the crude mRNA sample was initially purified using oligo$(dT)_{25}$ resin (as previously described) and concentrated by ultrafiltration (Amicon Ultra-15 100K, Millipore) prior loading on RP-HPLC column.

To remove contaminants, such as dsRNA and short mRNA species, second purification step was applied. Each mRNA sample was purified by RP-HPLC on an Agilent Infinity 1260 II using RNASep™ Prep or RNASep™ Semi-Prep column (ADS Biotec). mRNA was eluted with the linear gradient (10-14.5%) of acetonitrile in 0.1 M TEAA pH 7.0 at 55°C. Eluate quality was analyzed on 1× TBE 1.2% agarose gel, and the fractions containing mRNA of the highest purity were combined, desalted by ultrafiltration (Amicon Ultra-15 100K, Millipore), precipitated overnight at -20°C with 0.3 M NaOAc pH 5.2 and one volume of isopropanol, and washed with 80% ethanol. Prior further analysis, mRNA pellets were resuspended in RNase-free water.

**Protocol C**

IVT mix contained following components at indicated final concentration: 40 mM Tris-HCl pH 7.9 (for IVT with $m^7GpppA_mpG$) or 40 mM Bis-Tris pH 6.5 (for IVT with $m^7Gppp^{m6}A_mpG$ or $m^7Gppp^{Bn6}A_mpG$), 2 mM spermidine, 1 U/µL RNase inhibitor (RiboLock, ThermoFisher Scientific), 5 mM ATP, CTP and UTP, 4 mM GTP, 10 mM DTT, $MgCl_2$ (10 mM for IVT with $m^7GpppA_mpG$; 25 mM for IVT with $m^7Gppp^{m6}A_mpG$ or $m^7Gppp^{Bn6}A_mpG$), 0.002 U/µL inorganic pyrophosphatase (ThermoFisher Scientific), cap analog (10 mM $m^7GpppA_mpG$ or $m^7Gppp^{Bn6}A_mpG$; 12 mM $m^7Gppp^{m6}A_mpG$), 40 ng/µL linearized plasmid as the DNA template and 0.125 µg/µL T7 RNA polymerase (*in-house* prepared). All components were gently mixed by pipetting and incubated for 1 h at 37°C. After DNA template removal (incubation with 0.025 U DNase I/µL IVT for 30 min at 37°C), enzymes in IVT mix were inactivated by addition of one volume of 52 mM EDTA, and the crude mRNA sample was initially purified using oligo$(dT)_{25}$ resin (as previously described) and concentrated by ultrafiltration (Amicon Ultra-15 50K or 100K, Millipore) prior loading on preparative column.

To remove contaminants, such as dsRNA and short mRNA species, second purification step was applied. Each mRNA sample was purified by RP-HPLC on an Agilent Infinity 1260 II using RNASep™ Semi-Prep columns (ADS Biotec). mRNA was eluted with the linear gradient (10-14.5%) of acetonitrile in 0.1 M TEAA pH 7.0 at 55°C. Eluate quality was analyzed on 1× TBE 1.2% agarose gel, and the fractions containing mRNA of the highest purity were combined, desalted by ultrafiltration (Amicon Ultra-15 50K or 100K, Millipore), precipitated overnight at -20°C with 0.3 M NaOAc pH 5.2 and one volume of isopropanol, and washed with 80% ethanol. Prior formulation and quality control, mRNA pellets were resuspended in RNase-free water to obtain concentration of >1 µg/µL and – if necessary – stored at -80°C as single-use aliquots.

##### 2.5. RP-HPLC analysis of mRNA purity and capping efficiency

Quality of the mRNA samples after purification process and capping efficiency for mRNAs with $m^7Gppp^{Bn6}A_mpG$ were verified by RP-HPLC using Agilent Infinity II 1260 instrument and RNASep™ Prep 7.8 × 50 mm (ADS Biotec) or bioZen™ 2.6 µm Oligo LC column 100 × 2.1 mm (Phenomenex®). 1 µg of each mRNA sample was analysed by applying linear gradient of acetonitrile (for RNASep™ Prep column: 10-14.5% in 22 min at 0.9 mL/min flow rate; for bioZen™ column: 10-20% in 50 min at 0.5 mL/min) in 0.1 M triethylammonium acetate (TEAA) pH 7.0 at 55°C. Capping efficiency was determined based on the area of automatically integrated peaks (using Agilent OpenLAB CDS ChemStation Edition C.01.10), referring each of the peak to capped and uncapped mRNA.

##### 2.6. Analysis of double-stranded RNA content

Double-stranded RNA (dsRNA) impurities in mRNA samples were detected and semi-quantified using dot-blot assay. For each sample dsRNA presence was determined 1) after IVT and initial purification using oligo$(dT)_{25}$ resin (to check the influence of cap analog used for IVT on the dsRNA formation) or 2) after purification using RP-HPLC or cellulose (to verify purity of the sample prior mRNA application). To detect even traces of dsRNA impurities in the samples dedicated for in vivo experiments, analyzed portions contained 250 and 2500 ng of mRNA purified by RP-HPLC or cellulose. Analyzed amounts of the samples further used in the in vitro experiments were reduced to 25 and 250 ng. To avoid signal saturation, samples analyzed after initial purification using oligo$(dT)_{25}$ resin contained 5 and 25 ng of mRNA.

For each analysis intended amounts of mRNA were diluted to obtain final volume of 25 – 50 µL, which was further blotted on the Amersham™ Hybond™-N$^+$ membrane (GE Healthcare) soaked with 1× TBST using Bio-Dot microfiltration apparatus (Bio-Rad). Membrane with UV cross-linked mRNA was blocked for 1 h at RT with 5% non-fat dried milk in 1× TBST buffer and subsequently incubated overnight at 4°C with monoclonal mouse anti-dsRNA J2 antibody (SCICONS) diluted 1:5000 with 1× TBST buffer containing 2% non-fat dried milk. After incubation with primary antibody the membrane was washed 3 × 5 min with 1× TBST at RT and incubated for 1 h at RT with polyclonal anti-mouse IgG secondary antibody conjugated with horseradish peroxidase (Cell Sygnaling Technology) diluted 1:3000 with 1× TBST buffer containing 2% non-fat dried milk. Then, the membrane washed 3 × 5 min with 1× TBST at RT was incubated with Amersham™ ECL™ Prime Western Blotting Detection Reagent (Cytiva), and the chemiluminescence signal was detected on Amersham™ ImageQuant 800 (GE Healthcare) and quantified using ImageQuantTL software (GE

Healthcare), by the comparison to dsRNA standard in the range of 0.078-10 ng (Abnova), 0.062-8 ng, 0.156-20 ng or 0.391-50 ng (New England Biolabs), prepared separately for each membrane.

##### 2.7. Capping efficiency of full-length mRNAs

Capping efficiency could differ depending on nucleotide sequence of the template, therefore efficiency of $m^7GpppA_mpG$ and $m^7Gppp^{Bn6}A_mpG$ cap analogs incorporation was determined individually for the most of the transcripts.

For that purpose, ribozyme complementary to the 5´UTR sequence was designed and used for cleavage of mRNA close to the 5´ end. 10 µL of analyzed sample containing >10 µg (>100 nM) of mRNA was mixed with 1.5 µL of hybridization buffer containing 100 mM Tris-HCl pH 7.5 and 50 mM NaCl and 2 µL of 10 µM ribozyme. The mixture was incubated at 95°C for 2 min, and subsequently at RT for 10-15 min. Cleavage reaction with the ribozyme hybridized to mRNA was initiated by addition of 1.5 µL of 100 mM $MgCl_2$, conducted at 37°C for 1 h, and quenched by addition of 2 µL 100 mM EDTA.

To analyze cleaved RNA fragments 3 µL of quenched reaction was mixed with equal volume of loading dye (8 M urea, 50% formamide, 20 mM EDTA, 0.03% bromophenol blue, 0.03% xylene cyanol), heat denatured at 95°C for 3 min and loaded onto 15% polyacrylamide gel with 7 M urea and 1× TBE. The gel after electrophoresis was incubated at RT for 15 min with 50 mL of staining reagent (SYBR® Gold, Invitrogen, diluted 1:10000), and visualized using Typhoon scanner (GE Healthcare). Migration of short RNA fragments being the result of ribozyme cleavage differs depending on the cap structure present on the 5´ end of mRNA, and the bands intensities correlate with the amount of the RNA. Therefore, the intensity of the bands corresponding with $m^7GpppA_mpG$-RNA, $m^7Gppp^{Bn6}A_mpG$-RNA and pppG-RNA were quantified densitometrically using ImageQuantTL software (GE Healthcare). Percentage of capped mRNA in each sample was determined as the ratio of $m^7GpppA_mpG$-RNA or $m^7Gppp^{Bn6}A_mpG$-RNA and the summarized intensities for capped RNA and pppG-RNA.

#### 3. mRNA expression studies in cultured mammalian cells

##### 3.1. Cell lines

HEK293T (CRL-3216), A549 (CCL-185), and CT26 (CRL-2638) were purchased from ATCC. HEK293T and A549 cells were cultured in DMEM (ThermoFisher Scientific), supplemented with 10% Fetal Bovine Serum (Cytiva), 1% Penicillin-Streptomycin (ThermoFisher Scientific), 2 mM L-glutamine (ThermoFisher Scientific, 25030081) and 1 mM Sodium Pyruvate (ThermoFisher Scientific, 11360070). CT26 cells were cultured in RPMI-1640 (ThermoFisher Scientific), supplemented with 10% Fetal Bovine Serum (Cytiva), 1% Penicillin-Streptomycin (ThermoFisher Scientific), 2 mM L-glutamine (ThermoFisher Scientific) and 1 mM Sodium Pyruvate (ThermoFisher Scientific). All cells were grown at 37°C, 100% humidity and 5% $CO_2$. For mRNA transfection, 10,000 cells/well (HEK293T, CT26) or 5,000 cells/well (A549) were seeded on the previous day on 96-well plate in 100 µl of appropriate complete medium.

##### 3.2. Isolation and culture of murine bone marrow-derived macrophages

Bone marrow was spun out from the femurs of 6-week-old female C57BL/6 mice. RBCs were lysed using ACK (Ammonium-Chloride Potassium) Lysing Buffer (ThermoFisher Scientific Scientific) according to the manufacturer's protocol. All the remaining cells were suspended at $1 \times 10^6$/mL density in the culture medium [RPMI 1640 medium (Sigma-Aldrich) supplemented with heat-inactivated 10% (v/v) fetal bovine serum (HyClone), 2 mM ʟ-glutamine (Sigma-Aldrich), 100 U/mL penicillin and 100 µg/mL streptomycin (both from Sigma-Aldrich)], plated at ø10 cm non tissue culture-treated Petri dishes (Sarstedt) and placed at 37°C in the atmosphere of 5% CO2 in the air. Differentiation towards macrophages was induced with 50 ng/mL recombinant murine M-CSF (Peprotech). On day 11th the cells were collected for subsequent experiments.

##### 3.3. Isolation and culture of murine bone marrow-derived dendritic cells

Bone marrow was flushed with cold PBS from the femurs of 6-week-old female C57BL/6 mice. RBCs were lysed using ACK (Ammonium-Chloride Potassium) Lysing Buffer (ThermoFisher Scientific Scientific) according to the manufacturer's protocol. All the remaining cells were suspended at $1\times10^6$/mL density in the culture medium [RPMI 1640 medium (Sigma-Aldrich) supplemented with heat-inactivated 10% (v/v) fetal bovine serum (HyClone), 2 mM L-glutamine (Sigma-Aldrich), 100 U/mL penicillin (Sigma-Aldrich) and 100 µg/mL streptomycin (Sigma-Aldrich), 1% (v/v) MEM non-essential amino acids solution (Thermo Fisher Scientific), and 1 mM sodium pyruvate (Sigma-Aldrich), plated at ø10 cm non-tissue culture-treated Petri dishes (Sarstedt) and placed at 37°C in the atmosphere of 5% $CO_2$ in the air. Differentiation towards DCs was induced with 20 ng/mL of recombinant murine GM-CSF and 10 ng/mL of recombinant murine IL-4 (both from Peprotech). On day 4$^{th}$ 5 mL of fresh cytokine-supplemented medium was added for 3-day incubation. On day 9$^{th}$ loosely adherent and floating cells were collected and used for subsequent experiments.

##### 3.4. In vitro transfection of the cells with synthetic mRNA

Transfection of cells with mRNA encoding Fluc or hEPO was done using Lipofectamine MessengerMAX Reagent (ThermoFisher Scientific) according to manufacturer's protocol. Briefly, 0.15 µL of Lipofectamine MessengerMAX Reagent was diluted in 5 µL of Opti-MEM Medium (ThermoFisher Scientific), mixed and incubated for 10 minutes at RT. Next, 50 ng of mRNA was diluted in 5 µL Opti-MEM Medium and mixed well. Diluted mRNA mix was added to diluted Lipofectamine MessengerMAX™ Reagent (1:1 ratio) and incubated for 5 minutes at RT. mRNA-lipid complexes were then added to cells and incubated for designated amount of time at 37°C, 100% humidity, and 5% $CO_2$.

Transfection with mRNA for hA1AT was performed using Lipofectamine MessengerMAX Reagent according to manufacturer's protocol. Briefly, 3.75 µL Lipofectamine MessengerMAX Reagent was diluted in 125 µL Opti-MEM Medium and mixed well and incubated in Opti-MEM Medium for 10 minutes in RT. Next, mRNA was diluted by adding appropriate amount to 125 µL Opti-MEM Medium and mix well. Diluted mRNA mix (125 µL) was added to 125 µL of diluted Lipofectamine MessengerMAX™ Reagent (1:1 ratio) and incubated for 5 minutes at RT. mRNA-lipid complexes were then added to cells (HEK293T or A549) and incubated for 24 h and 48 h at 37°C. After this time cells and cell culture medium were collected for Western blot, EnzChek® Elastase Assay (ThermoFisher Scientific) and ELISA (Innovative Research).

##### 3.5. Fluc activity detection

Fluc activity was measured in cell lysates collected 6 h and 24 h post transfection. Briefly, cell media were removed and 100 µl of BrightGlo Reagent (Promega) was added to cells for 5 min incubation at RT in dark. Luminescence signal (Relative Light Units, RLU) was measured in EnVision plate reader (Perkin-Elmer).

##### 3.6. hEPO ELISA

hEPO concentrations in cell culture media was measured with hEPO ELISA kit (ThermoFisher Scientific, BMS2035-2) according to manufacturer instructions. Dilutions were made with sample diluent mixed with distilled water (1:1). The diluted samples were added in duplicates to washed ELISA wells precoated with anti-hEPO antibodies on a 96-well ELISA plate. Standard curve was prepared from 7 recombinant hEPO standard dilutions made according to ELISA kit instruction. The absorbance was measured in EnVision plate reader (Perkin-Elmer, 450 nm primary wave length and 620 nm reference). The mean values of absorbance from each samples were calculated in relation to absorbance values of the standard curve. The concentrations of hEPO in samples were calculated by interpolation of absorbance values for each sample from standard curve using GraphPad Prism 9.0. Obtained hEPO concentration values were expressed in mIU/mL and converted into ng/mL (assuming that 1 ng of hEPO is equal to 119.04 mIU).

##### 3.7. hA1AT detection

*Western blot analysis*

Cells in 6-well format were washed with 1× PBS. 100 µL/well of ice-cold lysis buffer (Cell Signaling) with protease (Roche) and phosphatase (Roche) inhibitor cocktail was added to each well and incubated on ice for 5 minutes. Next, cells were scraped off the plate and centrifuged for 10 minutes at 13,000 × g in a cold microfuge. Supernatants were collected for further Western blot analysis. Protein concentrations were measured using Pierce™ BCA Protein Assay Kit (ThermoFisher Scientific), and each samples were mixed

3:1 with 4× loading buffer and heated to 95°C for 10 min. Lysates were then separated by SDS-PAGE (30ug/lane), transferred to polyvinylidence fluoride (PVDF) membranes (BioRad), blocked with 5% milk in TBS-T for 1 hr at RT (25°C) and probed with appropriate dilutions of indicated primary antibodies overnight at 4 °C Both Polyclonal Anti-SERPINA1 Antibody (HPA001292, Protein Atlas) and Anti-Beta Actin Antibody (Abcam, ab8227-50) were diluted 1:1000 in 2% BSA in TBS-T. After 3× wash in TBS-T membranes were incubated with anti-rabbit IgG conjugated with horseradish peroxidase at RT for 1 hr (1:2000 dilution) and washed 3× in TBS-T. Then Clarity Max Western ECL Substrate (Bio-Rad) was used for chemiluminescent visualization.

*hA1AT ELISA*

Levels of secreted hA1AT translated from mRNAs with different 5' cap modifications were measured in cell culture medium using A1AT Total Antigen Elisa kit (Innovative Research). Briefly, cell culture medium collected from 6-well format was diluted 5000× in Blocking Buffer (3% BSA (w/v) in 1× PBS pH 7.4). Then, diluted samples were added in duplicates to ELISA wells precoated with anti-hA1AT antibody on 96 well ELISA plate. Standard curve was prepared from hA1AT standard dilutions according to human A1AT Total Antigen Elisa kit manual in duplicates and also added to precoated wells, followed by 30 min incubation with shaking and 3× wash. In the next steps, 30 min incubation with biotin-conjugated anti-hA1AT antibody, followed by 30 min streptavidin-HRP incubation and washed steps between first and second incubation were done. In the end, substrate solution was added to the wells with samples and standard curve dilutions for 5 minutes to obtain colored product. Reaction was stopped and absorbance was measured at 450 nm. The mean values of absorbance from each samples running in duplicates was calculated in relation to absorbance values of standard curve using a four parameter logistic curve (4PL). The concentrations of hA1AT in samples were calculated by interpolation of absorbance values for each sample from standard curve using GraphPad Prism 9.0. Obtained hA1AT concentration values were shown as ng/mL.

*EnzChek® Elastase Assay*

The assay was performed according to manufacturer's protocol (ThermoFisher Scientific). To investigate the inhibition role of hA1AT released from transfected cells after mRNA transfection, 50 µl of each culture medium was used as elastase inhibitor. Also, the inhibition role of commercially available hA1AT protein was evaluated in one experiment. Next, 50 µL of 100 µg/mL DQ elastin working solution was added to plate to obtain final concentration of 25 µg/mL in each well. In the final step, the enzyme of interest - porcine pancreatic elastase – was diluted in 1× Reaction Buffer to desired final concentration and added to well to obtain final volume of 200 µL in each well. Immediately after this step, fluorescence signal was measured at multiple time points every 10 min. For each time point, the background was corrected by subtracting the value derived from the no-enzyme control. Also, no inhibitor and no substrate samples were added as negative controls.

##### 3.8. Statistical analysis

Data points represent biological replicates. Bars represent means ± standard deviation (SD), n=3. Data passed normality test using Shapiro-Wilk method. * $P<0.05$, ** $P<0.01$, *** $P<0.001$, ns – not significant, one way ANOVA with Tukey's multiple comparisons test or two-tailed unpaired *t* test, as indicated in figure captions. For all analyses differences between the groups were considered statistically significant at $P<0.05$. Statistical analysis was performed in Prism version 9 (GraphPad Software Inc., San Diego, CA).

#### 4. mRNA expression studies in human dendritic cells (hDCs)

##### 4.1. Isolation, immunophenotyping and differentiation of human CD14$^+$ cells towards dendritic cells

All donors were healthy males aged 18-60. PBMCs were isolated by density-gradient centrifugation using Lymphoprep™ (STEMCELL Technologies) from fresh buffy coats obtained from the Regional Blood Centre in Warsaw, Poland . Next, CD14$^+$ PBMCs (monocytes) were immunomagnetically separated with anti-human CD14 antibodies-coated MicroBeads using LS columns (Miltenyi Biotec) according to manufacturer's protocol. The isolated CD14$^+$ cells were differentiated towards dendritic cells (hMDCs) as follows: CD14$^+$ PBMCs were seeded into ø10 cm tissue culture Petri dishes at a density of 1-2×$10^6$ cells/mL in 10 mL of RPMI-1640 medium

(Sigma-Aldrich) supplemented with 10% heat-inactivated fetal bovine serum (FBS, HyClone), 2 mM L-glutamine (Sigma-Aldrich), 100 U/mL penicillin and 100 µg/mL streptomycin (both Sigma-Aldrich), 1% (v/v) MEM non-essential amino acids solution (ThermoFisher Scientific), 1 mM sodium pyruvate (Sigma-Aldrich). Differentiation medium was supplied with 50 ng/mL of recombinant human GM-CSF and 40 ng/mL of recombinant human IL-4 (both from Peprotech). On day 3, half of the medium was replaced and supplied with fresh cytokines. On day 6, only non-adherent and loosely adherent cells were collected for further immunophenotyping and culture. Zombie UV™ Fixable Viability Kit (BioLegend) was used for the detection of viable cells followed by blocking of unspecific binding with TruStain FcX™ (Biolegend).The antibodies mixture (CD1c/PerCp-Cy5.5: Cat. No. 331512, CD11c/PE-Cy7: Cat. No. 337216, CD83/PE: Cat. No. 305308, CD86/APC: Cat. No. 305412, HLA-DR/APC-Cy7: Cat. No. 307618, all from Biolegend) were added to the cells to detect surface markers of differentiated dendritic cells. Flow cytometry was performed using Fortessa X20 analyzer (BD Biosciences) operated by FACSDiva 8.3 software. FlowJo v10.6.1 software (BD Biosciences) was used for data analysis.

##### 4.2. *In vitro* transfection of the cells with synthetic mRNA

Transfection of cells with mRNA encoding Gluc was done using Lipofectamine MessengerMAX Reagent (ThermoFisher Scientific) according to manufacturer’s protocol. Briefly, 0.15 µL of Lipofectamine MessengerMAX Reagent was diluted in 5 µL of Opti-MEM Medium (ThermoFisher Scientific), mixed and incubated for 10 minutes at RT. Next, 5, 25 and 100 ng of mRNA was diluted in 5 µL Opti-MEM Medium and mixed well. Diluted mRNA mix was added to diluted Lipofectamine MessengerMAX™ Reagent (1:1 ratio) and incubated for 5 minutes at RT. mRNA-lipid complexes were then added to cells and incubated for designated amount of time at 37°C, 100% humidity, and 5% $CO_2$.

##### 4.3. Gluc activity detection

Gluc activity in mRNA-transfected monocyte-derived human dendritic cells was measured in culture medium every 24 h for 6 consecutive days post transfection using NanoFuel GLOW Assay (NanoLight Technology) according to the manufacturer’s manual.

##### 4.4. Statistical analysis

Data points represent biological replicates from three healthy blood donors. Bars represent means ± standard deviation (SD), n=3. Data passed normality test using Shapiro-Wilk method. * P<0.05, **P<0.01, ns – not significant two-tailed unpaired *t* test. For all analyses differences between the groups were considered statistically significant at P<0.05. Statistical analysis was performed in Prism version 9.

#### 5. Reporter mRNAs expression in vivo

##### 5.1. Mice

The experiments were carried out in 10-12-week-old (~25 g) female C57BL/6 mice, under a protocol approved by the Local Ethical Committee for Experiments on Animals in Warsaw, Poland (WAW2/063/2023) and in 10-12-week-old (~25 g) female BALB/c mice under a protocol approved by the Local Ethical Committee for Experiments on Animals in Warsaw, Poland (WAW2/126/2021). All experiments were conducted in accordance with the Directive of the European Parliament and Council No. 2010/63/EU on the protection of animals used for scientific purposes. Mice were obtained from the Breeding Facility of the Mossakowski Institute, Polish Academy of Science, Warsaw and maintained in specific pathogen-free (SPF) environment in the individually ventilated cages (IVC) under the conditions of a 12-h day/night cycle with unrestricted access to food and drinking water.

##### 5.2. mRNA formulation

Each mRNA was complexed with TransIT® transfection reagent and mRNA boost, at the mRNA/ TransIT®/mRNA boost ratio of 1:1:1 (µg mRNA: µL reagent: µL boost), according to manufacturer’s instruction (Mirus Bio).

*Lipid nanoparticle preparation in GenVoy-ILM™ mix*

The ionizable lipid mix GenVoy-ILM™ was purchased from Precision NanoSystems, Canada, and diluted 1:1 (v/v) with absolute ethanol to final concentration of 12.5 mM. All mRNAs were diluted in 100 mM sodium citrate buffer pH 4.0 at the final concentration of 120 ng/µL. The GenVoy-ILM™ and mRNA solution were combined in a microfluidic device (NanoAssemblr® Ignite™, Precision NanoSystems) equipped with NxGen Cartridge (cat#NIN0002) at a flow ratio of 3:1 (aqueous phase:ethanol) with a total flow rate of 12 mL/min. The final N/P ratio was 6, where N/P represents the ratio of ionizable nitrogen atoms to phosphate groups in the mixture.

*Lipid nanoparticle preparation with SM-102 and MC3 lipids*

The SM-102 and MC3 (DLin-MC3-DMA) lipids were purchased from BroadPharm (USA). The 1,2-dimyristoyl-rac-glycero-3-methoxypolyethylene glycol-2000 (DMG-PEG2k), cholesterol (from ovine wool) and 1,2-dioctadecanoyl-*sn*-glycero-3-phosphocholine (DSPC) were purchased from Avanti Polar Lipids (USA). The stock solutions of lipids were prepared in absolute ethanol (Thermo Fisher Scientific) at the concentration of 100 mg/mL for SM-102 and MC3 and 10 mg/mL for the rest of the lipids. The lipid mixes were prepared by combining SM-102/MC3, DSPC, cholesterol and DMG-PEG2k at molar ratio of 50:10:38.5:1.5 in absolute ethanol at total concentration of 15 mM for SM-102 lipid mix and 12.5 mM for MC3 lipid mix. Stock solution of mRNA was diluted in 100 mM sodium citrate buffer pH 4.0 at the mRNA final concentration of 102 ng/µL for SM-102 lipid mix and 120 ng/µL MC3 lipid mix. The lipid mix and mRNA solution were combined together in a microfluidic device (NanoAssemblr® Ignite™, Precision NanoSystems) equipped with NxGen Cartridge (cat#NIN0002) at a flow ratio of 4:1 with a total flow rate of 12 mL/min for SM-102 lipid mix and at a flow ratio 3:1 with a total flow rate 10 mL/min for MC3 lipid mix. The final N/P ratio was 6, where N/P represents the ratio of ionizable nitrogen atoms to phosphate groups in the mixture.

All resulting lipid nanoparticles (LNPs) were diluted 20-40 times in 1× PBS sterile buffer (w/o calcium and magnesium ions) and concentrated by ultrafiltration using Amicon ultracentrifugal tubes (Merck Millipore) with 50kDa MWCO (2000 × g, 20°C). Final LNPs were stored at 4°C and diluted in 1× PBS (w/o calcium and magnesium ions) sterile buffer before application into mice.

*LNPs size and encapsulation efficiency measurement*

Size distribution and polydispersity index were determined using dynamic light scattering (DLS) on Malvern Zetasizer Ultra Red (Malvern, UK) in PBS buffer at 25°C in back scatter mode. Encapsulation efficiency and concentration of mRNA entrapped in LNPs was determined using the Quant-iT Ribogreen RNA assay (Thermo Fisher Scientific, USA) by compering fluorescence intensities in the presence or absence of 0.1% (w/v) Triton X-100.

##### 5.3. Assessment of *in vivo* translation efficacy of Fluc-encoding mRNA-LNPs

mRNA coding for Fluc was administered intravenously (i.v.) into the lateral tail vein of BALB/c mice in different formulations. Each mouse was inoculated with 10 µg of LNP-formulated mRNA. The total body bioluminescence signal was recorded intravitally at 4 h, 8 h and 24 h post mRNA administration with IVIS Spectrum imaging system (Perkin-Elmer). Briefly, mice were administered intraperitoneally 150 mg/kg D-luciferin (Sydlabs) and anesthetized by continuous inhalation of 3% isoflurane (Baxter) mixture in 100% $O_2$. Imaging conditions were as follows: auto acquisition time set on Auto (max 1 min), F/Stop 1 and binning small. Bioluminescence values in the regions of interest (ROIs) were analyzed using Living IMAGE Software provided by Perkin-Elmer.

##### 5.4. Assessment of *in vivo* translation efficacy of hEPO-encoding mRNA-LNPs

mRNA coding for hEPO was administered i.v. into the lateral tail vein of C57BL/6 mice in different formulations. Each mouse was inoculated with 1 µg of LNP-formulated mRNA. Blood was collected at 4 h, 8 h, 24 h and 48 h time points after mRNA administration. hEPO serum concentrations were measured with ELISA (ThermoFisher Scientific).

##### 5.5. Assessment of *in vivo* translation efficacy of hA1AT

C57BL/6 mice were injected i.v. with 10 µg of A1AT-encoding $m^7GpppA_mpG$ or $m^7Gppp^{Bn6}A_mpG$-capped mRNA formulated in TransIT®. Serum A1AT concentration at 4 and 24 hrs post injection measured with ELISA.

##### 5.6. Statistical analysis

Data points represent independent biological replicates. Bars represent means ± standard deviation (SD). One-way ANOVA with Dunnett's post hoc test (***P<0.001; ****P<0.0001) or Mann-Whitney test (in the Fig. 5A and C q values are shown) were used. For all analyses differences between the groups were considered statistically significant at P<0.05. Statistical analysis was performed in Prism version 9.

#### 6. In vivo immunization experiments

##### 6.1. Mice

The experiments were carried out in 10-12-week-old (~25 g) female C57BL/6 mice under the protocols approved by the Local Ethical Committee for Experiments on Animals in Warsaw, Poland (WAW2/123/2021 and WAW2/087/2022) and in 10-12-week-old (~25 g) female BALB/c mice under a protocol approved by the Local Ethical Committee for Experiments on Animals in Warsaw, Poland (WAW2/157/2022). All experiments were conducted in accordance with the Directive of the European Parliament and Council No. 2010/63/EU on the protection of animals used for scientific purposes. Mice were obtained from the Breeding Facility of the Mossakowski Institute, Polish Academy of Science, Warsaw (C57BL/6 and BALB/c) or from the Animal Facility of the Department of Immunology, MUW (OT-I donor mice). All animals were maintained in specific pathogen-free (SPF) environment in the individually ventilated cages (IVC) under the conditions of a 12-h day/night cycle with unrestricted access to food and drinking water.

##### 6.2. mRNA formulation

Each mRNA was complexed with TransIT® transfection reagent and mRNA boost, at the mRNA/ TransIT®/mRNA boost ratio of 1:1:1 (µg mRNA: µL reagent: µL boost), according to manufacturer's instruction (Mirus Bio).

##### 6.3. OT-I T cells in vivo proliferation assay

$CD8^+$ T-cells were isolated from the spleens and lymph nodes of OT-I mice using EasySep™ Mouse $CD8^+$ T Cell Isolation Kit (Stem Cell Technologies, according to the manufacturer's manual) and labeled with Cell Trace Violet (CTV, Thermo Fisher Scientific) for 20 min at 37°C at a final concentration of 5 µM. Next, $2\times10^6$ of the CTV-stained OT-I cells in 200 µL of PBS were transferred into the caudal vein of recipient C57BL/6 mice. On the next day, mRNAs encoding OVA with different 5' mRNA cap analogues were formulated in TransIT® transfection reagent (Mirus Bio) and injected i.v. into C57BL/6 recipient mice. 72 hours later, spleens were harvested, mashed through a 70 µm nylon strainer followed by the RBC lysis using ACK Lysing Buffer (ThermoFisher Scientific). Next, splenocytes were stained with anti-CD45.2, anti-CD3ε, anti-CD8 antibodies, and H-2-Kb-SIINFEKL tetramers (see table below) and analyzed for proliferation in flow cytometry (FACSCanto II, BD Biosciences). Number of proliferating cells was calculated using the FlowJo Software v10.6.1 (Tree Star) in comparison to CountBright Absolute Counting Beads (ThermoFisher Scientific).

| **Antibodies:** | | |
|---|---|---|
| **Antigen** | **Label** | **Company** |
| CD8α | PerCP Cy5.5 | eBioscience |
| PD-1 | APC | BioLegend |
| CD25 | PE-Cy 7 | BioLegend |
| **Tetramer:** | | |
| **Class I MHC - peptide** | **Label** | **Company** |
| H-2-Kb - SIINFEKL | PE | MBL International |

##### 6.4. Obtaining of model antigen-expressing tumor cells

LLC-OVA and CT26-NY-ESO1 cells were obtained using 2nd generation lentiviral transduction. For virus production, HEK293T cells were transiently transfected with pLVX-OVA-puro or pLVX-NY-ESO1-puro plasmid together with envelope and packaging plasmids (pMG2.D and psPAX2) to generate lentiviral particles using $CaCl_2$ and 2× HBSS solution. Then, 48 h later the cell supernatant with viral particles was collected. The virus solution was purified in 0.45 mm Microcon centrifugal filters and concentrated by ultrafiltration on Amicon® Ultra-450 kDa membranes. LLC and CT26 cells were seeded 24 h before transduction. At the day of transduction, cell medium was replaced with a complete medium with polybrene (Sigma) at a final concentration of 4 µg/mL. The next day, medium from transduced cells was changed to a fresh complete medium, and 48 h post-transduction puromycin (Sigma) at a concentration of 10 µg/mL was added to cells. NY-ESO1 and OVA expression were confirmed by PCR and Western blotting.

##### 6.5. Tumor growth and treatment

At day 0 of the experiment, $3\times10^5$ LLC-OVA cells (C57BL/6 mouse) or $3\times10^5$ CT26-NY-ESO1 (BALB/c mouse) were inoculated subcutaneously into the fat pad. Mice were observed for tumor appearance and tumor growth was measured with digital calipers (AOS 100 mm with roll). Tumor dimensions - maximum length and width were measured 3 times a week, and the tumor volume was calculated according to the following formula:

$TV = ½ \times a^2 \times b$, where TV is the tumor volume, a is the shorter diameter and b is the longer diameter. To compare tumor growth-inhibiting activity between the experimental groups, tumor growth inhibition values (TGI) were estimated using the following formula:

$TGI=100-TVN/TVC\times100$, where TGI is the tumor growth inhibition value, TVN is the mean tumor volume calculated for the studied experimental group and TVC is the mean tumor volume calculated for the control group of animals.

The days of mRNA administration were determined based on the individual tumor growth kinetics in each of the experiments. mRNA for OVA or Fluc (negative control) was administered intravenously (i.v.) on days 8, 15, and 22 after inoculation of tumor cells. mRNA for NY-ESO1 was administered i.v. on days 11, 18, and 25 after inoculation of tumor cells.

##### 6.6. Statistical analysis

Data points represent independent biological replicates. Bars represent means ± standard deviation (SD). One-way ANOVA with Dunnett's post hoc test, mixed-effects analysis with Dunnett's multiple comparisons test or two-tailed unpaired *t* test were used (as indicated in the Fig. 6 caption). For all analyses differences between the groups were considered statistically significant at $P<0.05$. Statistical analysis was performed in Prism version 9.

#### 7. Biochemical and biophysical studies

##### 7.1. Recombinant proteins expression and purification

*heIF4E isoforms:* The human eukaryotic initiation factor 4E isoforms eIF4E1a and 4EHP (eIF4E2) were expressed without affinity tags in *E. Coli* Rosseta 2 (DE3)pLysS strain (Novagen) under conditions leading to accumulation of the produced target protein in inclusion bodies from which was purified as described previously.(Zuberek et al., 2007) The human eIF4E3 isoform containing the N-terminal $His_6$-Tag was expressed in *E. Coli* Rosseta 2 (DE3)pLysS strain (Novagen) under conditions leading to accumulation protein in soluble fraction and next was purified using $Ni^{2+}$ affinity (HIS-Select Nickel Affinity Gel, SIGMA) and size exclusion chromatography (ENrich SEC70, Bio-Rad) as described previously.(Zuberek & Stelmachowska, 2017) Before fluorescence binding assay proteins were filtrated using Ultrafree MC-HV Centrifugal PVDF filters with 0.45 µm membrane pore (Millipore) to remove protein aggregates.

*FTO:* Full-length human FTO (1-505) was expressed and purified similar to previous reports.(Han et al., 2010) Briefly, full-length human FTO with N-terminal 6×His-tag and TEV cleavage site was cloned into a modified pET21a bacterial expression vector, verified by whole plasmid sequencing, and transformed into E. coli BL21

Rosetta cells. Starting from 10 mL of an overnight culture, cells were grown in 1 L of LB media to $OD_{600}$ ~0.6 at 37 °C with shaking at 200 rpm. Cells were then induced with 1 mM isopropyl β-D-1-thiogalactopyranoside for 18 hours at 18°C with shaking at 200 rpm. Cells were harvested by centrifugation at 7,800 × g and cell pellets were frozen and stored at -80°C. Cell pellets were later thawed on ice and resuspended in lysis buffer (25 mM Tris-Base pH 8.0, 300 mM NaCl with Roche cOmplete EDTA-free protease inhibitor tablets), lysed by sonication, and clarified by centrifugation at 23,000 × g for 45 min. His-tagged FTO in the supernatant was purified using HisPur Ni-NTA affinity resin (ThermoFisher Scientific) and eluted with 25 mM Tris-HCl pH 7.5, 300 mM NaCl, 300 mM imidazole. The protein was buffer exchanged into 25 mM Tris pH 7.5, 25 mM NaCl and further purified by ion exchange chromatography using a 5 mL HiTrapQ column (Cytiva), followed by size exclusion chromatography using a Superdex 200pg 16/60 gel filtration column (Cytivia) in storage buffer (10 mM Hepes pH 7.5, 50 mM KCl). Fractions were analyzed using SDS-PAGE, combined and concentrated to 330 μM, flash frozen in storage buffer with 20% glycerol, and stored at -80°C.

*PNRC2-Dcp1/Dcp2:* Human decapping complex PNRC2-Dcp1/Dcp2 was expressed in Escherichia coli and purified as described previously.(Bednarczyk et al., 2022)

##### 7.2. FTO susceptibility assay

*In vitro* FTO activity assays were performed in 90 μL reaction volumes, incubating either 20 μM or 200 μM $m^7Gppp(x^6A_m)pG$ (x = methyl or benzyl) cap compound with 2 μM FTO in FTO reaction buffer (50 mM HEPES pH 7.0, 150 mM KCl, 75 μM $(NH_4)_2Fe(SO_4)_2{\cdot}6H_2O$, 300 μM 2-oxoglutarate, and 2 mM ascorbic acid). Reactions were initiated by addition of 0.55 μL of concentrated FTO aliquot to 1× substrate solution with mixing, and individual reaction timepoints were quenched by the addition of EDTA (final EDTA concentration of 1 mM). Protein was precipitated and removed from the quenched timepoint samples by the addition of 20% TCA, centrifugation, and collection of the supernatant. Analysis of cap substrate N6-methylation or benzylation status was performed on an Agilent Bio-Inert 1260 Infinity II UHPLC system with Infinity Lab LC/MSD iQ. Separation was conducted using an Agilent Poroshell 120 EC-E18 column (2.1 × 100 mm, 2.7 μm particle size) using mobile phase containing 10mM ammonium acetate (A) and 100% acetonitrile (B) and then detected by mass spectrometry in positive ionization mode. The gradient was as follows: 0-1.0 min, 100% A; 1.0-5.0 min, to 50% A/50%B; 5.0-5.5 min, 50% A/50% B; 5.5-6.0 min, to 100% B, 6.0-6.5 min, 100% B. The retention time for $m^7Gppp(Bn^6A_m)pG$ was 4.3 min with a m/z of 626.8 and the retention time for $m^7Gppp(m^6A_m)pG$ was 3.6 min with a m/z of 580.7.

##### 7.3. Determination of binding affinities for eIF4Es by fluorescence quenching titration (FQT)

Fluorescence titration measurements were carried out on a LS-55 spectrofluorometer (Perkin Elmer Co., Norwalk, CT., USA), in 50 mM HEPES/KOH (pH 7.2), 0.5 mM EDTA, 100 mM KCl, at 20.0 ± 0.2°C using 0.1 μM concentration of proteins. The eIF4Es fluorescence was excited at 280 nm and the fluorescence intensity was monitored at a single wavelength 340 nm. The measured fluorescence intensities were corrected for dilution and for the inner filter effect. The equilibrium association constants ($K_{AS}$) were obtained by fitting a theoretical dependence of the fluorescence intensity on the total concentration of cap analogue to the experimental data points, according to equation described previously.(Niedzwiecka et al., 2002) The final $K_{AS}$ values were calculated as a weighted average from three independent titrations and represented as the $K_D$ values in **Table 1**. Numerical least-squares nonlinear regression analysis was performed using ORIGIN 6.0 from Microcal Software Inc., USA.

##### 7.4. mRNA translation in rabbit reticulocyte lysate (RRL)

An RRL System (Promega) was used to compare translation efficiency of Fluc-coding mRNA capped with $m^7GpppA_mpG$, $m_7Gppp^{m6}A_mpG$ or $m^7Gppp^{Bn6}A_mpG$. The initial translation mix (8 μL) was incubated for 1 h at 30°C, and contained: 4 μL RRL, 20 μM amino acid mixture without leucine, 20 μM amino acid mixture without methionine, 190 mM potassium acetate, and 1 mM magnesium acetate. Next, 2 μL of 0.5 ng/μL RP-HPLC-purified Fluc-coding mRNA was added to 8 μL of initial translation mix and further incubated for 1 h at 30°C. The reaction was stopped by freezing in liquid nitrogen.

Translation efficiency was determined as a correlation of the Fluc activity and measured luminescence emitted during luciferin oxidation by the enzyme. To detect luminescence 50 µL of the Luciferase Assay Reagent (tricine, $Mg(HCO_3)_2$, $MgSO_4$, 0.1 mM EDTA, DTT, ATP, luciferin (VivoGlo™ Luciferin, *In Vivo* Grade; Promega) and coenzyme A) was added to 10 µL of the translation mix diluted 2× with $H_2O$ prior luminescence measurement performed on a Synergy H1 (BioTek) microplate reader.

##### 7.5. Dcp1/Dcp2 susceptibility assay

#### 8. 20 ng of capped RNA (see 2. mRNA synthesis, purification, and quality control

##### 8.1. Preparation of DNA plasmid vectors

Luciferase gene (Fluc) from firefly (Lampyridae) was purchased from Invitrogen (ThermoFisher Scientific) with restriction sites for AdeI and BamHI endonucleases, and cloned into pJet1.2 (ThermoFisher Scientific) plasmid vector using sticky-end cloning method. The ordered plasmid with Fluc sequence (2 µg) was digested (15 min, 37°C) with AdeI (ThermoFisher Scientific) and BamHI (ThermoFisher Scientific) restriction enzymes and 10× Fast Digest Buffer (ThermoFisher Scientific). The digested DNA was then separated on an 1% agarose gel and the strand containing the Fluc sequence was excised and purified using a commercial DNA purification kit (Macherey-Nagel) according to the protocol. The pJet1.2 plasmid vector (2 µg) was linearized (16 h, 37°C) with AarI restriction enzyme (ThermoFisher Scientific), 10× AarI buffer (ThermoFisher Scientific) and 50× Oligo (ThermoFisher Scientific) and purified using a commercial DNA purification kit (Macherey-Nagel). Then the pJET1.2 solution including 100 ng of DNA was mixed with the insert (Fluc sequence) at a molar ratio of 1:3 (vector- insert), added 10× Ligase buffer (ThermoFisher Scientific) and T4 DNA Ligase (ThermoFisher Scientific), incubated (1 h, 25°C), and then reaction mix (20 µl) cooled to 4°C and incubated an additional 1 h. Half of the ligation mixture (10 µl) was mixed with pre-melted on ice commercially available chemo competent bacteria (50 µl) Top10 (ThermoFisher Scientific) and incubated on ice for 30 min. The mixture was transferred to 42°C (heat shock) and cooled on ice for 2 min and 500 µl of SOC outgrowth medium (ThermoFisher Scientific) was added and incubated at 37°C for 1 h with shaking (300 RPM). The transformation mixture (200 µl) was spread on LB-agar plate (Roth) with 100 mg/mL Ampicillin (Roth) and incubated (37°C, 16 h). Then, single colonies were selected and inoculated with liquid LB medium (Roth) in a volume of 5 mL and supplemented with Ampicillin (Roth) at a concentration of 100 mg/mL and the cultures were incubated (37°C, 16 h). The bacterial cultures were then centrifuged (4000g, 10 min) and the plasmids were purified using the commercial GeneJET Plasmid Miniprep Kit (ThermoFisher Scientific). Concentration was measured using a Nanodrop 2000c spectrophotometer (ThermoFisher Scientific) and plasmids were sent for DNA sequencing using the Sanger method (Genomed). The pJet1.2_FLuc plasmid contained a short form of the poly(A) tail ~30 adenine nucleotide. In order to obtain a DNA template with a poly(A) tail consist of 90 adenine nucleotides, a several-step procedure was carried out to insert an adenine oligonucleotide into the 3′ end of Fluc sequence. The insert was designed as a double-stranded DNA oligonucleotide and added to the pJet plasmid vector that encodes Fluc using a blunt end cloning method. Solutions of two DNA oligonucleotides (Genomed) with sequence: $A_{60}$ (coding strand); $T_{60}$ (template strand) were mixed in a 1:1 ratio (final 50 µM of each DNA strand) and an enzyme reaction was set up to phosphorylate the 5' ends of the oligonucleotides by adding T4 Polynucleotide Kinase (NEB) and 10× buffer for T4 Polynucleotide Kinase (NEB). The reaction was carried out for 30 minutes at 37°C. The strands were then hybridized by heating to 95°C and slowly cooling to 25°C for 2 h, step gradient ~2°C/~3 min. The insert DNA was purified using a commercial DNA purification kit (Macherey-Nagel) according to the protocol. The circular plasmid pJET1.2_Fluc that encodes firefly luciferase (2 µg) was linearized (16 h, 37°C) with AarI restriction enzyme (ThermoFisher Scientific), 10× AarI buffer (ThermoFisher Scientific) and 50× Oligo (ThermoFisher Scientific) and purified using a commercial DNA purification kit (Macherey-Nagel). The enzymatic reaction was then prepared with DNA Polymerase I, Large Fragment (Klenow), 10× buffer 3 (NEB), and 10 mM NTP (ThermoFisher Scientific) to remove 3′ overhangs and filling in 5′ overhangs to form blunt ends (15 min, 25°C) and again purified the vector using a commercial DNA purification kit (Macherey-Nagel). Finally, dephosphorylation reaction was performed using FastAP Thermosensitive Alkaline Phosphatase (ThermoFisher Scientific) and 10× Fast AP buffer (ThermoFisher Scientific) by incubating (10 min, 37°C) and purifying the final vector using a commercial DNA purification kit

(Macherey-Nagel). The final vector pJET1.2_Fluc solution including 100 ng of DNA was mixed with the insert at a molar ratio of 1:3 (vector- insert), added 10× Ligase buffer (ThermoFisher Scientific), PEG 2000 (ThermoFisher Scientific), ATP (ThermoFisher Scientific) and T4 DNA Ligase (ThermoFisher Scientific) and incubated (1 h, 25°C) and then reaction mix (20 µl) cooled to 4°C and incubated an additional 1 h. Then transformation and clonal selection as well as DNA sequencing were carried out analogous to the procedure described above.

Human erythropoietin gene (hEPO), *Gaussia* luciferase gene (Gluc) and ovalbumin (OVA) were purchased from Invitrogen (ThermoFisher Scientific) with restriction sites for AdeI and BamHI endonucleases, and was cloned (analogously to the method for Fluc) into pJet1.2 (ThermoFisher Scientific) plasmid vector using sticky-end cloning method. Then, an analogous procedure to that described above was carried out to extend the poly(A) tail for the pJet1.2_EPO, pJet1.2_Gluc and pJet1.2_OVA plasmids to the final poly(A) tail length of 90 adenine nucleotides. α1-antitrypsin gene (hA1AT) and NY-ESO1 gene were purchased from Invitrogen (ThermoFisher Scientific) in pUC57 plasmid vector. Then, an analogous procedure to that described above was carried out to extend the poly(A) tail for the pUC57_A1AT and pUC57_NY-ESO1 plasmids to the final poly(A) tail length of 90 adenine nucleotides.

##### 8.2. Capping efficiency of short RNAs

Short RNAs were obtained by IVT using T7 class II promoter Φ2.5 (initiated by ATP – TAATACGACTCACTATT**A**) or class III promoter Φ6.5 (initiated by GTP – TAATACGACTCACTATA**G**). The typical IVT reaction (100 µL) was incubated for 4 h at 37°C and contained: 3 mM each of UTP/GTP/CTP for Φ2.5 or UTP/ATP/CTP for Φ6.5, 0.75 mM of ATP or GTP (NTPs were purchased from ThermoFisher Scientific Scientfic), respectively, and 6 mM of trinucleotide cap analog ($m^7GpppA_mpG$, $m^7Gppp^{m6}A_mpG$ or $m^7Gppp^{Bn6}A_mpG$), 1.25 µM of annealed oligonucleotides as DNA template, 1 U/µL RiboLock RNase Inhibitor (ThermoFisher Scientific), 20 mM $MgCl_2$, RNA Polymerase Buffer (ThermoFisher Scientific) and T7 RNA polymerase (0.3 mg/mL, in-house prepared). The preparation of uncapped RNAs was performed as before, but with 3 mM of ATP/GTP and without the presence of cap analog. After 4 hours sample was treated with DNase I (5 U, ThermoFisher Scientific) and incubated for another 30 min at 37°C. The reaction mixture was stopped by addition of equimolar amount of $Na_2EDTA$ water solution (EDTA to $Mg^{2+}$) and purified using Monarch® RNA Cleanup Kit (NEB). Such prepared RNA sample was then purified using HPLC with Phenomenex® Clarity 3 µM Oligo-RP column with a linear gradient of buffer B (0.2 M triethylammonium acetate pH 7.0 and acetonitrile, v/v) from 10% to 33.3% in buffer A (0.1 M triethylammonium acetate pH 7.0) in 35 min at 1 mL/min. Collected fractions of RNA were then precipitated with 0.33 M NaOAc pH 5.2 and 2.5 volumes of 80% ethanol. Moreover, to obtain homogenous 3´ end of synthetized RNAs, obtained 35-nt long uncapped and 36- or 37-nt long capped transcripts (1 µM) were incubated with 1 µM DNAzyme 10-23 (TGATCGGCTAGGCTAGCTACAACGAGGCTGGCCGC) in 50 mM Tris pH 8.0 and 50 mM $MgCl_2$ for 1 hour at 37°C, purified using HPLC and precipitated as above. The quality of obtained short RNAs and capping efficiency were checked on 15% acrylamide / 7 M urea / 1× TBE gel with SYBR Gold (Invitrogen) staining and the concentration was determined spectrophotometrically using NanoDrop 2000c. The PAGE band intensities corresponding to capped and uncapped RNAs were quantified densitometrically using 1D Gel Image Analysis Software TotalLab CLIQS.

##### 8.3. Optimization of in vitro transcription conditions for improvement of capping efficiency with $m^7Gppp^{Bn6}A_mpG$

The conditions of the IVT have been optimized to achieve ~90% capping efficiency with $m^7Gppp^{Bn6}A_mpG$. For these studies DNA template encoding ~1000 nt long transcript (NY-ESO1) was chosen to ensure sufficient separation of capped and uncapped mRNA using analytical RP-HPLC.

In the first set of optimization experiments capping efficiency was determined for mRNA obtained in IVT at various pH, ranging from 6.0 to 9.0. 20 µL of each prepared IVT mix contained following components at indicated final concentration: transcription buffer prepared in two sets of variants: 1) containing 40 mM Bis-Tris at the pH ranging from 6.0 to 7.0, and 2) containing 40 mM Tris-HCl at the pH ranging from 7.0 to 9.0, 2 mM spermidine, 1 U/µL RNase inhibitor (RiboLock, ThermoFisher Scientific), 5 mM ATP, CTP and UTP, 4 mM GTP, 10 mM DTT, 25 mM $MgCl_2$, 0.002 U/µL inorganic pyrophosphatase (ThermoFisher Scientific), 10 mM $m^7Gppp^{Bn6}A_mpG$, 40 ng/µL linearized plasmid as the DNA template and 0.125 µg/µL T7 RNA polymerase (in-house prepared). All components were gently mixed by pipetting and incubated for one hour at 37°C. To remove DNA template, IVT mix was incubated for next 30 min at 37°C with 0.025 U/µL DNase I (ThermoFisher Scientific). The enzymes in the IVT mix were inactivated by addition of one volume of 52 mM EDTA, and full-length mRNAs were isolated from abortive transcripts using oligo dT chromatography (POROS™ Oligo $(dT)_{25}$ Affinity Resin, ThermoFisher Scientific) according to manufacturer's protocol. In short, crude IVT mix was diluted with high salt buffer (500 mM NaCl, 10 mM Tris-HCl pH 7.5, 1 mM EDTA) and loaded on the accordingly equilibrated oligo$(dT)_{25}$ resin. After binding step (10 min, RT), resin was washed with high and low salt buffer (200 mM NaCl, 10 mM Tris-HCl pH 7.5, 1 mM EDTA), respectively. mRNA was eluted with RNase-free water heated to 65°C. At this step IVT efficiency was estimated by measuring the volume and the absorbance of eluted mRNA at 260 nm (NanoDrop One$^C$, ThermoFisher Scientific) and calculated as mRNA yield obtained from 1 µL of starting IVT volume.

Capping efficiency was determined using RP-HPLC method based on delayed retention of mRNA containing $m^7Gppp^{Bn6}A_mpG$ in a linear gradient of organic solvent. Agilent Infinity 1260 II instrument and RNASep™ Prep (ADS Biotec) was used for analysis of 1 µg of each mRNA sample initially purified by oligo$(dT)_{25}$. Sufficient separation of capped and uncapped mRNA was obtained for a linear gradient of acetonitrile (10-14.5% in 22 min at 0.9 mL/min flow rate) in 0.1 M triethylammonium acetate (TEAA) pH 7.0 at 55°C. Capping efficiency was determined based on the area of automatically integrated peaks (using Agilent OpenLAB CDS ChemStation Edition C.01.10), referring each of the peak to capped and uncapped mRNA. To confirm results obtained using RP-HPLC, capping efficiency was also determined by ribozyme-based assay (see *2.7 Capping efficiency of full-length mRNAs*).

Second set of experiments was performed for the buffers at pH in the range of 6.0-7.0 with 0.25 intervals. mRNA yield was estimated as described previously, and for RP-HPLC analysis bioZen™ 2.6 µm Oligo LC column (Phenomenex®) was used at the linear gradient of acetonitrile (10-20% in 50 min at 0.5 mL/min flow rate) in 0.1 M TEAA pH 7.0 at 55°C.

##### 8.4. In vitro transcription protocols

Depending on the application, mRNA was *in vitro* transcribed and purified following one of optimized IVT protocols:

**Protocol A**

IVT mix contained following components at indicated final concentration: 40 mM Tris-HCl pH 7.9, 2 mM spermidine, 1 U/µL RNase inhibitor (RiboLock, ThermoFisher Scientific), 2 mM ATP, CTP and UTP, 1 mM GTP, 1 mM DTT, 10 mM $MgCl_2$, 0.002 U/µL inorganic pyrophosphatase (ThermoFisher Scientific), 2 mM $m^7GpppA_mpG$ or $m^7Gppp^{Bn6}A_mpG$, 50 ng/µL linearized plasmid as the DNA template and 0.125 µg/µL T7 RNA polymerase (*in-house* prepared). All components were gently mixed by pipetting and incubated for 2 h at 37°C. After DNA template removal (incubation with 0.025 U DNase I/µL IVT for 30 min at 37°C) the crude mRNA sample was initially purified using Monarch® RNA Cleanup Kit (New England Biolabs).

Next, uncapped transcripts were enzymatically removed from the sample by two-step reaction. First, initially purified mRNA was treated with 5′ polyphosphatase (Lucigen), which hydrolizes 5′ triphosphate to 5′ monophosphate. Typical reaction mix contained: 30 µg of mRNA, 2 U/µL RNA 5′ polyphosphatase, 1× reaction buffer (50 mM HEPES-KOH pH 7.5, 100 mM NaCl, 1 mM EDTA, 0.1% β-mercaptoethanol, 0.01% Triton X-100) and 1 U/µL RNase inhibitor (RiboLock, ThermoFisher Scientific). After 1 h incubation at 37°C, each sample was purified using Monarch® RNA Cleanup Kit (New England Biolabs). In the second step uncapped mRNA molecules with 5′ monophosphate were degraded by XRN-1 5′→3′ exonuclease (New England Biolabs). The reaction mix contained: 30 µg of mRNA, 0.5 U/µL XRN-1, 1× reaction buffer (50 mM Tris-HCl pH 7.9, 100 mM NaCl, 10 mM $MgCl_2$, 1 mM DTT) and 1 U/µL RNase inhibitor (RiboLock, ThermoFisher Scientific). After 2 h incubation at 37°C, each sample was purified using Monarch® RNA Cleanup Kit (New England Biolabs).

To remove contaminants, such as dsRNA and short mRNA species, second purification step was applied. Each mRNA sample was purified by RP-HPLC on Shimadzu Nexera Prep using RNASep™ Prep column (ADS Biotec). mRNA was eluted with the linear gradient (10-14.5%) of acetonitrile in 0.1 M TEAA pH 7.0 at 55°C. Eluate quality was analyzed on 1× TBE 1.2% agarose gel, and the fractions containing mRNA of the highest purity were combined, precipitated overnight at -20°C with 0.3 M NaOAc pH 5.2 and one volume of isopropanol, and washed with 80% ethanol. Prior further analysis, mRNA pellets were resuspended in RNase-free water.

**Protocol B**

IVT mix contained following components at indicated final concentration: 40 mM Tris-HCl pH 7.9, 2 mM spermidine, 1 U/µL RNase inhibitor (RiboLock, ThermoFisher Scientific), 5 mM ATP, CTP and UTP, 4 mM GTP, 10 mM DTT, 25 mM $MgCl_2$, 0.002 U/µL inorganic pyrophosphatase (ThermoFisher Scientific), 10 mM $m^7GpppA_mpG$ or $m^7Gppp^{Bn6}A_mpG$, 40 ng/µL linearized plasmid as the DNA template and 0.125 µg/µL T7 RNA polymerase (*in-house* prepared). All components were gently mixed by pipetting and incubated for 1 h at 37°C. After DNA template removal (incubation with 0.025 U DNase I/µL IVT for 30 min at 37°C), enzymes in IVT mix were inactivated by addition of one volume of 52 mM EDTA, and the crude mRNA sample was initially purified using oligo$(dT)_{25}$ resin (as previously described) and concentrated by ultrafiltration (Amicon Ultra-15 100K, Millipore) prior loading on RP-HPLC column.

To remove contaminants, such as dsRNA and short mRNA species, second purification step was applied. Each mRNA sample was purified by RP-HPLC on an Agilent Infinity 1260 II using RNASep™ Prep or RNASep™ Semi-Prep column (ADS Biotec). mRNA was eluted with the linear gradient (10-14.5%) of acetonitrile in 0.1 M TEAA pH 7.0 at 55°C. Eluate quality was analyzed on 1× TBE 1.2% agarose gel, and the fractions containing mRNA of the highest purity were combined, desalted by ultrafiltration (Amicon Ultra-15 100K, Millipore), precipitated overnight at -20°C with 0.3 M NaOAc pH 5.2 and one volume of isopropanol, and washed with 80% ethanol. Prior further analysis, mRNA pellets were resuspended in RNase-free water.

**Protocol C**

IVT mix contained following components at indicated final concentration: 40 mM Tris-HCl pH 7.9 (for IVT with $m^7GpppA_mpG$) or 40 mM Bis-Tris pH 6.5 (for IVT with $m^7Gppp^{m6}A_mpG$ or $m^7Gppp^{Bn6}A_mpG$), 2 mM spermidine, 1 U/µL RNase inhibitor (RiboLock, ThermoFisher Scientific), 5 mM ATP, CTP and UTP, 4 mM GTP, 10 mM DTT, $MgCl_2$ (10 mM for IVT with $m^7GpppA_mpG$; 25 mM for IVT with $m^7Gppp^{m6}A_mpG$ or $m^7Gppp^{Bn6}A_mpG$), 0.002 U/µL inorganic pyrophosphatase (ThermoFisher Scientific), cap analog (10 mM $m^7GpppA_mpG$ or $m^7Gppp^{Bn6}A_mpG$; 12 mM $m^7Gppp^{m6}A_mpG$), 40 ng/µL linearized plasmid as the DNA template and 0.125 µg/µL T7 RNA polymerase (*in-house* prepared). All components were gently mixed by pipetting and incubated for 1 h at 37°C. After DNA template removal (incubation with 0.025 U DNase I/µL IVT for 30 min at 37°C), enzymes in IVT mix were inactivated by addition of one volume of 52 mM EDTA, and the crude mRNA sample was initially purified using oligo$(dT)_{25}$ resin (as previously described) and concentrated by ultrafiltration (Amicon Ultra-15 50K or 100K, Millipore) prior loading on preparative column.

To remove contaminants, such as dsRNA and short mRNA species, second purification step was applied. Each mRNA sample was purified by RP-HPLC on an Agilent Infinity 1260 II using RNASep™ Semi-Prep columns (ADS Biotec). mRNA was eluted with the linear gradient (10-14.5%) of acetonitrile in 0.1 M TEAA pH 7.0 at 55°C. Eluate quality was analyzed on 1× TBE 1.2% agarose gel, and the fractions containing mRNA of the highest purity were combined, desalted by ultrafiltration (Amicon Ultra-15 50K or 100K, Millipore), precipitated overnight at -20°C with 0.3 M NaOAc pH 5.2 and one volume of isopropanol, and washed with 80% ethanol. Prior formulation and quality control, mRNA pellets were resuspended in RNase-free water to obtain concentration of >1 µg/µL and – if necessary – stored at -80°C as single-use aliquots.

##### 8.5. RP-HPLC analysis of mRNA purity and capping efficiency

Quality of the mRNA samples after purification process and capping efficiency for mRNAs with $m^7Gppp^{Bn6}A_mpG$ were verified by RP-HPLC using Agilent Infinity II 1260 instrument and RNASep™ Prep 7.8 × 50 mm (ADS Biotec) or bioZen™ 2.6 µm Oligo LC column 100 × 2.1 mm (Phenomenex®). 1 µg of each mRNA sample was analysed by applying linear gradient of acetonitrile (for RNASep™ Prep column: 10-14.5%

in 22 min at 0.9 mL/min flow rate; for bioZen™ column: 10-20% in 50 min at 0.5 mL/min) in 0.1 M triethylammonium acetate (TEAA) pH 7.0 at 55°C. Capping efficiency was determined based on the area of automatically integrated peaks (using Agilent OpenLAB CDS ChemStation Edition C.01.10), referring each of the peak to capped and uncapped mRNA.

##### 8.6. Analysis of double-stranded RNA content

Double-stranded RNA (dsRNA) impurities in mRNA samples were detected and semi-quantified using dot-blot assay. For each sample dsRNA presence was determined 1) after IVT and initial purification using oligo$(dT)_{25}$ resin (to check the influence of cap analog used for IVT on the dsRNA formation) or 2) after purification using RP-HPLC or cellulose (to verify purity of the sample prior mRNA application). To detect even traces of dsRNA impurities in the samples dedicated for in vivo experiments, analyzed portions contained 250 and 2500 ng of mRNA purified by RP-HPLC or cellulose. Analyzed amounts of the samples further used in the in vitro experiments were reduced to 25 and 250 ng. To avoid signal saturation, samples analyzed after initial purification using oligo$(dT)_{25}$ resin contained 5 and 25 ng of mRNA.

For each analysis intended amounts of mRNA were diluted to obtain final volume of 25 – 50 µL, which was further blotted on the Amersham™ Hybond™-N$^{+}$ membrane (GE Healthcare) soaked with 1× TBST using Bio-Dot microfiltration apparatus (Bio-Rad). Membrane with UV cross-linked mRNA was blocked for 1 h at RT with 5% non-fat dried milk in 1× TBST buffer and subsequently incubated overnight at 4°C with monoclonal mouse anti-dsRNA J2 antibody (SCICONS) diluted 1:5000 with 1× TBST buffer containing 2% non-fat dried milk. After incubation with primary antibody the membrane was washed 3 × 5 min with 1× TBST at RT and incubated for 1 h at RT with polyclonal anti-mouse IgG secondary antibody conjugated with horseradish peroxidase (Cell Sygnaling Technology) diluted 1:3000 with 1× TBST buffer containing 2% non-fat dried milk. Then, the membrane washed 3 × 5 min with 1× TBST at RT was incubated with Amersham™ ECL™ Prime Western Blotting Detection Reagent (Cytiva), and the chemiluminescence signal was detected on Amersham™ ImageQuant 800 (GE Healthcare) and quantified using ImageQuantTL software (GE Healthcare), by the comparison to dsRNA standard in the range of 0.078-10 ng (Abnova), 0.062-8 ng, 0.156-20 ng or 0.391-50 ng (New England Biolabs), prepared separately for each membrane.

##### 8.7. Capping efficiency of full-length mRNAs

Capping efficiency could differ depending on nucleotide sequence of the template, therefore efficiency of $m^{7}GpppA_{m}pG$ and $m^{7}Gppp^{Bn6}A_{m}pG$ cap analogs incorporation was determined individually for the most of the transcripts.

For that purpose, ribozyme complementary to the 5´UTR sequence was designed and used for cleavage of mRNA close to the 5´ end. 10 µL of analyzed sample containing >10 µg (>100 nM) of mRNA was mixed with 1.5 µL of hybridization buffer containing 100 mM Tris-HCl pH 7.5 and 50 mM NaCl and 2 µL of 10 µM ribozyme. The mixture was incubated at 95°C for 2 min, and subsequently at RT for 10-15 min. Cleavage reaction with the ribozyme hybridized to mRNA was initiated by addition of 1.5 µL of 100 mM $MgCl_2$, conducted at 37°C for 1 h, and quenched by addition of 2 µL 100 mM EDTA.

To analyze cleaved RNA fragments 3 µL of quenched reaction was mixed with equal volume of loading dye (8 M urea, 50% formamide, 20 mM EDTA, 0.03% bromophenol blue, 0.03% xylene cyanol), heat denatured at 95°C for 3 min and loaded onto 15% polyacrylamide gel with 7 M urea and 1× TBE. The gel after electrophoresis was incubated at RT for 15 min with 50 mL of staining reagent (SYBR® Gold, Invitrogen, diluted 1:10000), and visualized using Typhoon scanner (GE Healthcare). Migration of short RNA fragments being the result of ribozyme cleavage differs depending on the cap structure present on the 5´ end of mRNA, and the bands intensities correlate with the amount of the RNA. Therefore, the intensity of the bands corresponding with $m^{7}GpppA_{m}pG$-RNA, $m^{7}Gppp^{Bn6}A_{m}pG$-RNA and pppG-RNA were quantified densitometrically using ImageQuantTL software (GE Healthcare). Percentage of capped mRNA in each sample was determined as the ratio of $m^{7}GpppA_{m}pG$-RNA or $m^{7}Gppp^{Bn6}A_{m}pG$-RNA and the summarized intensities for capped RNA and pppG-RNA.

) was subjected to digestion with 11 nM PNRC2-Dcp1/Dcp2 in 50 mM Tris·HCl pH=8.0, 50 mM $NH_4Cl$, 0.01% Igepal, 5 mM $MgCl_2$ and 1 mM DTT. Reactions were carried out at 37°C and at indicated time terminated by adding equal volume of loading dye (4.5 M urea, 50% formamide, 20 mM EDTA, 0.03% bromophenol blue, 0.03% xylene cyanol) and flash freezed. Samples were then resolved by PAGE on denaturing 15% acrylamide / 7 M urea / 1× TBE gel and were stained with SYBR Gold (Invitrogen) and visualized using a Typhoon FLA 9500 (GE Healthcare). The band intensities corresponding to capped and uncapped RNAs were quantified densitometrically using 1D Gel Image Analysis Software TotalLab CLIQS.

##### 8.8. mRNA stability measurements by RT-qPCR

Cells (HEK293T) were harvested using standard procedure with trypsin, washed in OPTIMEM and counted. 4 µg of mRNA for Gluc was transferred to chilled electroporation cuvette and mixed with 200 µl of cell suspension containing $4\times10^6$ cells. The cuvettes were then incubated on ice for 5 min. The electroporation took place in a Gene Pulser Xcell (BioRad) - square wave 25 ms, 110 V. Immediately after electroporation 800 µl of DMEM medium was added to cuvette and solution was incubated for 10 min on ice. Next cells were seeded on 6 well plate and incubated with indicated time. In order to isolate RNA, cells were suspended in Trizol reagent immediately after electroporation procedure and at 1, 2, 4 and 8 hours post electroporation. Briefly, the further steps included (centrifugation in 4°C in-between each step) a) chloroform addition and upper phase collection, b) isopropanol addition, c) 75% ethanol addition. Finally, pellet was left to dry and RNA was resuspended in water. 1 µg RNA was used for reverse transcription reaction and all further steps were according to protocol (High-Capacity cDNA Reverse Transcription Kit with RNase Inhibitor, Applied Biosystems) with one modification – 0.1 µg oligo(dT) per reaction was also added. Afterwards, qPCR reaction was performed in LightCycler® 480 (Roche). Mastermix was prepared according to manufacturer protocol (SG qPCR MasterMix, EurX) and qPCR reaction was set as follows: 50°C / 2min, preincubation: 95°C / 10min, 45 cycles: 94°C / 15sec, 59°C / 30sec, 72°C / 30sec. Primers for Gluc and β-actin were used in qPCR: B-Actinf1: GCCGGGACCTGACTGACTAC, B-Actinr1: TTCTCCTTAATGTCACGCACGAT, Gluc_f1: AAGACTTCAACATCGTGGCCG, Gluc_r1 GCTTCCAACTCTTTGAGCACC.

##### 8.9. Pull-down assay

The protein extract from HEK293F cells was prepared according to the published protocol (Mukherjee, Fritz, Kilpatrick, Gao, & Wilusz, 2004). The resins (100 µL of the settled resin) were loaded into 2 mL columns equipped with a filter and washed first with water (3 × 2 mL) and then with buffer A (50 mM HEPES pH 8.0, 100 mM KCl, 0.5 mM EDTA; 3 × 2 mL). The outlets of the columns were closed and 1 mL of protein extract from HEK293F cells (10.7 mg/mL stocks diluted 5-times with buffer A) with 200 µM GTP was added. The columns were tightly closed and shaken gently at 4°C overnight. The resins were washed with buffer A (5 × 2 mL) and the proteins were eluted with buffer B (buffer A + 200 µM cap analog: $m^7GpppA_mpG$ for **AR-1**, $m^7Gppp^{m6}A_mpG$ for **AR-2**, or $m^7Gppp^{Bn6}A_mpG$ for **AR-3**; 2 × 200 µL).

To the solutions of proteins eluted from the beads, TCEP (10 mM) and chloroacetamide (15 mM) were added, and the mixtures were subjected to overnight enzymatic digestion (0.5 µg, Sequencing Grade Modified Trypsin, Promega) at 37°C. Tryptic peptides were then desalted with the use of AttractSPE™ Disks Bio C18 (Affinisep) and TMT-labelled on the solid support (Myers et al., 2019). Individual TMT-labelled samples (3 replicates of **AR-1**, 3 replicates of **AR-2**, 3 replicates of **AR-3**) were compiled into a TMT-9 dataset and concentrated using a SpeedVac concentrator. Prior to LC-MS measurement, the sample was resuspended in 0.1% TFA, 2% acetonitrile in water. Chromatographic separation was performed on an Easy-Spray Acclaim PepMap column 50 cm long × 75 µm inner diameter (ThermoFisher Scientific) at 55°C by applying a 120 min acetonitrile gradient in 0.1% aqueous formic acid at a flow rate of 300 nL/min. An UltiMate 3000 nano-LC system was coupled to a Q Exactive HF-X mass spectrometer via an easy-spray source (all ThermoFisher Scientific). The spectrometer was operated in TMT mode with survey scans acquired at a resolution of 60,000 at m/z 200. Up to 15 of the most abundant isotope patterns with charges 2–5 from the survey scan were selected with an isolation window of 0.7 m/z and fragmented by higher-energy collision dissociation (HCD) with normalized collision energies of 32, while the dynamic exclusion was set to 35 s. The maximum ion injection times for the survey scan and the MS/MS scans (acquired with a resolution of 45,000 at m/z 200) were 50 and 96 ms, respectively. The ion target value for MS was set to 3e6 and for MS/MS to 1e5, and the minimum AGC target was set to 1e3. The data were processed with MaxQuant v. 1.6.17.0 (Cox & Mann, 2008), and the peptides were identified from the MS/MS spectra searched against the reference human proteome UP000005640 (Uniprot.org) using the built-in Andromeda search engine. Cysteine carbamidomethylation was set as a fixed modification, and methionine oxidation and protein N-terminal acetylation were set as variable modifications. For in silico digests of the reference proteome, cleavages of arginine or lysine followed by any amino acid were allowed (trypsin/P), and up to two missed cleavages were allowed. Reporter ion MS2 quantification was performed with the min. reporter PIF was set to 0.75. The FDR was set to 0.01 for peptides, proteins and sites. The second peptide search was disabled. Other parameters were used as pre-set in the software. Unique and razor peptides were used for quantification enabling protein grouping (razor peptides are the peptides uniquely assigned to protein groups and not to individual proteins). Data were further analyzed using Perseus version 1.6.10.0 (Tyanova et al., 2016), and Microsoft Office Excel 2016. Reporter intensity corrected values for protein groups were loaded. Standard filtering steps were applied to clean up the dataset: reverse (matched to decoy database), only identified by site, and potential contaminants (from a list of commonly occurring contaminants included in MaxQuant) protein groups were removed. Reporter intensity values were Log2 transformed and normalized by median subtraction within TMT channels. 1247 Protein groups identified by at least 2 razor peptides and with a complete set of 9 values were subjected to further analysis. Two-sided T-tests (permutation-based FDR=0.005, $S_0$=1) were performed for 3 comparisons of sample groups: **AR-3** vs **AR-1**; **AR-2** vs **AR-1**; **AR-3** vs **AR-2** to identify proteins differently bound to the resins under investigation. The raw LC-MS/MS data and the output from MaxQuant have been deposited to the ProteomeXchange Consortium ref via the PRIDE ref partner repository with the dataset identifier PXD046838.

##### 8.10. Cells treatment with INK128

Human monocyte-derived dendritic cells (5 x $10^4$) were pretreated for 4 h with 10 µM INK128 (mTOR inhibitor blocking eIF4E-dependent mRNA translation initiation) or DMSO at concentration corresponding to 10 µM INK128 (no INK128). INK128 (MedChemExpress Cat. No. HY-13328), was dissolved in DMSO to make 100uM stock. Next, the cells were transfected with 50 ng of $m^7GpppA_mpG$ or $m^7Gppp^{Bn6}A_mpG$-capped mRNA for hEPO using LipotectAMINE Messenger Max. hEPO concentration in culture medium 6, 24 and 48 h post transfection measured with ELISA.

#### 9. Compounds characterization

<table>
<tr><th colspan="2">(1) m<sup>7</sup>Gppp<sup>Bn6</sup>A<sub>m</sub>pG</th></tr>
<tr><td>Chemical structure</td><td>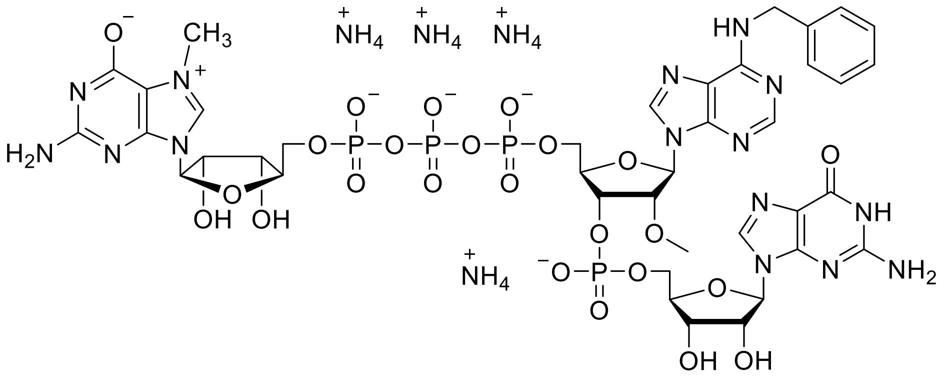</td></tr>
<tr><td>RP HPLC<br>Abs. @ 254 nm</td><td>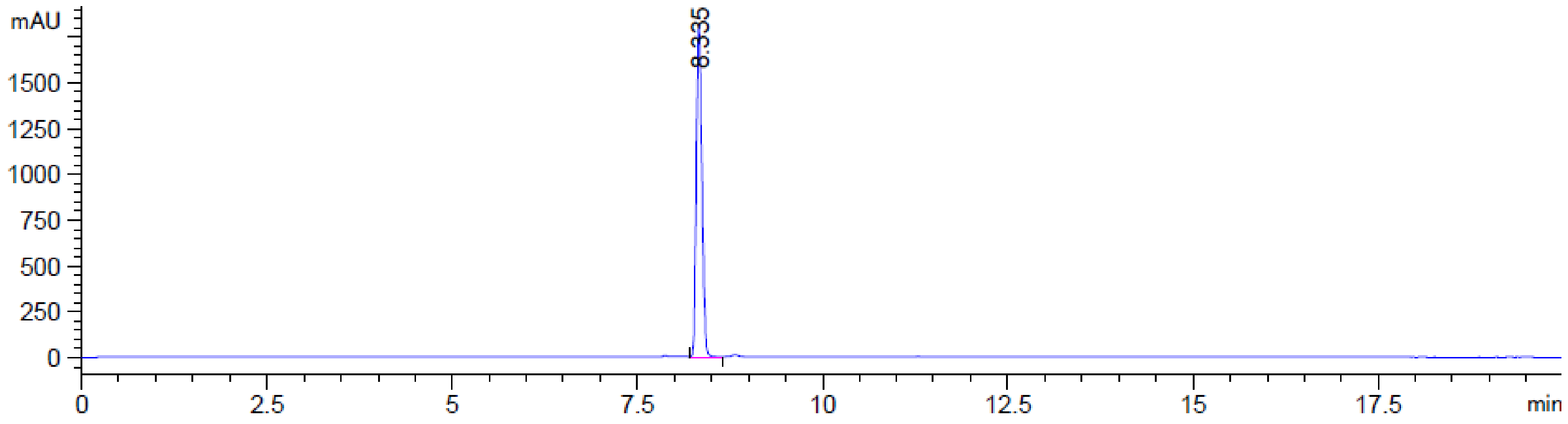
</td></tr>
</table>

**MS (-) ESI**

(Calc. [M-H]$^-$ $C_{39}H_{48}N_{15}O_{24}P_4$: 1234.19526)

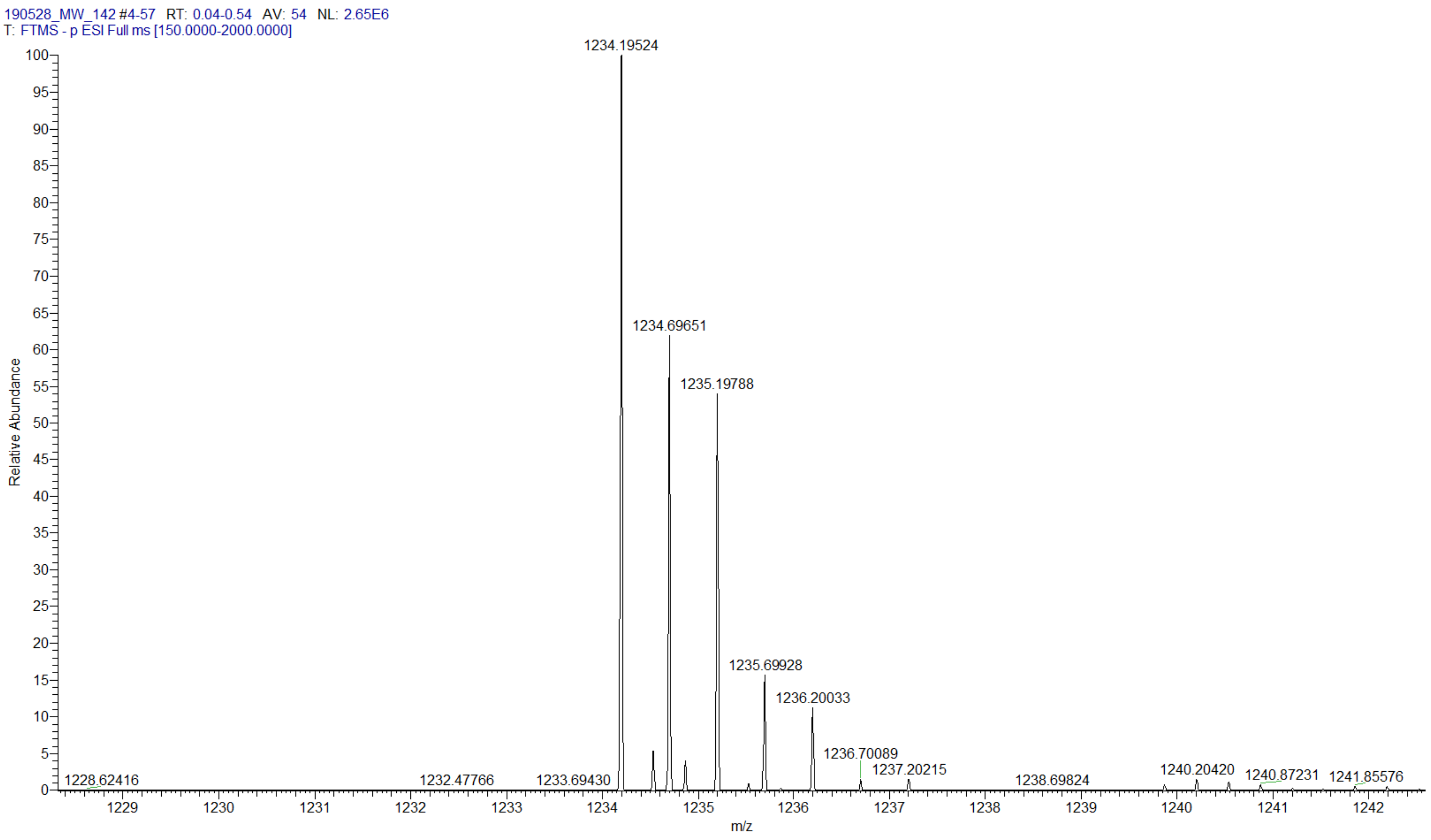

$^{1}H$ NMR (500 MHz, $D_2O$, 25°C)

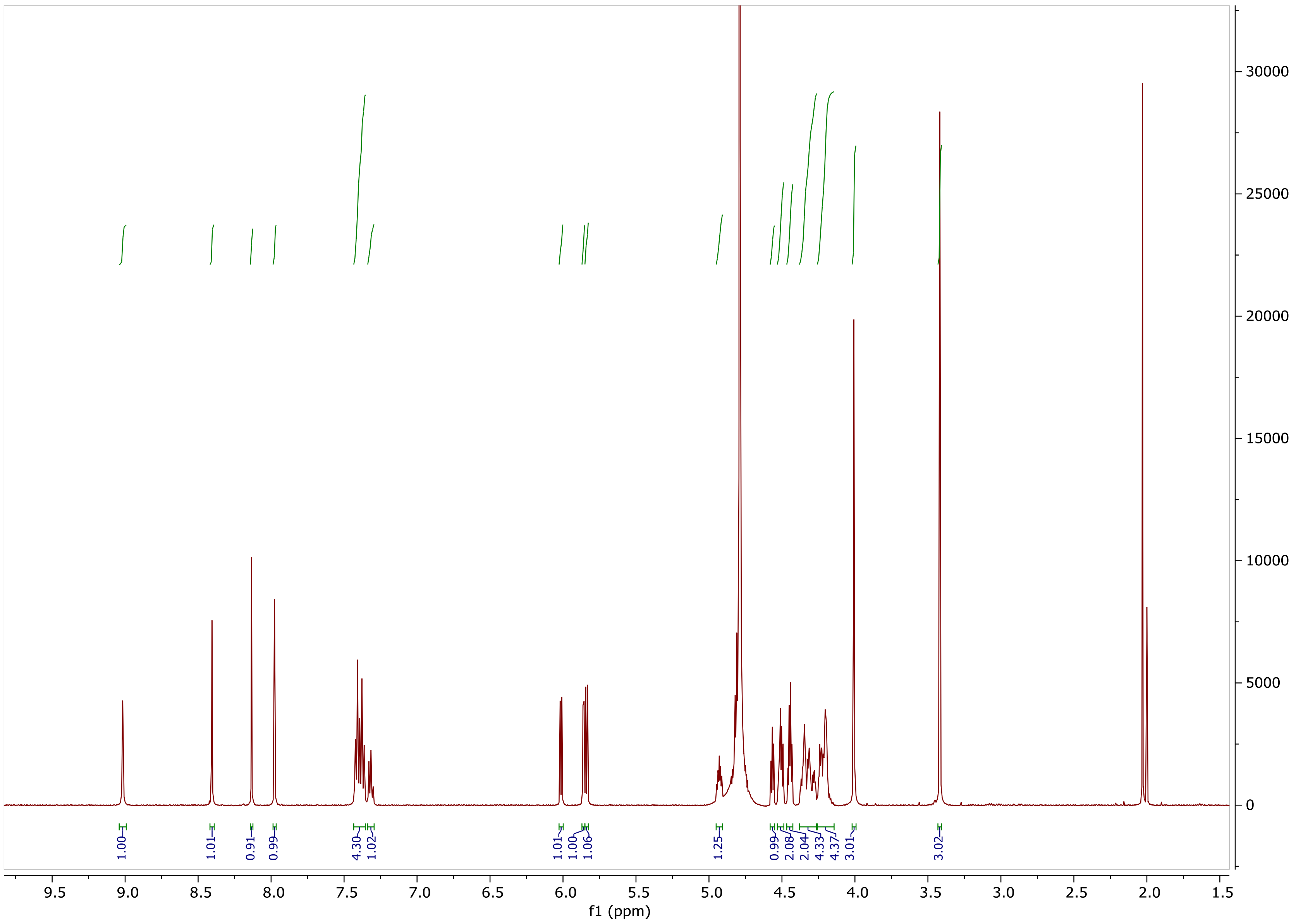

COSY NMR ($D_2O$, 25°)

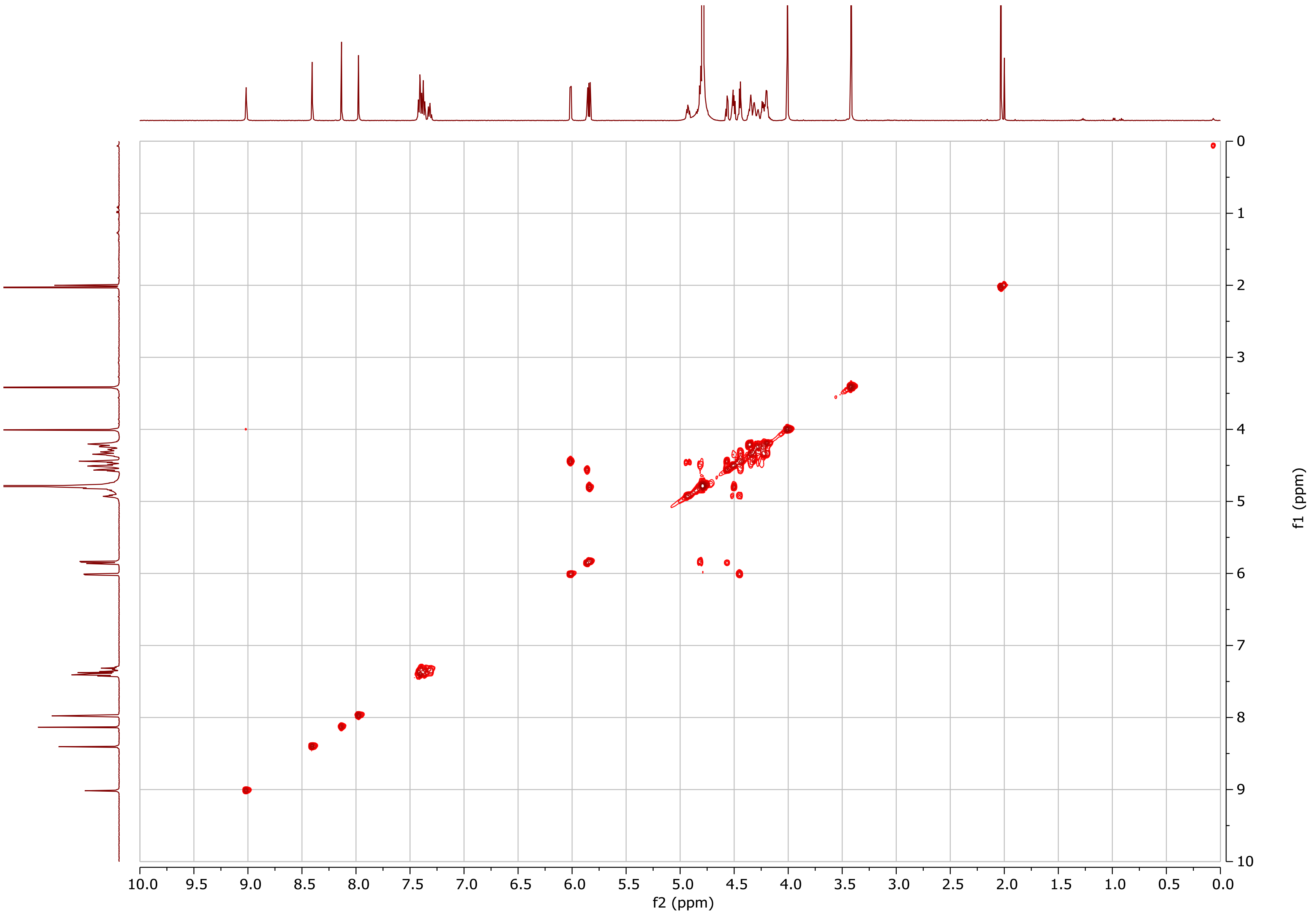

$^{31}P$ NMR (202.5 MHz, $D_2O$, 25°C)

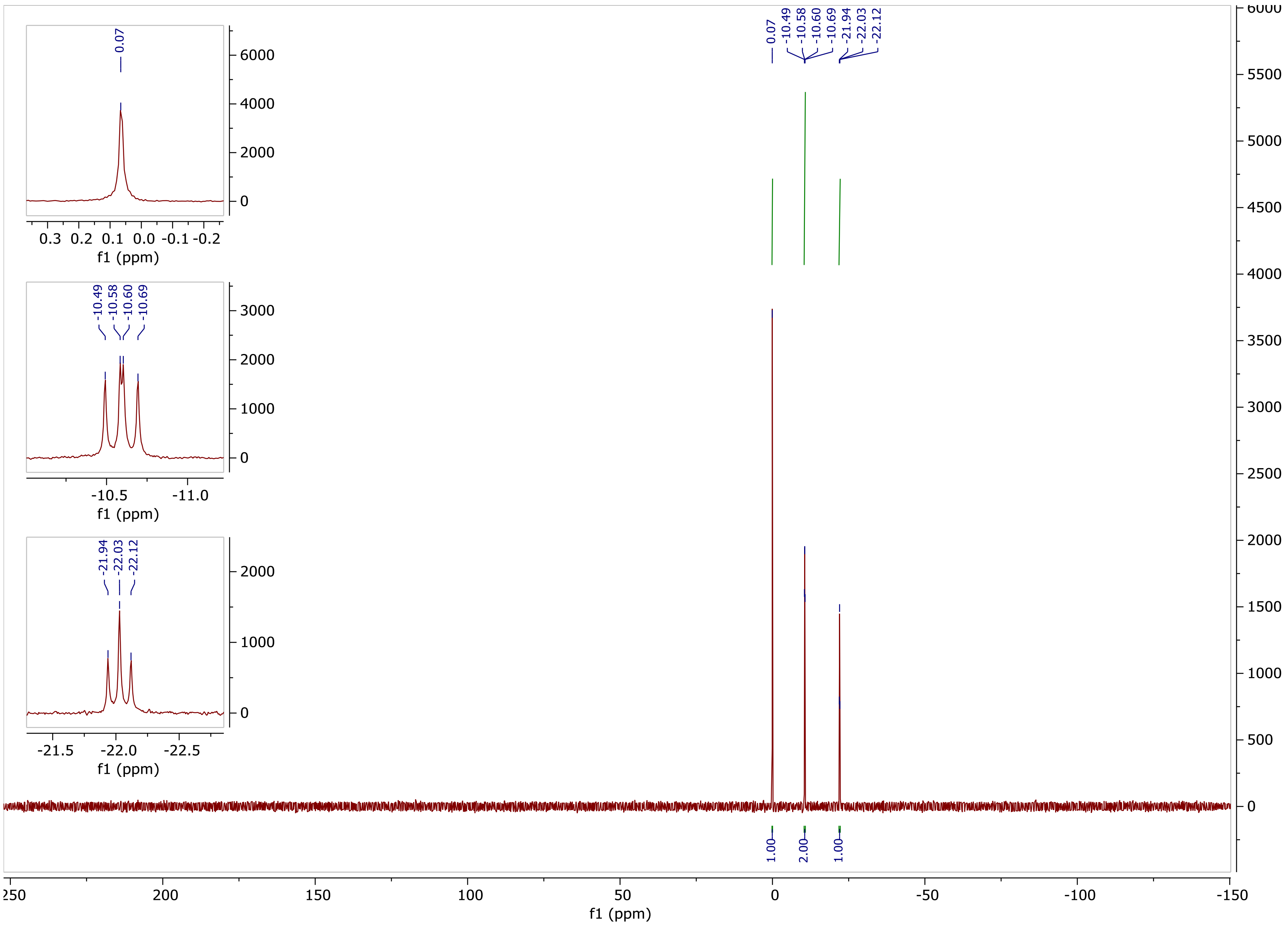

$^{1}H$-$^{13}C$ HSQC ($D_2O$, 25°C)

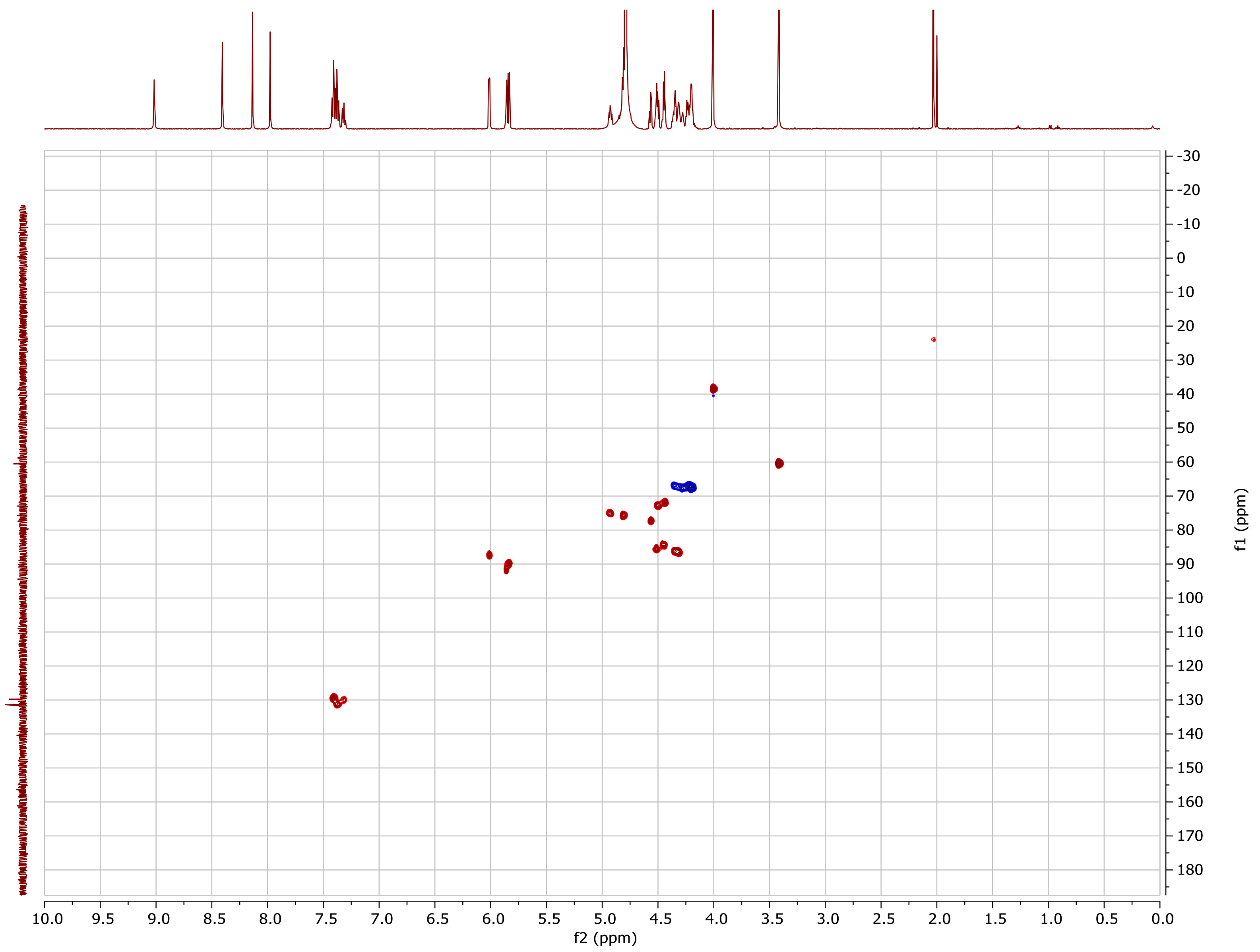

| (2a) m$^7$GpppA$_m$pG-L13$_N$ | |
|---|---|
| **Chemical structure** | 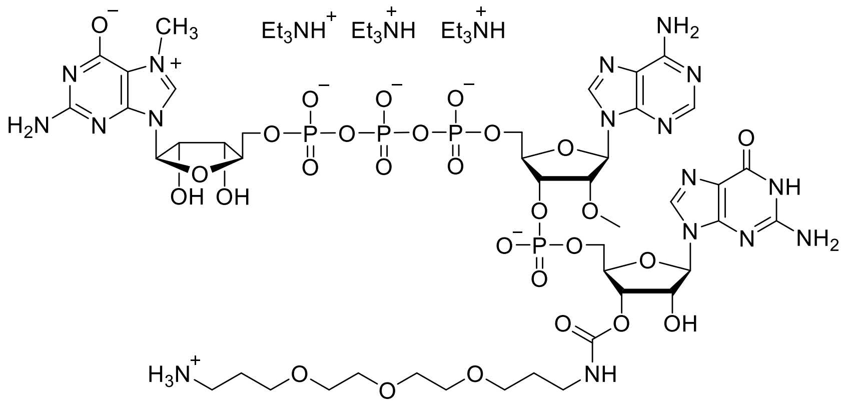 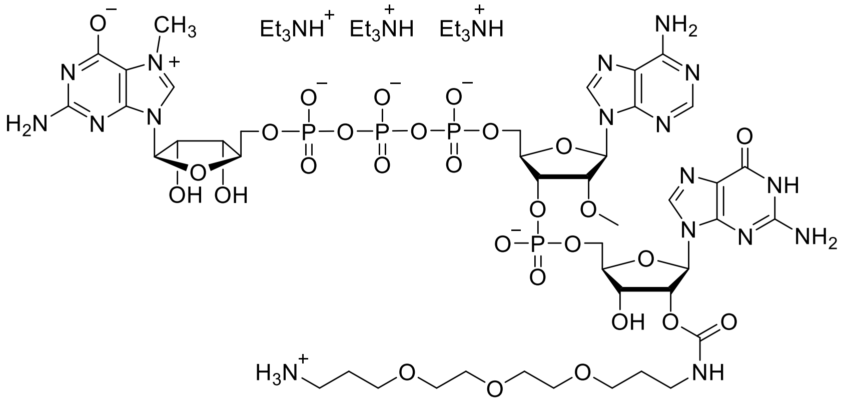 |
| **RP HPLC** Abs. @ 254 nm | 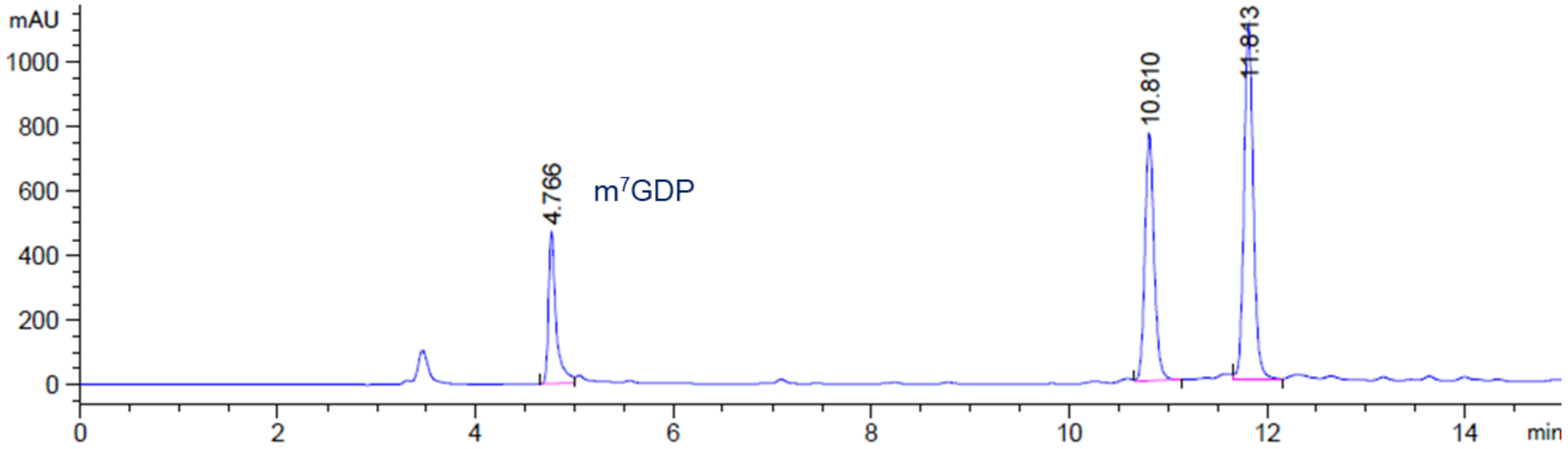 |

**MS (-) ESI**
(Calc. $[M-H]^- C_{43}H_{64}N_{17}O_{28}P_4$: 1390.30626)

*Isomer 1*

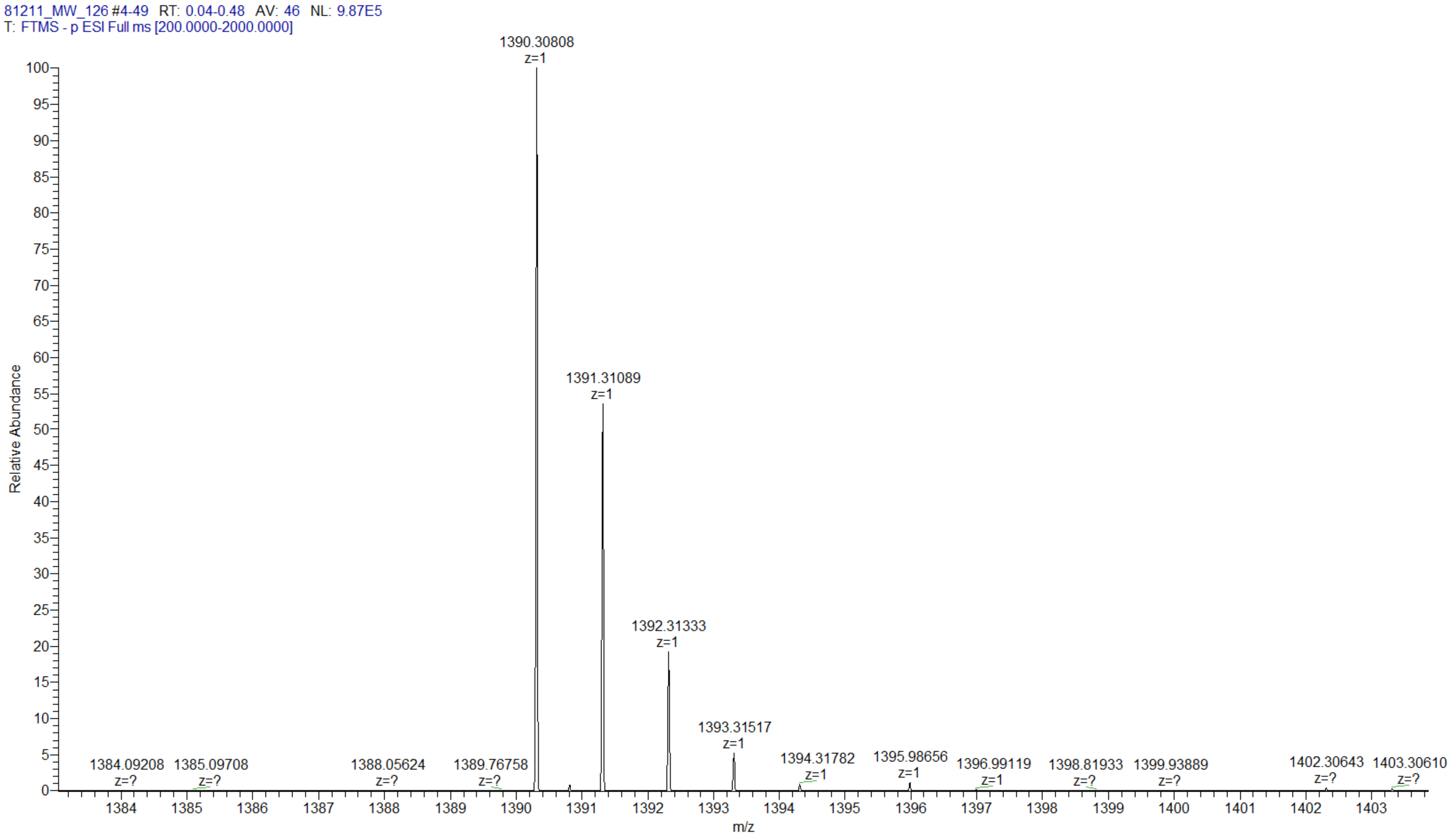

**MS (-) ESI**
(Calc. $[M-H]^-$ $C_{43}H_{64}N_{17}O_{28}P_4$: 1390.30626)

*Isomer 2*

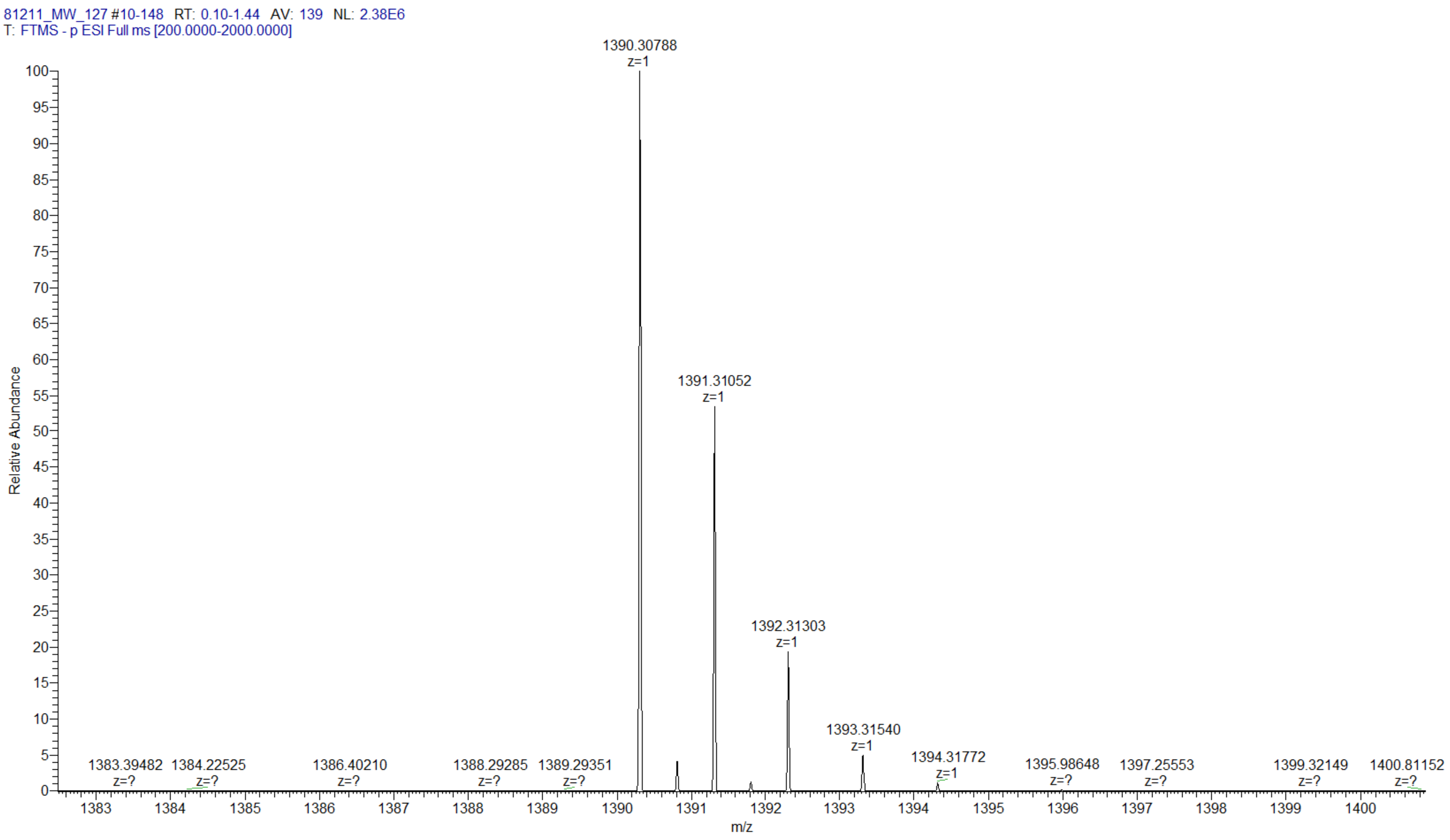

#### (2b) m$^7$Gppp$^{m6}$A$_m$pG-L13$_N$

**Chemical structure**

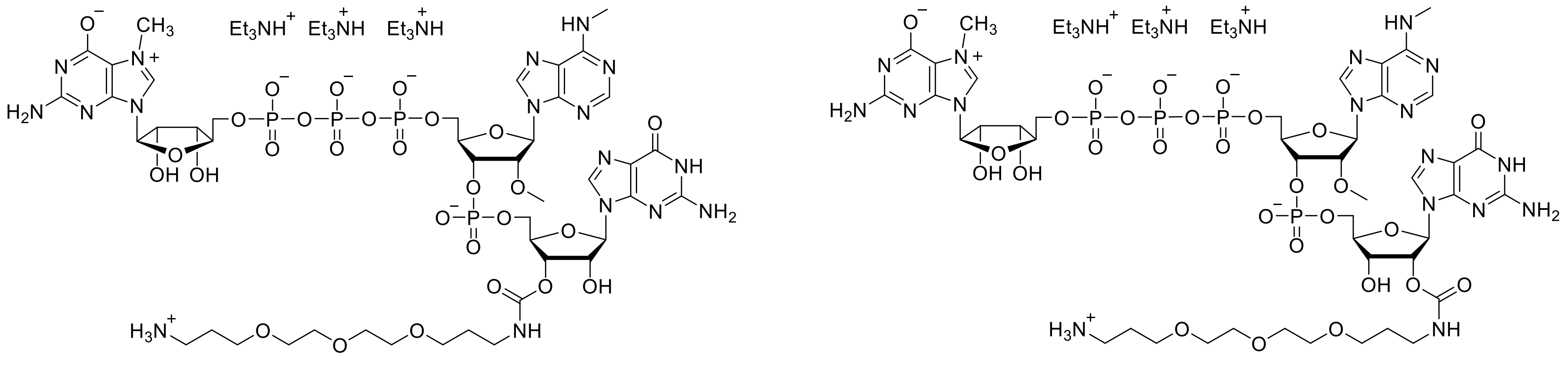

**RP HPLC**
Abs. @ 254 nm

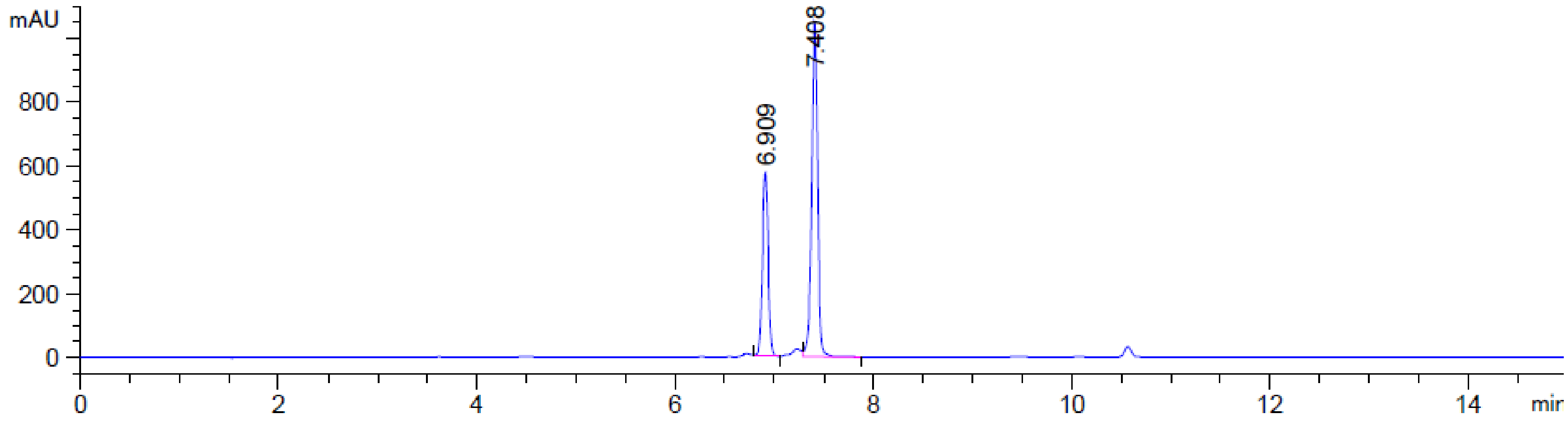

**MS (-) ESI**
(Calc. $[M\text{-}H]^-$ $C_{44}H_{66}N_{17}O_{28}P_4^-$ 1404.32191)

*Isomer 1*

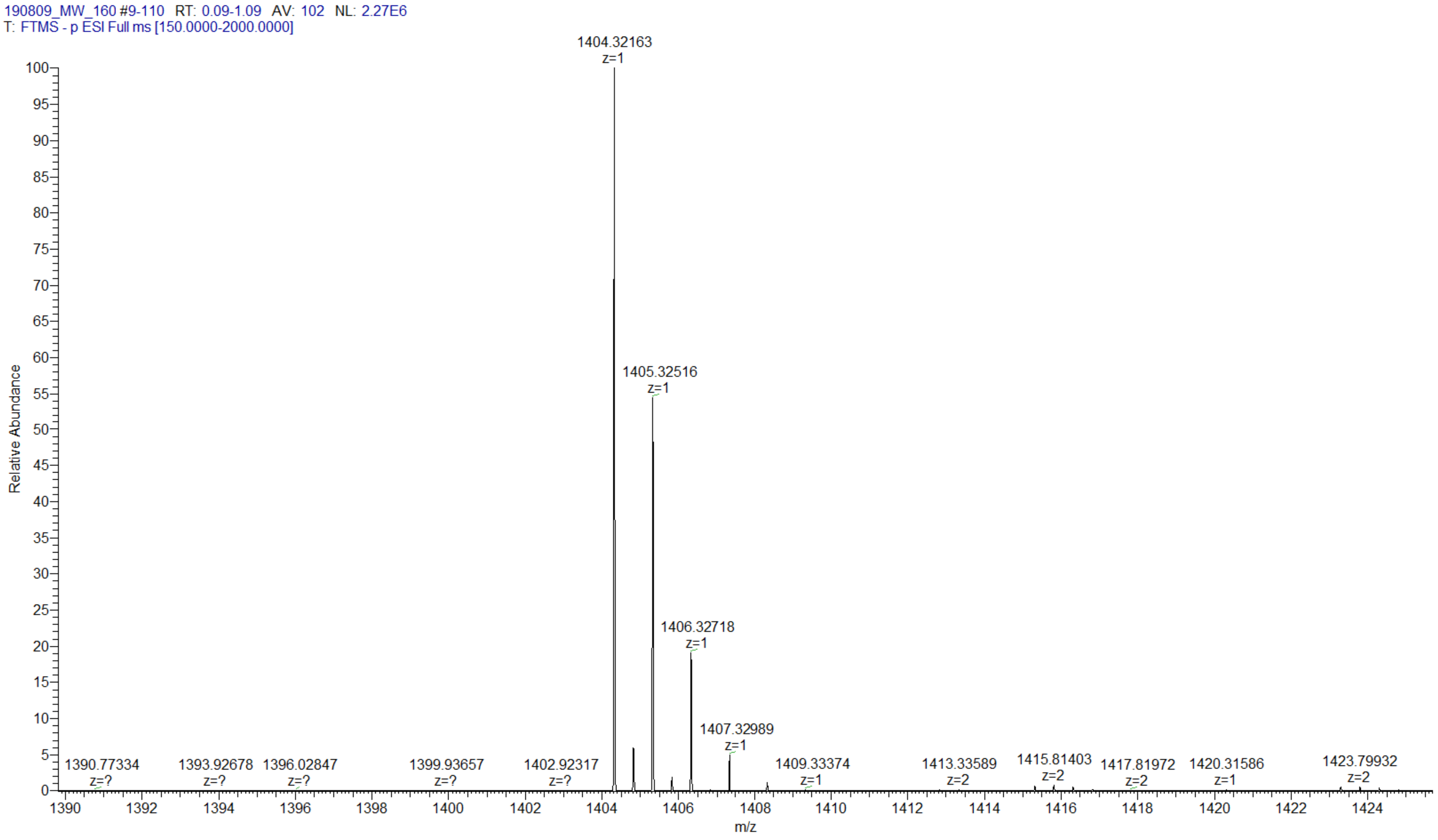

**MS (-) ESI**

(Calc. $[M-H]^-$ $C_{44}H_{66}N_{17}O_{28}P_4^-$ 1404.32191)

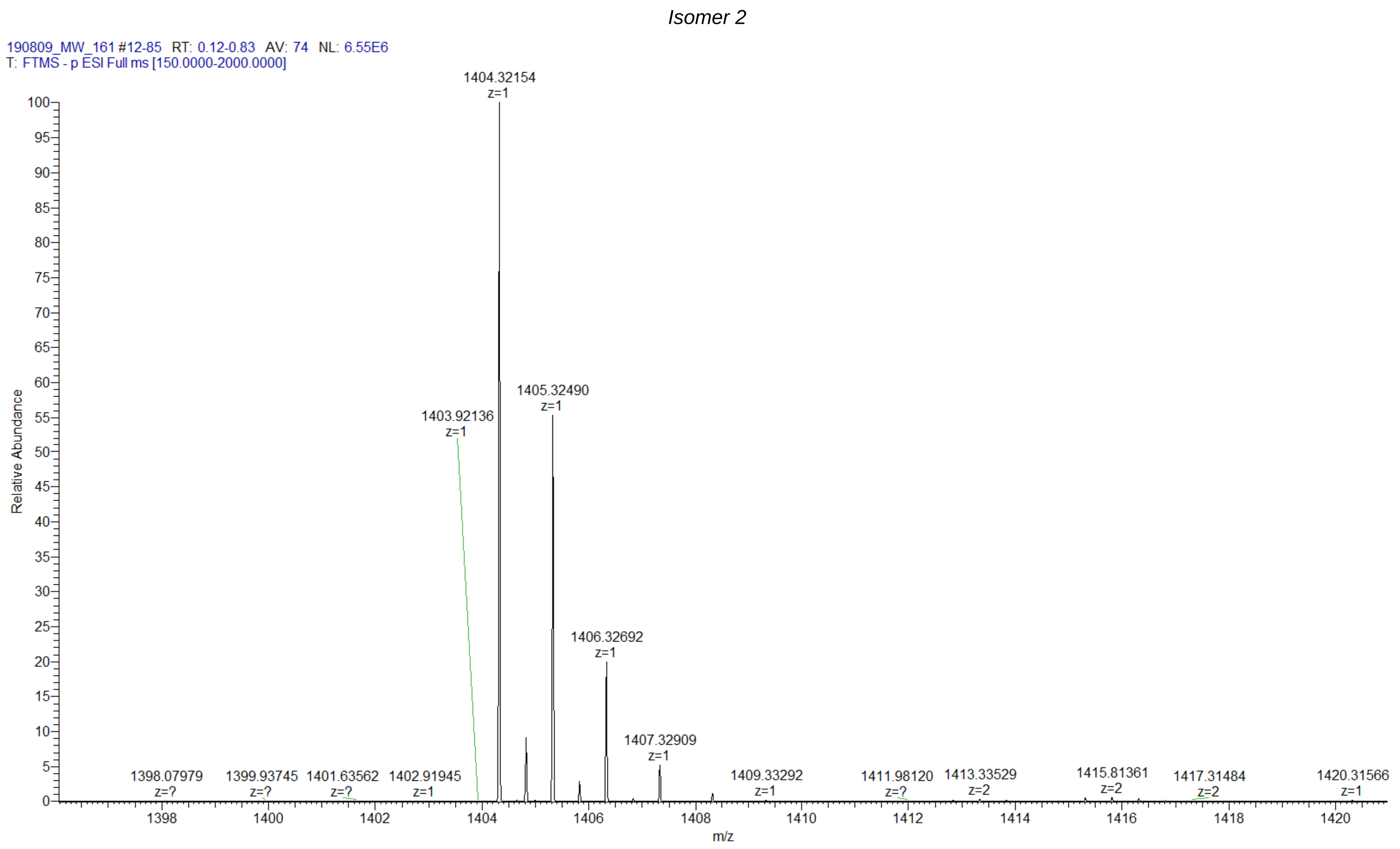

| (2c) m$^7$Gppp$^{Bn6}$A$_m$pG-L13$_N$ | |
|---|---|
| **Chemical structure** | 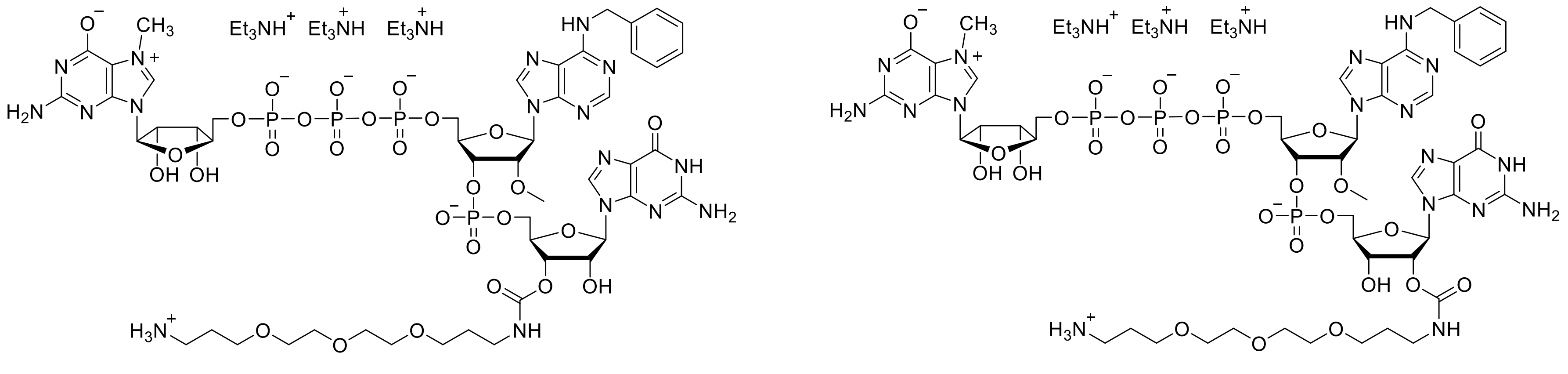 |
| **RP HPLC** Abs. @ 254 nm | 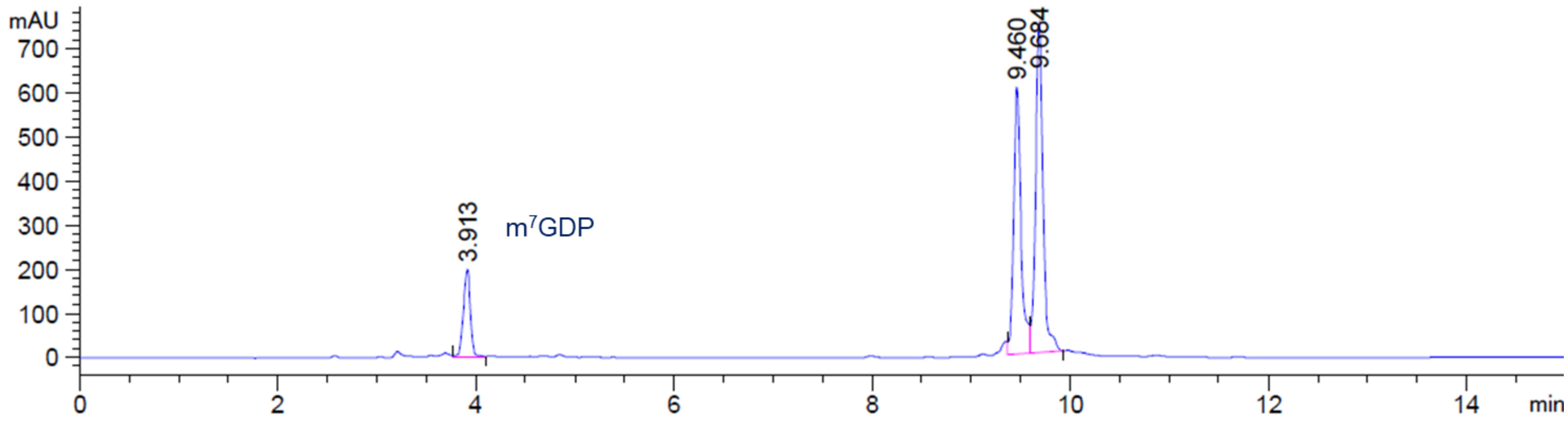 |

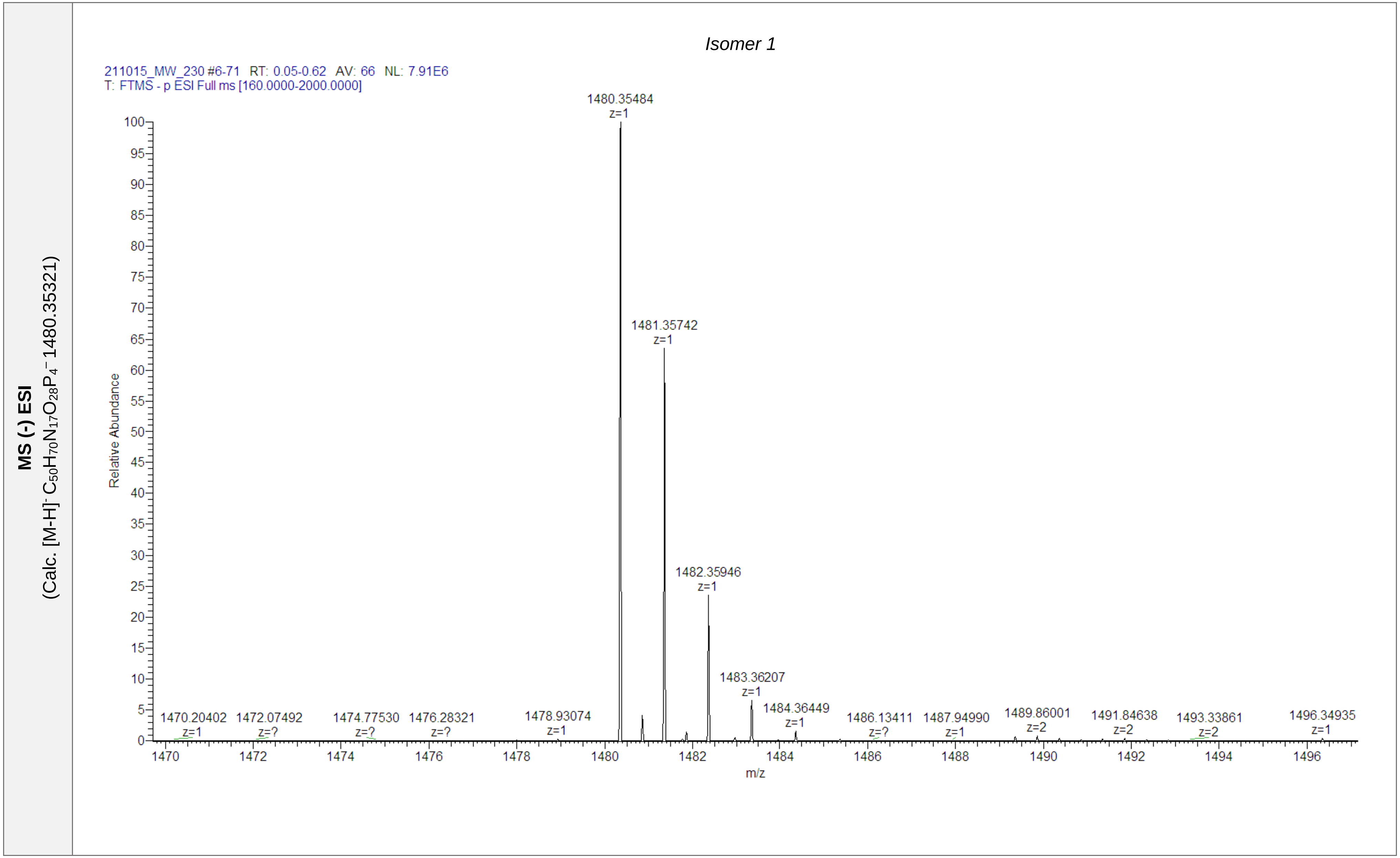
MS (-) ESI
(Calc. [M-H]- C50H70N17O28P4- 1480.35321)
Isomer 1
211015_MW_230 #6-71 RT: 0.05-0.62 AV: 66 NL: 7.91E6
T: FTMS - p ESI Full ms [160.0000-2000.0000]
1480.35484
z=1
1481.35742
z=1
1482.35946
z=1
1483.36207
z=1
1484.36449
z=1
1470.20402
z=1
1472.07492
z=?
1474.77530
z=?
1476.28321
z=?
1478.93074
z=1
1486.13411
z=?
1487.94990
z=1
1489.86001
z=2
1491.84638
z=2
1493.33861
z=2
1496.34935
z=1
Relative Abundance
m/z

**MS (-) ESI**
(Calc. [M-H]⁻ $C_{50}H_{69}N_{17}O_{28}P_4^{2-}$ 739.67297)

*Isomer 2*

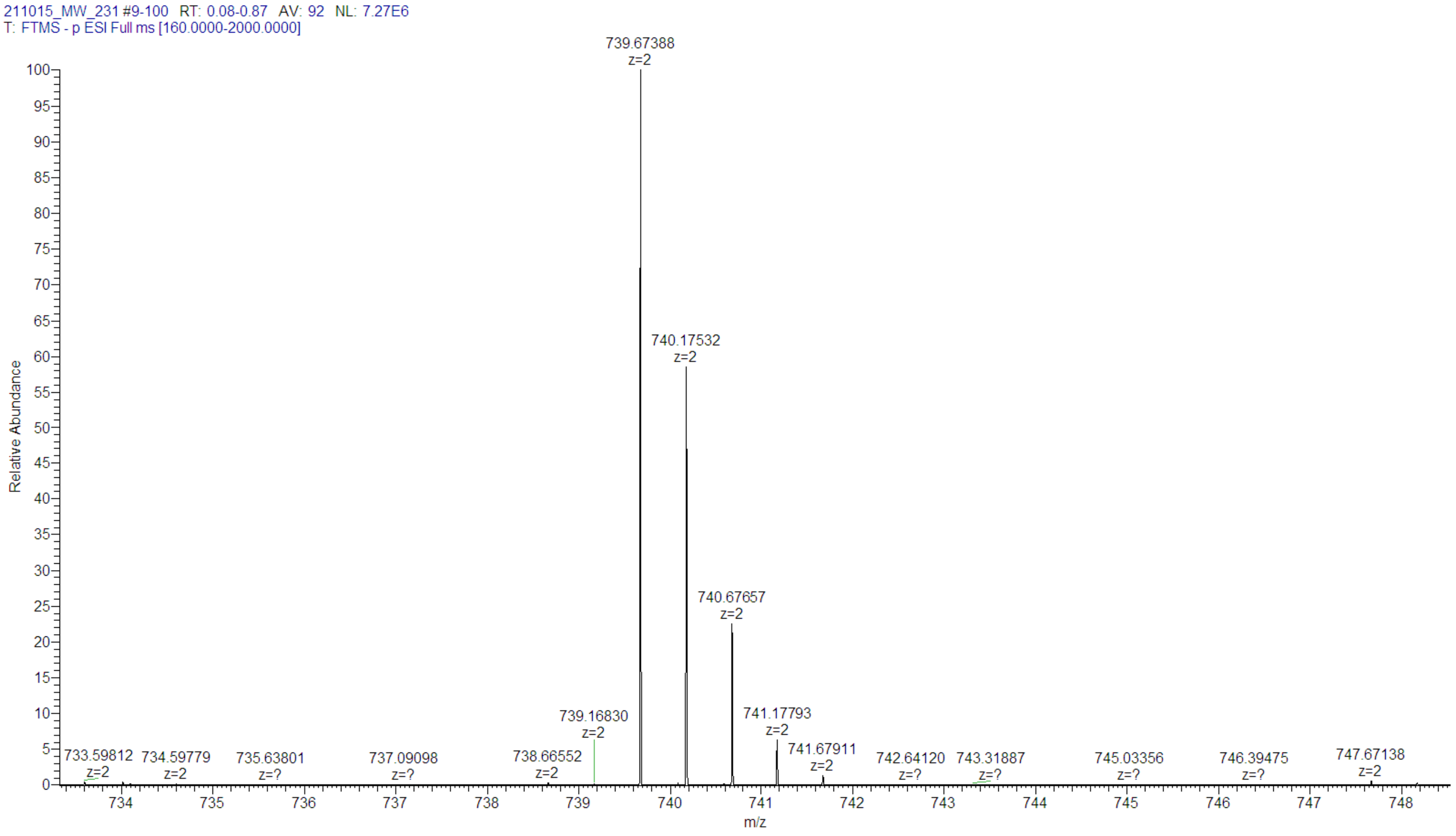

#### 10. Rerefences

Bednarczyk, M., Peters, J. K., Kasprzyk, R., Starek, J., Warminski, M., Spiewla, T., et al. (2022). Fluorescence-Based Activity Screening Assay Reveals Small Molecule Inhibitors of Vaccinia Virus mRNA Decapping Enzyme D9. *ACS Chemical Biology, 17*(6), 1460-1471.

Cox, J., & Mann, M. (2008). MaxQuant enables high peptide identification rates, individualized p.p.b.-range mass accuracies and proteome-wide protein quantification. *Nat. Biotechnol., 26*(12), 1367-1372.

Han, Z., Niu, T., Chang, J., Lei, X., Zhao, M., Wang, Q., et al. (2010). Crystal structure of the FTO protein reveals basis for its substrate specificity. *Nature, 464*(7292), 1205-1209.

Mukherjee, D., Fritz, D. T., Kilpatrick, W. J., Gao, M., & Wilusz, J. (2004). Analysis of RNA exonucleolytic activities in cellular extracts. *Methods Mol. Biol., 257*, 193-212.

Myers, S. A., Rhoads, A., Cocco, A. R., Peckner, R., Haber, A. L., Schweitzer, L. D., et al. (2019). Streamlined Protocol for Deep Proteomic Profiling of FAC-sorted Cells and Its Application to Freshly Isolated Murine Immune Cells. *Mol. Cell Proteomics, 18*(5), 995-1009.

Niedzwiecka, A., Marcotrigiano, J., Stepinski, J., Jankowska-Anyszka, M., Wyslouch-Cieszynska, A., Dadlez, M., et al. (2002). Biophysical Studies of eIF4E Cap-binding Protein: Recognition of mRNA 5′ Cap Structure and Synthetic Fragments of eIF4G and 4E-BP1 Proteins. *Journal of Molecular Biology, 319*(3), 615-635.

Sikorski, P. J., Warminski, M., Kubacka, D., Ratajczak, T., Nowis, D., Kowalska, J., et al. (2020). The identity and methylation status of the first transcribed nucleotide in eukaryotic mRNA 5′ cap modulates protein expression in living cells. *Nucleic Acids Research, 48*(4), 1607-1626.

Szczepaniak, S. A., Zuberek, J., Darzynkiewicz, E., Kufel, J., & Jemielity, J. (2012). Affinity resins containing enzymatically resistant mRNA cap analogs--a new tool for the analysis of cap-binding proteins. *RNA, 18*(7), 1421-1432.

Tyanova, S., Temu, T., Sinitcyn, P., Carlson, A., Hein, M. Y., Geiger, T., et al. (2016). The Perseus computational platform for comprehensive analysis of (prote)omics data. *Nat. Methods, 13*(9), 731-740.

Ziemkiewicz, K., Warminski, M., Wojcik, R., Kowalska, J., & Jemielity, J. (2022). Quick Access to Nucleobase-Modified Phosphoramidites for the Synthesis of Oligoribonucleotides Containing Post-Transcriptional Modifications and Epitranscriptomic Marks. *The Journal of Organic Chemistry, 87*(15), 10333-10348.

Zuberek, J., Kubacka, D., Jablonowska, A., Jemielity, J., Stepinski, J., Sonenberg, N., et al. (2007). Weak binding affinity of human 4EHP for mRNA cap analogs. *RNA, 13*(5), 691-697.

Zuberek, J., & Stelmachowska, A. (2017). Tryptophan Residues from Cap Binding Slot in eIF4E Family Members: Their Contributions to Near-UV Circular Dichroism Spectra. *Journal of Physical Chemistry & Biophysics, 7*(2).
