## Supplementary material for "Trinucleotide mRNA cap analog N6-benzylated at the site of posttranscriptional ^m6^Am mark facilitates mRNA purification and confers superior translational properties in vitro and in vivo": S2-Figures and Tables

[#]*Division of Biophysics, Institute of Experimental Physics, Faculty of Physics, University of Warsaw, 02-089 Warsaw, Poland;* [§]*Centre of New Technologies, University of Warsaw, 02-089 Warsaw, Poland;* [$]*Explorna Therapeutics sp. z o.o. Zwirki i Wigury 93, 02-089 Warsaw, Poland;* [«]*Laboratory of Epitranscriptomics, Department of Environmental Microbiology and Biotechnology, Institute of Microbiology, Faculty of Biology, Biological and Chemical Research Centre, University of Warsaw, 02-089 Warsaw, Poland;* [&]*Department of Chemistry & Biochemistry, University of Delaware, Newark DE 19716;* [♠]*Laboratory of Experimental Medicine, Faculty of Medicine, Medical University of Warsaw, 02-097 Warsaw, Poland;* [♣]*Proteomics Core Facility, IMol Polish Academy of Sciences, 02-247 Warsaw, Poland;* *correspondence to:

**Table S1: DNA sequences following the T7 promoter φ6.5 corresponding to in vitro transcribed mRNAs encoding firefly luciferase (Fluc), human erythropoietin (hEPO), Gaussia luciferase (Gluc), α1-antitrypsin (A1AT), SIINFEKL, ovalbumin (OVA), and NY-ESO1.**

| Name | DNA sequence |
|---|---|
| Firefly luciferase (Fluc) | GGGATAATCTAGACATTTGCTTCTGACACAACTGTGTTCACTAGCAACCTCAAACAGACACCATGGAAGACGCCAAAAACATAAAGAAAGGCCCGGCGCCATTCTATCC<br>TCTAGAGGATGGAACCGCTGGAGAGCAACTGCATAAGGCTATGAAGAGATACGCCCTGGTTCCTGGAACAATTGCTTTTACAGATGCACATATCGAGGTGAACATCACG<br>TACGCGGAATACTTCGAAATGTCCGTTCGGTTGGCAGAAGCTATGAAACGATATGGGCTGAATACAAATCACAGAATCGTCGTATGCAGTGAAAACTCTCTTCAATTCT<br>TTATGCCGGTGTTGGGCGCGTTATTTATCGGAGTTGCAGTTGCGCCCGCGAACGACATTTATAATGAACGTGAATTGCTCAACAGTATGAACATTTCGCAGCCTACCGT<br>AGTGTTTGTTTCCAAAAAGGGGTTGCAAAAAATTTTGAACGTGCAAAAAAAATTACCAATAATCCAGAAAATTATTATCATGGATTCTAAAACGGATTACCAGGGATTT<br>CAGTCGATGTACACGTTCGTCACATCTCATCTACCTCCCGGTTTTAATGAATACGATTTTGTACCAGAGTCCTTTGATCGTGACAAAACAATTGCACTGATAATGAATT<br>CCTCTGGATCTACTGGGTTACCTAAGGGTGTGGCCCTTCCGCATAGAACTGCCTGCGTCAGATTCTCGCATGCCAGAGATCCTATTTTTGGCAATCAAATCATTCCGGA<br>TACTGCGATTTTAAGTGTTGTTCCATTCCATCACGGTTTTGGAATGTTTACTACACTCGGATATTTGATATGTGGATTTCGAGTCGTCTTAATGTATAGATTTGAAGAA<br>GAGCTGTTTTTACGATCCCTTCAGGATTACAAAATTCAAAGTGCGTTGCTAGTACCAACCCTATTTTCATTCTTCGCCAAAAGCACTCTGATTGACAAATACGATTTAT<br>CTAATTTACACGAAATTGCTTCTGGGGGCGCACCTCTTTCGAAAGAAGTCGGGGAAGCGGTTGCAAAACGCTTCCATCTTCCAGGGATACGACAAGGATATGGGCTCAC<br>TGAGACTACATCAGCTATTCTGATTACACCCGAGGGGGATGATAAACCGGGCGCGGTCGGTAAAGTTGTTCCATTTTTTGAAGCGAAGGTTGTGGATCTGGATACCGGG<br>AAAACGCTGGGCGTTAATCAGAGAGGCGAATTATGTGTCAGAGGACCTATGATTATGTCCGGTTATGTAAACAATCCGGAAGCGACCAACGCCTTGATTGACAAGGATG<br>GATGGCTACATTCTGGAGACATAGCTTACTGGGACGAAGACGAACACTTCTTCATAGTTGACCGCTTGAAGTCTTTAATTAAATACAAAGGATATCAGGTGGCCCCCGC<br>TGAATTGGAATCGATATTGTTACAACACCCCAACATCTTCGACGCGGGCGTGGCAGGTCTTCCCGACGATGACGCCGGTGAACTTCCCGCCGCCGTTGTTGTTTTGGAG<br>CACGGAAAGACGATGACGGAAAAAGAGATCGTGGATTACGTCGCCAGTCAAGTAACAACCGCGAAAAAGTTGCGCGGAGGAGTTGTGTTTGTGGACGAAGTACCGAAAG<br>GTCTTACCGGAAAACTCGACGCAAGAAAAATCAGAGAGATCCTCATAAAGGCCAAGAAGGGCGGAAAGTCCAAATTGTAAGGATCCTAGGGCCGCAGCTCGCTTTCTTG<br>CTGTCCAATTTCTATTAAAGGTTCCTTTGTTCCCTAAGTCCAACTACTAAACTGGGGGATATTATGAAGGGCCTTGAGCATTTGGATTCTGCCTAATAAAAAACATTTA<br>TTTTCATTGCGTTTAGCTCGCTTTCTTGCTGTCCAATTTCTATTAAAGGTTCCTTTGTTCCCTAAGTCCAACTACTAAACTGGGGGATATTATGAAGGGCCTTGAGCAT<br>TTGGATTCTGCCTAATAAAAAACATTTATTTTCATTGCATTTAAATGTTTAAACAAAAAAAAAAAAAAAAAAAAAAAAAAAAAAAAAAAAAAAAAAAAAAAAAAA<br>AAAAAAAAAAAAAAAAAAAAAAAAAAAAAAAAAAA |
| Human erythropoietin (hEPO) | GGGATAATCTAGACATTTGCTTCTGACACAACTGTGTTCACTAGCAACCTCAAACAGACACCATGGGCGTGCACGAGTGCCCCGCCTGGCTGTGGCTGCTGCTGAGCCT<br>GCTGAGCCTGCCCCTGGGCCTGCCCGTGCTGGGCGCCCCCCCCCGGCTGATCTGCGACAGCCGGGTGCTGGAGCGGTACCTGCTGGAGGCCAAGGAGGCCGAGAACATC<br>ACCACCGGCTGCGCCGAGCACTGCAGCCTGAACGAGAACATCACCGTGCCCGACACCAAGGTGAACTTCTACGCCTGGAAGCGGATGGAGGTGGGCCAGCAGGCCGTGG<br>AGGTGTGGCAGGGCCTGGCCCTGCTGAGCGAGGCCGTGCTGCGGGGCCAGGCCCTGCTGGTGAACAGCAGCCAGCCCTGGGAGCCCCTGCAGCTGCACGTGGACAAGGC<br>CGTGAGCGGCCTGCGGAGCCTGACCACCCTGCTGCGGGCCCTGGGCGCCCAGAAGGAGGCCATCAGCCCCCCCGACGCCGCCAGCGCCGCCCCCCTGCGGACCATCACC<br>GCCGACACCTTCCGGAAGCTGTTCCGGGTGTACAGCAACTTCCTGCGGGGCAAGCTGAAGCTGTACACCGGCGAGGCCTGCCGGACCGGCGACCGGTGAGGATCCTAGG<br>GCCGCAGCTCGCTTTCTTGCTGTCCAATTTCTATTAAAGGTTCCTTTGTTCCCTAAGTCCAACTACTAAACTGGGGGATATTATGAAGGGCCTTGAGCATTTGGATTCT<br>GCCTAATAAAAAACATTTATTTTCATTGCGTTTAGCTCGCTTTCTTGCTGTCCAATTTCTATTAAAGGTTCCTTTGTTCCCTAAGTCCAACTACTAAACTGGGGGATAT<br>TATGAAGGGCCTTGAGCATTTGGATTCTGCCTAATAAAAAACATTTATTTTCATTGCATTTAAATGTTTAAACAAAAAAAAAAAAAAAAAAAAAAAAAAAAAAAAAAAA<br>AAAAAAAAAAAAAAAAAAAAAAAAAAAAAAAAAAAAAAAAAAAAAAAAAAAAAA |
| *Gaussia* luciferase (Gluc) | GGGATAATCTAGACATTTGCTTCTGACACAACTGTGTTCACTAGCAACCTCAAACAGACACCATGGGAGTCAAAGTTCTGTTTGCCCTGATCTGCATCGCTGTGGCCGA<br>GGCCAAGCCCACCGAGAACAACGAAGACTTCAACATCGTGGCCGTGGCCAGCAACTTCGCGACCACGGATCTCGATGCTGACCGCGGGAAGTTGCCCGGCAAGAAGCTG<br>CCGCTGGAGGTGCTCAAAGAGTTGGAAGCCAATGCCCGGAAAGCTGGCTGCACCAGGGGCTGTCTGATCTGCCTGTCCCACATCAAGTGCACGCCCAAGATGAAGAAGT<br>TCATCCCAGGACGCTGCCACACCTACGAAGGCGACAAAGAGTCCGCACAGGGCGGCATAGGCGAGGCGATCGTCGACATTCCTGAGATTCCTGGGTTCAAGGACTTGGA<br>GCCCTTGGAGCAGTTCATCGCACAGGTCGATCTGTGTGTGGACTGCACAACTGGCTGCCTCAAAGGGCTTGCCAACGTGCAGTGTTCTGACCTGCTCAAGAAGTGGCTG<br>CCGCAACGCTGTGCGACCTTTGCCAGCAAGATCCAGGGCCAGGTGGACAAGATCAAGGGGGCCGGTGGTGACTAAGGATCCTAGGGCCGCAGCTCGCTTTCTTGCTGTC<br>CAATTTCTATTAAAGGTTCCTTTGTTCCCTAAGTCCAACTACTAAACTGGGGGATATTATGAAGGGCCTTGAGCATTTGGATTCTGCCTAATAAAAAACATTTATTTTC<br>ATTGCGTTTAGCTCGCTTTCTTGCTGTCCAATTTCTATTAAAGGTTCCTTTGTTCCCTAAGTCCAACTACTAAACTGGGGGATATTATGAAGGGCCTTGAGCATTTGGA<br>TTCTGCCTAATAAAAAACATTTATTTTCATTGCATTTAAATGTTTAAACAAAAAAAAAAAAAAAAAAAAAAAAAAAAAAAAAAA |

| | |
|---|---|
| α1-antitrypsin (hA1AT) | GGGATAATACATTTGCTTCTGACACAACTGTGTTCACTAGCAACCTCAAACAGACACCGCCACCATGCCTAGCAGCGTGTCATGGGGTATTCTGCTGCTGGCCGGACTG<br>TGTTGTCTGGTGCCTGTGTCTCTGGCTGAAGATCCTCAAGGCGACGCCGCTCAGAAAACCGATACAAGCCACCACGACCAGGATCACCCCACCTTCAACAAGATCACCC<br>CTAACCTGGCCGAGTTCGCCTTCAGCCTGTATAGACAGCTGGCCCACCAGAGCAACAGCACCAACATCTTTTTCAGCCCCGTGTCTATCGCCACCGCCTTTGCTATGCT<br>GAGCCTGGGCACAAAGGCCGACACACACGATGAGATCCTGGAAGGCCTGAACTTCAACCTGACAGAGATCCCCGAGGCTCAGATCCACGAGGGCTTTCAAGAGCTGCTG<br>AGAACCCTGAACCAGCCTGACTCTCAGCTCCAGCTGACAACCGGCAATGGCCTGTTTCTGTCTGAGGGCCTGAAGCTGGTGGACAAGTTCCTGGAAGATGTGAAGAAGC<br>TGTACCACAGCGAGGCCTTCACCGTGAACTTCGGCGATACCGAGGAAGCCAAGAAGCAGATCAACGACTACGTGGAAAAGGGCACCCAGGGCAAGATCGTGGACCTGGT<br>CAAAGAGCTGGACAGAGACACCGTGTTCGCCCTGGTCAACTACATCTTCTTCAAAGGCAAGTGGGAACGCCCCTTCGAAGTGAAGGACACAGAGGAAGAGGACTTCCAC<br>GTCGACCAAGTGACCACCGTGAAGGTGCCCATGATGAAGCGGCTGGGCATGTTCAACATCCAGCACTGCAAGAAACTGAGCAGCTGGGTGCTGCTGATGAAGTACCTGG<br>GCAACGCCACAGCCATATTCTTTCTGCCCGATGAGGGCAAGCTGCAGCACCTGGAAAATGAGCTGACCCACGACATCATCACCAAGTTCCTAGAGAACGAGGACAGAAG<br>AAGCGCCAGCCTGCATCTGCCTAAGCTGAGCATCACCGGCACCTACGATCTGAAGTCTGTGCTGGGACAGCTGGGCATCACAAAGGTGTTCAGCAATGGCGCCGATCTG<br>TCCGGCGTTACAGAAGAGGCTCCTCTGAAGCTGTCCAAGGCCGTGCACAAAGCCGTGCTGACAATCGATGAGAAGGGAACAGAGGCCGCTGGCGCCATGTTTCTGGAAG<br>CTATCCCTATGAGCATCCCGCCTGAAGTGAAGTTCAACAAGCCCTTCGTGTTCCTGATGATCGAACAGAATACCAAGTCTCCCCTGTTCATGGGCAAAGTGGTCAACCC<br>CACACAGAAATGAGGATCCAGCTCGCTTTCTTGCTGTCCAATTTCTATTAAAGGTTCCTTTGTTCCCTAAGTCCAACTACTAAACTGGGGGATATTATGAAGGGCCTTG<br>AGCATTTGGATTCTGCCTAATAAAAAACATTTATTTTCATTGCCTCGAGAGCTCGCTTTCTTGCTGTCCAATTTCTATTAAAGGTTCCTTTGTTCCCTAAGTCCAACTA<br>CTAAACTGGGGGATATTATGAAGGGCCTTGAGCATTTGGATTCTGCCTAATAAAAAACATTTATTTTCATTGCACTAGTAAAAAAAAAAAAAAAAAAAAAAAAAAAAAA<br>AAAAAAAAAAAAAAAAAAAAAAAAAAAAAAAAAAAAAAAAAAAAAAAAAAAAAAAAAAAA |
| SIINFEKL | GGGATAATCTAGACATTTGCTTCTGACACAACTGTGTTCACTAGCAACCTCAAACAGACACCATGAGAGTGACAGCCCCTCGGACACTGATTCTGCTGCTTTCTGGTGC<br>CCTGGCTCTGACAGAAACATGGGCCGGATCTCTGGAACAGCTGGAATCCATCATCAACTTCGAGAAGCTGACCGAGTGGACCAGCTGAGGATCCTAGGGCCGCAGCTCG<br>CTTTCTTGCTGTCCAATTTCTATTAAAGGTTCCTTTGTTCCCTAAGTCCAACTACTAAACTGGGGGATATTATGAAGGGCCTTGAGCATTTGGATTCTGCCTAATAAAA<br>AACATTTATTTTCATTGCGTTTAGCTCGCTTTCTTGCTGTCCAATTTCTATTAAAGGTTCCTTTGTTCCCTAAGTCCAACTACTAAACTGGGGGATATTATGAAGGGCC<br>TTGAGCATTTGGATTCTGCCTAATAAAAAACATTTATTTTCATTGCATTTAAATGTTTAAACAAAAAAAAAAAAAAAAAAAAAAAAAAAAAAAAAAAAAAAAAAAAAAA<br>AAAAAAAAAAAAAAAAAAAAAAAAAAAAAAAAAAAAAAAAAAA |
| Ovalbumin (OVA) | GGGATAATACATTTGCTTCTGACACAACTGTGTTCACTAGCAACCTCAAACAGACACCGCCACCATGGGCAGCATCGGCGCCGCCAGCATGGAGTTCTGCTTCGACGTG<br>TTCAAGGAGCTGAAGGTGCACCACGCCAACGAGAACATCTTCTACTGCCCCATCGCCATCATGAGCGCCCTGGCCATGGTGTACCTGGGCGCCAAGGACAGCACCCGGA<br>CCCAGATCAACAAGGTGGTGCGGTTCGACAAGCTGCCCGGCTTCGGCGACAGCATCGAGGCCCAGTGCGGCACCAGCGTGAACGTGCACAGCAGCCTGCGGGACATCCT<br>GAACCAGATCACCAAGCCCAACGACGTGTACAGCTTCAGCCTGGCCAGCCGGCTGTACGCCGAGGAGCGGTACCCCATCCTGCCCGAGTACCTGCAGTGCGTGAAGGAG<br>CTGTACCGGGGCGGCCTGGAGCCCATCAACTTCCAGACCGCCGCCGACCAGGCCCGGGAGCTGATCAACAGCTGGGTGGAGAGCCAGACCAACGGCATCATCCGGAACG<br>TGCTGCAGCCCAGCAGCGTGGACAGCCAGACCGCCATGGTGCTGGTGAACGCCATCGTGTTCAAGGGCCTGTGGGAGAAGACCTTCAAGGACGAGGACACCCAGGCCAT<br>GCCCTTCCGGGTGACCGAGCAGGAGAGCAAGCCCGTGCAGATGATGTACCAGATCGGCCTGTTCCGGGTGGCCAGCATGGCCAGCGAGAAGATGAAGATCCTGGAGCTG<br>CCCTTCGCCAGCGGCACCATGAGCATGCTGGTGCTGCTGCCCGACGAGGTGAGCGGCCTGGAGCAGCTGGAGAGCATCATCAACTTCGAGAAGCTGACCGAGTGGACCA<br>GCAGCAACGTGATGGAGGAGCGGAAGATCAAGGTGTACCTGCCCCGGATGAAGATGGAGGAGAAGTACAACCTGACCAGCGTGCTGATGGCCATGGGCATCACCGACGT<br>GTTCAGCAGCAGCGCCAACCTGAGCGGCATCAGCAGCGCCGAGAGCCTGAAGATCAGCCAGGCCGTGCACGCCGCCCACGCCGAGATCAACGAGGCCGGCCGGGAGGTG<br>GTGGGCAGCGCCGAGGCCGGCGTGGACGCCGCCAGCGTGAGCGAGGAGTTCCGGGCCGACCACCCCTTCCTGTTCTGCATCAAGCACATCGCCACCAACGCCGTGCTGT<br>TCTTCGGCCGGTGCGTGAGCCCCTGAGGATCCAGCTCGCTTTCTTGCTGTCCAATTTCTATTAAAGGTTCCTTTGTTCCCTAAGTCCAACTACTAAACTGGGGGATATT<br>ATGAAGGGCCTTGAGCATTTGGATTCTGCCTAATAAAAAACATTTATTTTCATTGCCTCGAGAGCTCGCTTTCTTGCTGTCCAATTTCTATTAAAGGTTCCTTTGTTCC<br>CTAAGTCCAACTACTAAACTGGGGGATATTATGAAGGGCCTTGAGCATTTGGATTCTGCCTAATAAAAAACATTTATTTTCATTGCACTAGTAAAAAAAAAAAAAAAAA<br>AAAAAAAAAAAAAAAAAAAAAAAAAAAAAAAAAAAAAAAAAAAAAAAAAAAAAAAAAAAAAAAAAAAAAAAAA |
| NY-ESO1 | GGGATAATACATTTGCTTCTGACACAACTGTGTTCACTAGCAACCTCAAACAGACACCGCCACCATGCAGGCCGAAGGCCGCGGAACCGGCGGCAGCACCGGCGACGCC<br>GACGGACCAGGCGGCCCAGGCATCCCGGACGGCCCAGGCGGGAACGCAGGCGGACCCGGCGAGGCAGGCGCCACAGGCGGCAGAGGCCCCAGAGGCGCAGGCGCCGCAC<br>GCGCAAGCGGCCCAGGCGGCGGAGCCCCACGGGGCCCACACGGCGGAGCGGCCAGCGGCCTGAACGGCTGCTGCCGCTGCGGCGCCAGAGGGCCCGAGAGCCGCCTGCT<br>CGAGTTCTACCTGGCCATGCCGTTCGCGACGCCCATGGAAGCCGAGCTGGCCAGGCGGAGCCTGGCGCAGGACGCCCCACCGCTGCCGGTGCCGGGCGTGCTGCTGAAA<br>GAGTTCACCGTCAGCGGCAACATCCTGACCATCCGGCTGACAGCCGCCGACCACCGCCAGCTCCAGCTGAGCATCAGCAGCTGCCTCCAGCAGCTGAGCCTGCTGATGT<br>GGATCACCCAATGCTTTCTGCCCGTGTTCCTGGCACAGCCGCCAAGCGGACAGCGGCGCTAGGGATCCCAAGCACGCAGCAATGCAGCTCAAAACGCTTAGCCTAGCCA<br>CACCCCCACGGGAAACAGCAGTGATTAACCTTTAGCAATAAACGAAAGTTTAACTAAGCTATACTAACCCCAGGGTTGGTCAATTTCGTGCCAGCCACACCCTGGTACT<br>GCATGCACGCAATGCTAGCTGCCCCTTTCCCGTCCTGGGTACCCCGAGTCTCCCCCGACCTCGGGTCCCAGGTATGCTCCCACCTCCtCCTGCCCCACTCACCACCTCT<br>GCTAGTTCCAGACACCTCCACTAGTAAAAAAAAAAAAAAAAAAAAAAAAAAAAAAAAAAAAAAAAAAAAAAAAAAAAAAAAAAAAAAAAAAAAAAAAAAAAAA<br>AAAAAA |

**Table S2. Quality parameters of mRNAs prepared for experiments.**

| No. | Transcript length | CDS | Cap | IVT efficiency [μg/μL] [a] | Capping before HPLC [b] | Capping after HPLC [c] | dsRNA content [d] | Figure [e] | Protocol for mRNA preparation [f] |
|---|---|---|---|---|---|---|---|---|---|
| 1 | 2089 nt | Fluc | $m^{7}GpppA_{m}pG$ | 1.2 | nd. | 95% | not detected [g] | 4A | C |
| 2 | 2089 nt | Fluc | $m^{7}Gppp^{m6}A_{m}pG$ | 2.9 | nd. | 84% | not detected [g] | 4A | C |
| 3 | 2089 nt | Fluc | $m^{7}Gppp^{Bn6}A_{m}pG$ | 2.8 | nd. | 91% | not detected [g] | 4A | C |
| 4 | 1040 nt | hEPO | $m^{7}GpppA_{m}pG$ | 3.71 | 88% | 90% | <0.125 ng [h]<br><0.05 ‰ [i] | 4B | C |
| 5 | 1040 nt | hEPO | $m^{7}Gppp^{Bn6}A_{m}pG$ | 3.85 | 77% | 93% | not detected [j] | 4B | C |
| 6 | 956 nt | Gluc | $m^{7}GpppA_{m}pG$ | 0.86 | nd. | nd. | not detected [k] | 4C, S5 | A |
| 7 | 956 nt | Gluc | $m^{7}Gppp^{Bn6}A_{m}pG$ | 0.72 | nd. | nd. | not detected [k] | 4C, S5 | A |
| 8 | 2089 nt | Fluc | $m^{7}GpppA_{m}pG$ | 4.14 | 94% | 95% | 0.078-0.156 ng [h]<br>0.03-0.06‰ [i] | 5A, 5B, S2, S3 | C |
| 9 | 2089 nt | Fluc | $m^{7}Gppp^{Bn6}A_{m}pG$ | 4.28 | 90% | 97% | not detected [l] | 5A, 5B, S2, S3 | C |
| 10 | 1040 nt | hEPO | $m^{7}GpppA_{m}pG$ | 3.5 | 92% | 93% | not detected [l] | 5C, S2, S4 | C |
| 11 | 1040 nt | hEPO | $m^{7}Gppp^{Bn6}A_{m}pG$ | 3.66 | 87% | 97% | not detected [l] | 5C, S2, S4 | C |
| 12 | 588 nt | SIINFEKL | $m^{7}GpppA_{m}pG$ | 3.47 | nd. | nd. | not detected [k] | 6A | B |
| 13 | 588 nt | SIINFEKL | $m^{7}Gppp^{Bn6}A_{m}pG$ | 2.49 | nd. | nd. | not detected [k] | 6A | B |
| 14 | 1599 nt | OVA | $m^{7}GpppA_{m}pG$ | 3.25 | nd. | nd. | not detected [m] | 6B | B |
| 15 | 1599 nt | OVA | $m^{7}Gppp^{Bn6}A_{m}pG$ | 3.22 | nd. | nd. | not detected [m] | 6B | B |
| 16 | 2089 nt | Fluc | $m^{7}Gppp^{Bn6}A_{m}pG$ | 3.79 | nd. | nd. | not detected [m] | 6B | B |
| 17 | 987 nt | NY-ESO1 | $m^{7}GpppA_{m}pG$ | 3.71 | 97% | 93% | not detected [n] | 6C | C |
| 18 | 987 nt | NY-ESO1 | $m^{7}Gppp^{Bn6}A_{m}pG$ | 2.47 | 79% | 91% | not detected [n] | 6C | C |
| 19 | 2089 nt | Fluc | $m^{7}GpppA_{m}pG$ | 3.61 | nd. | nd. | not detected [k] | S6 | B |
| 20 | 2089 nt | Fluc | $m^{7}Gppp^{Bn6}A_{m}pG$ | 2.53 | nd. | nd. | not detected [k] | S6 | B |
| 21 | 1695 nt | hA1AT | $m^{7}GpppA_{m}pG$ | 4.46 | nd. | nd. | not detected [o] | S7A-C | B |

| | | | | | | | | | |
|---|---|---|---|---|---|---|---|---|---|
| 22 | 1695 nt | hA1AT | $m^7Gppp^{Bn6}A_mpG$ | 4.3 | nd. | nd. | not detected [o] | S7A-C | B |
| 23 | 1695 nt | hA1AT | $m^7GpppA_mpG$ | 4.0 | nd. | nd. | not detected [o] | S7D | B |
| 24 | 1695 nt | hA1AT | $m^7Gppp^{Bn6}A_mpG$ | 4.33 | nd. | nd. | not detected [o] | S7D | B |
| 25 | 956 nt | Gluc | $m^7GpppA_mpG$ | 2.71 | nd. | nd. | not detected [k] | S10 | B |
| 26 | 956 nt | Gluc | $m^7Gppp^{m6}A_mpG$ | 2.7 | nd. | nd. | not detected [k] | S10 | B |
| 27 | 956 nt | Gluc | $m^7Gppp^{Bn6}A_mpG$ | 2.52 | nd. | nd. | not detected [k] | S10 | B |
| 28 | 1040 nt | hEPO | $m^7GpppA_mpG$ | 3.0 | nd. | nd. | not detected [k] | S13A | B |
| 29 | 1040 nt | hEPO | $m^7Gppp^{Bn6}A_mpG$ | 3.03 | nd. | nd. | not detected [k] | S13A | B |
| 30 | 2046 nt | Fluc | $m^7GpppA_mpG$ | 3.63 | nd. | nd. | not detected [p] | S13C | C |
| 31 | 2046 nt | Fluc | $m^7Gppp^{Bn6}A_mpG$ | 3.29 | nd. | nd. | not detected [p] | S13C | C |

[a] *In vitro* transcription (IVT) efficiency estimated by measuring the volume and the absorbance at 260 nm (NanoDrop One$^C$, ThermoFisher Scientific) of the eluate after initial purification step with oligo$(dT)_{25}$ resin (POROS$^{TM}$ Oligo $(dT)_{25}$ Affinity Resin, ThermoFisher Scientific). IVT efficiency is calculated as mRNA yield obtained from 1 µL of starting IVT volume; [b] efficiency of capping determined by ribozyme-based assay after initial step of mRNA purification with oligo$(dT)_{25}$ resin; [c] ribozyme-based analysis of the mRNA purity after completed purification process expressed as a percentage of capped fraction of the total purified mRNA; [d] dsRNA contamination of mRNA after completed purification process analyzed by dot-blot assay using anti-dsRNA J2 antibody (SCICONS) and compared to dsRNA standards (Abnova or New England Biolabs); [e] information in this column indicates the figures, where analyzed data is shown; [f] information in this column indicates which protocol was used for *in vitro* transcription and mRNA purification (see Experimental section); [g] dsRNA content in 250 ng of the analyzed mRNA sample is below detection limit of the method (0.31‰), which is determined based on the dsRNA standard in the range of 0.078-10 ng (Abnova); [h] amount of dsRNA detected in 2500 ng of the analyzed mRNA after completed purification process; [i] contamination with dsRNA expressed as a percentage of dsRNA in 2500 ng of analyzed mRNA sample; [j] dsRNA content in 2500 ng of the analyzed mRNA sample is below detection limit of the method (0.05‰), which is determined based on the dsRNA standard in the range of 0.062-8 ng (New England Biolabs); [k] dsRNA content in 250 ng of the analyzed mRNA sample is below detection limit. 25 ng and 50 ng of dsRNA standard (New England Biolabs) was used as the reference; [l] dsRNA content in 2500 ng of the analyzed mRNA sample is below detection limit of the method (0.03‰), which is determined based on the dsRNA standard in the range of 0.078-10 ng (Abnova); [m] dsRNA content in 2500 ng of the analyzed mRNA sample is below detection limit of the method (0.156‰), which is determined based on the dsRNA standard in the range of 0.391-50 ng (New England Biolabs); [n] dsRNA content in 2500 ng of the analyzed mRNA sample is below detection limit of the method (0.125‰), which is determined based on the dsRNA standard in the range of 0.156-20 ng (New England Biolabs); [o] dsRNA content in 2500 ng of the analyzed mRNA sample is below detection limit of the method (1.25‰), which is determined based on the dsRNA standard in the range of 0.391-50 ng (New England Biolabs); [p] dsRNA content in 250 ng of the analyzed mRNA sample is below detection limit of the method (6.25‰), which is determined based on the dsRNA standard in the range of 0.391-50 ng (New England Biolabs).

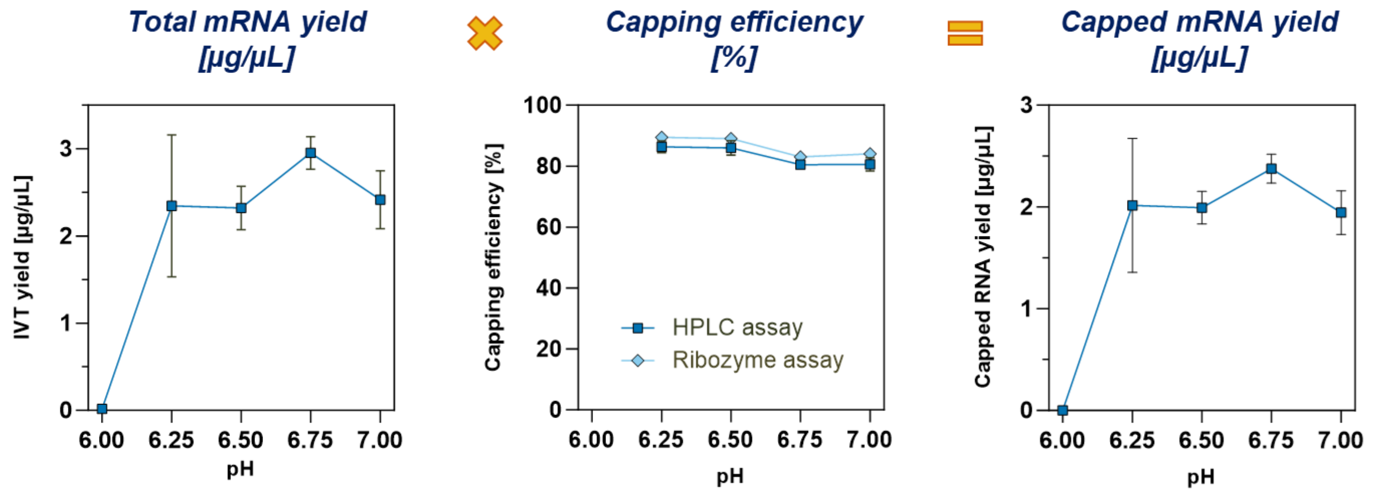

**Figure S1. Capping efficiencies (determined by two methods) and IVT yields (determined spectrophotometrically for the samples purified using oligo(dT)$_{25}$ resin) as a function of pH in the range of 6.0-7.0.** IVT reactions were performed in the presence of 25 mM $MgCl_2$, 40 ng/µL template, 5 mM ATP, CTP, UTP, 4 mM GTP and 10 mM *AvantCap*.

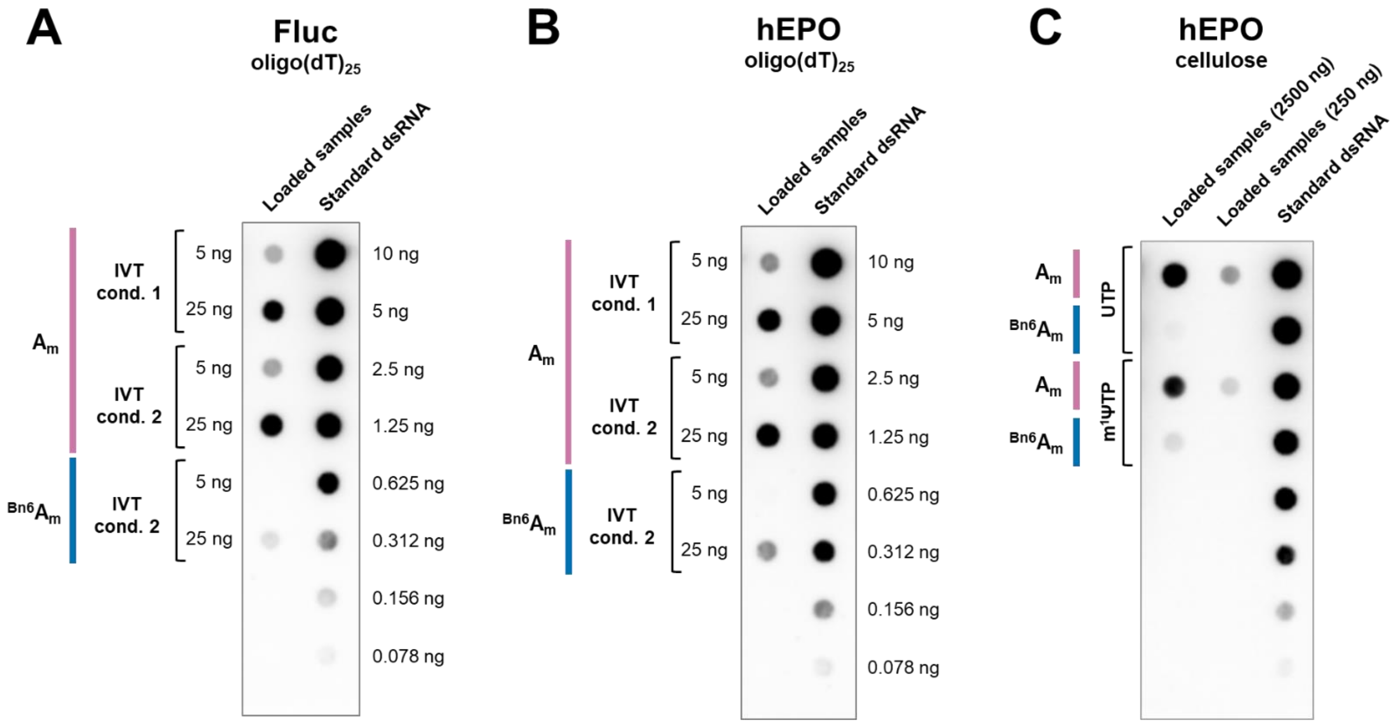

**Figure S2. Comparison of dsRNA content in the initially purified mRNA samples co-transcriptionally capped with m$^7$GpppA$_m$pG (A$_m$) or *AvantCap* ($^{Bn6}$A$_m$).** mRNA encoding (**A**) firefly luciferase (Fluc) or (**B, C**) human erythropoietin (hEPO) was purified by (**A, B**) oligo(dT)$_{25}$ affinity chromatography or (**C**) using cellulose. Each mRNA sample was immobilized on the nylon membrane (Hybond$^{TM}$-N$^+$, Amersham$^{TM}$), incubated with dsRNA-specific J2 antibody (SCICONS), followed by the incubation with secondary anti-mouse HRP-conjugated antibody (Cell Signaling Technology) and HRP substrate (ECL™ Prime Western Blotting Detection Reagents). Chemiluminescence signal was detected using ImageQuant 800 (Amersham$^{TM}$) imaging system. dsRNA content was semi-quantitative determined using ImageQuantTL software by the comparison of the detected signals to the measured intensities of standard dsRNA (Abnova).

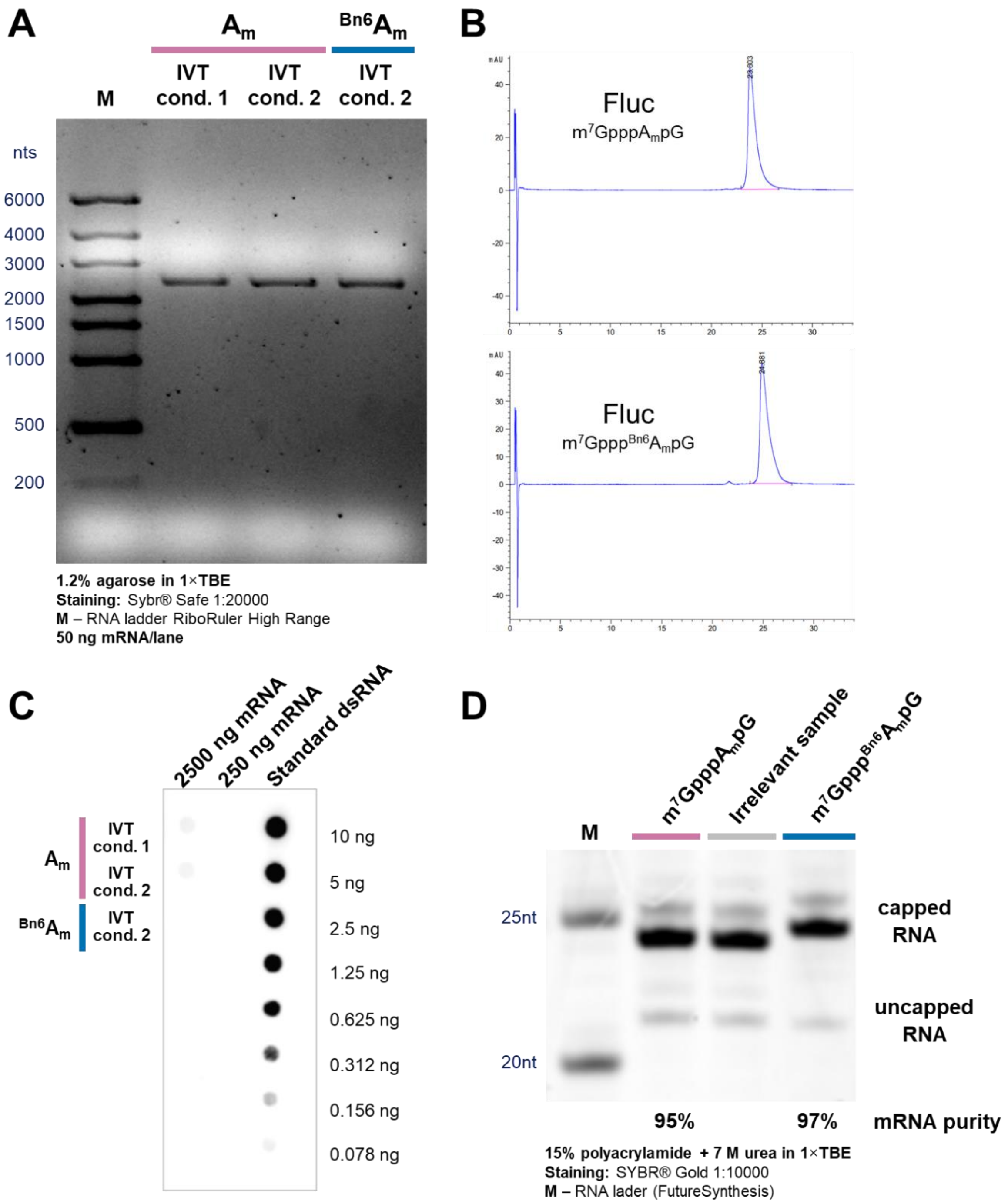

**Figure S3. Representative quality control of the final samples of mRNA encoding firefly luciferase (Fluc) after whole purification process.** (**A**) Integrity analysis: 50 ng of each mRNA was analysed using 1.2% agarose gel in 1×TBE. (**B**) Purity analysis by RP-HPLC: 1 µg of mRNA was applied on bioZen™ 2.6 µm Oligo LC column (Phenomenex) and separated using linear gradient of acetonitrile (10-20%) in 0.1 M TEAA pH 7.0 at 55°C. (**C**) Detection of dsRNA: 0.25 and 2.5 µg of mRNA sample was immobilized on the nylon membrane (Hybond™-N$^{+}$, Amersham™), incubated with dsRNA-specific J2 antibody (SCICONS) and secondary anti-mouse HRP-conjugated antibody (Cell Signaling Technology), followed by chemiluminescence signal detection (ImageQuant 800, Amersham™). (**D**) Capping efficiency: densitometric quantification of bands intensities corresponding to 5´ mRNA fragments cleaved by 5´UTR complementary ribozyme. 15% polyacrylamide gel with 7 M urea and 1× TBE was used for separation of digested RNA fragments.

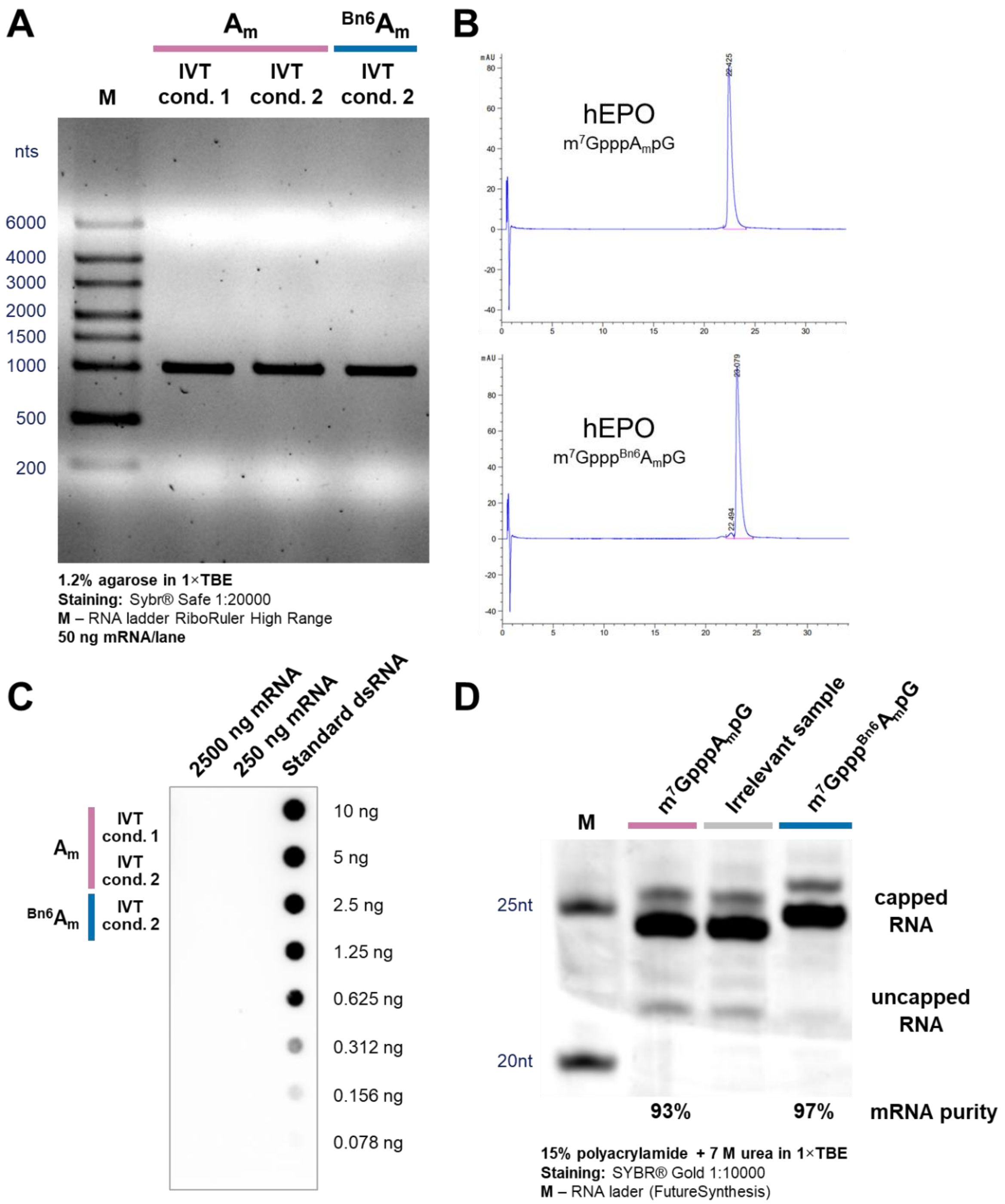

**Figure S4. Representative quality control of the final samples of mRNA encoding human erythropoietin (hEPO) after whole purification process.** (**A**) Integrity analysis: 50 ng of each mRNA was analysed using 1.2% agarose gel in 1×TBE. (**B**) Purity analysis by RP-HPLC: 1 µg of mRNA was applied on bioZen™ 2.6 µm Oligo LC column (Phenomenex) and separated using linear gradient of acetonitrile (10-20%) in 0.1 M TEAA pH 7.0 at 55°C. (**C**) Detection of dsRNA: 0.25 and 2.5 µg of mRNA sample was immobilized on the nylon membrane (Hybond™-N$^{+}$, Amersham™), incubated with dsRNA-specific J2 antibody (SCICONS) and secondary anti-mouse HRP-conjugated antibody (Cell Signaling Technology), followed by chemiluminescence signal detection (ImageQuant 800, Amersham™). (**D**) Capping efficiency: densitometric quantification of bands intensities corresponding to 5´ mRNA fragments cleaved by 5´UTR complementary ribozyme. 15% polyacrylamide gel with 7 M urea and 1× TBE was used for separation of digested RNA fragments.

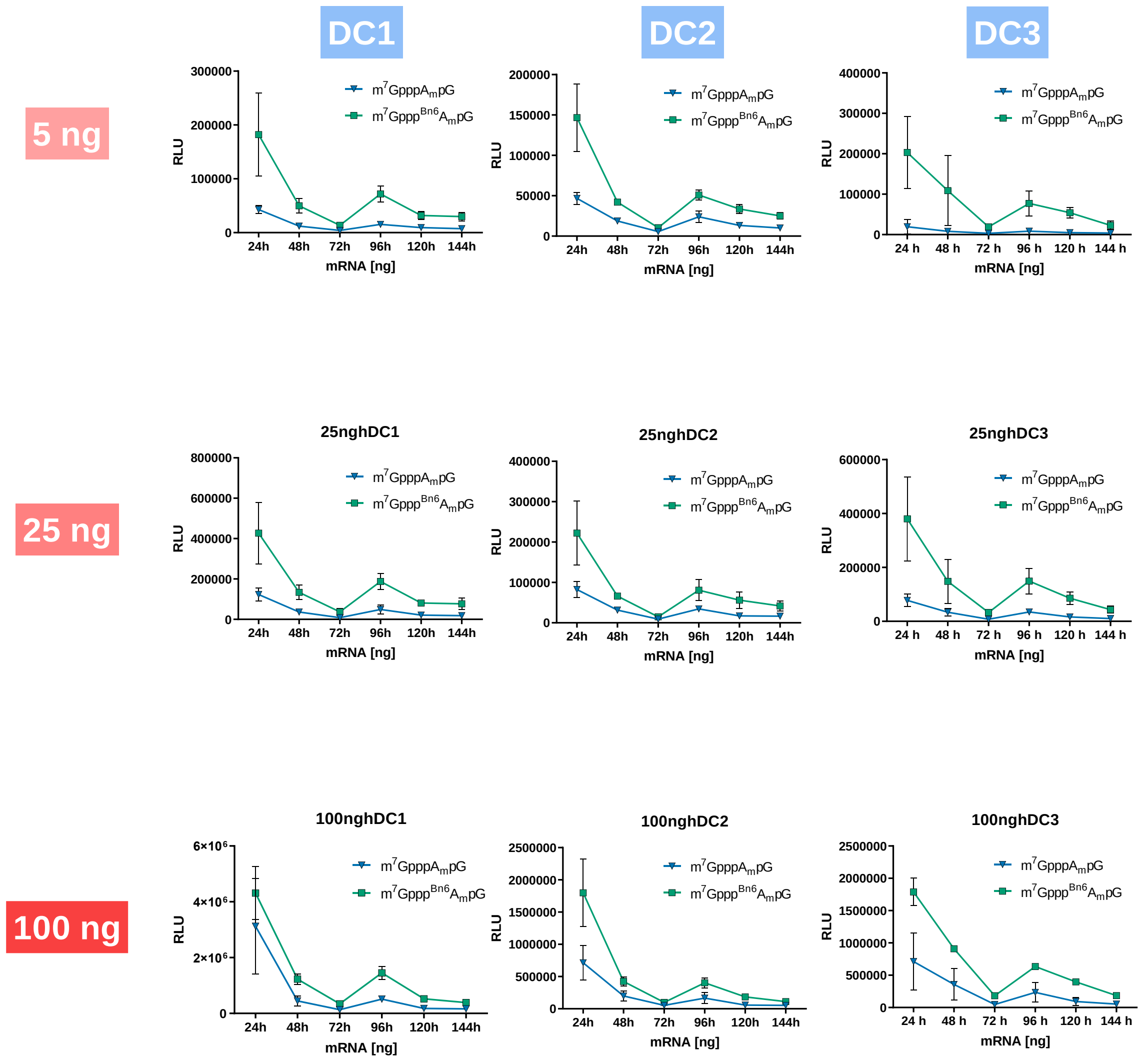

**Figure S5. Time-dependent expression profiles of differently capped Gluc mRNAs in human dendritic cells.** *Gaussia* luciferase (Gluc)-dependent luminescence of human monocyte-derived dendritic cells (hDCs) transfected with 5, 25 or 100 ng of Gluc mRNA;

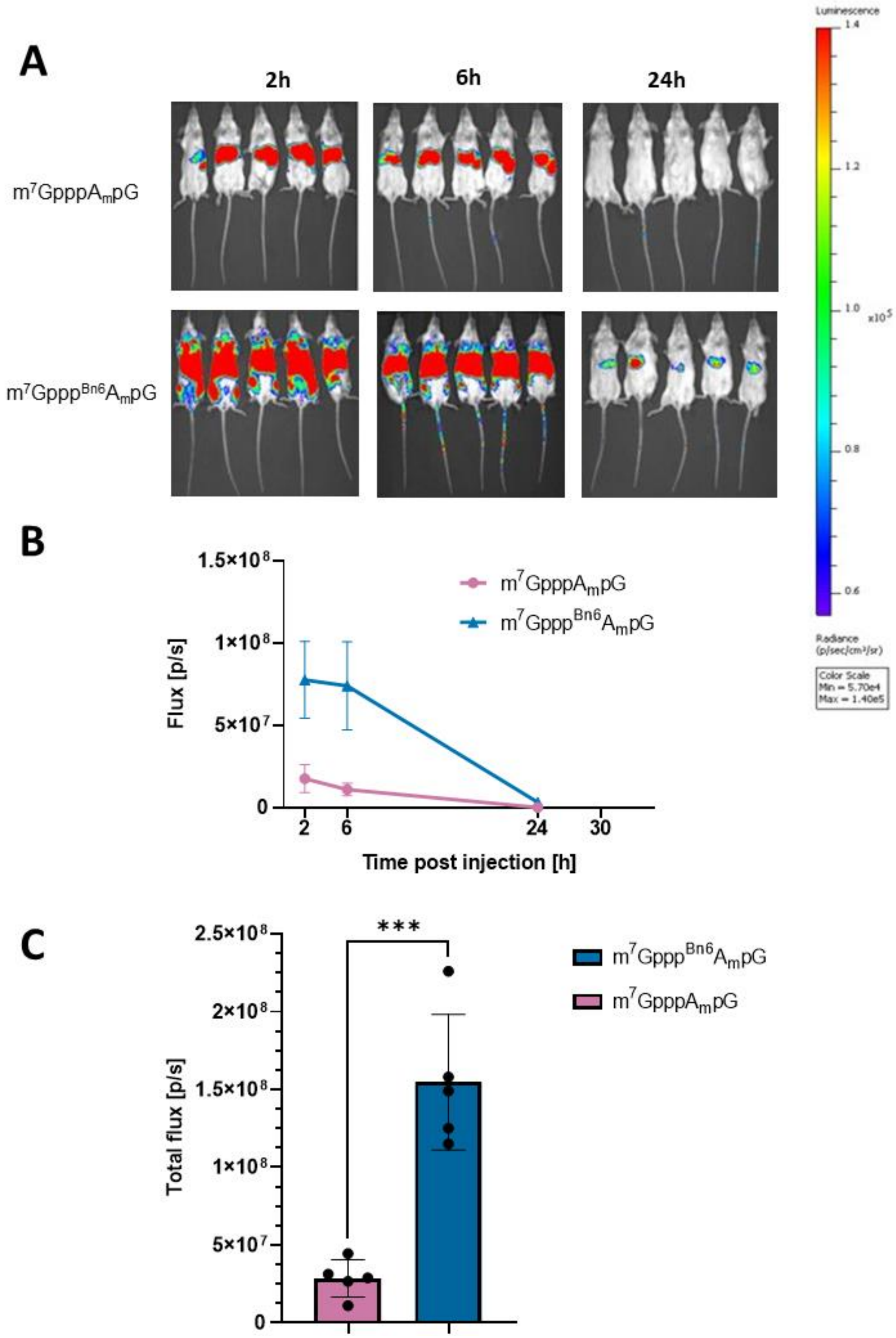

**Figure S6. mRNA capped with *AvantCap* ($m^7Gppp^{Bn6}A_mpG$) yields superior protein expression in vivo.** Intravital bioluminescence at 2 h, 6 h and 24 h in BALB/c mice injected intravenously with 10 µg of firefly luciferase (Fluc) encoding mRNAs, capped with $m^7GpppA_mpG$ or $m^7Gppp^{Bn6}A_mpG$, and formulated using TransIT®. (**A**) Raw bioluminescence images of mice with the scale for bioluminescent signals at the right. (**B**) Flux [p/s] mean values in time ± SD, n=5. (**C**) Total flux (sum of flux values for each time point, [p/s]) means ± SD, n=5, ***P<0.001, two-tailed unpaired *t* test.

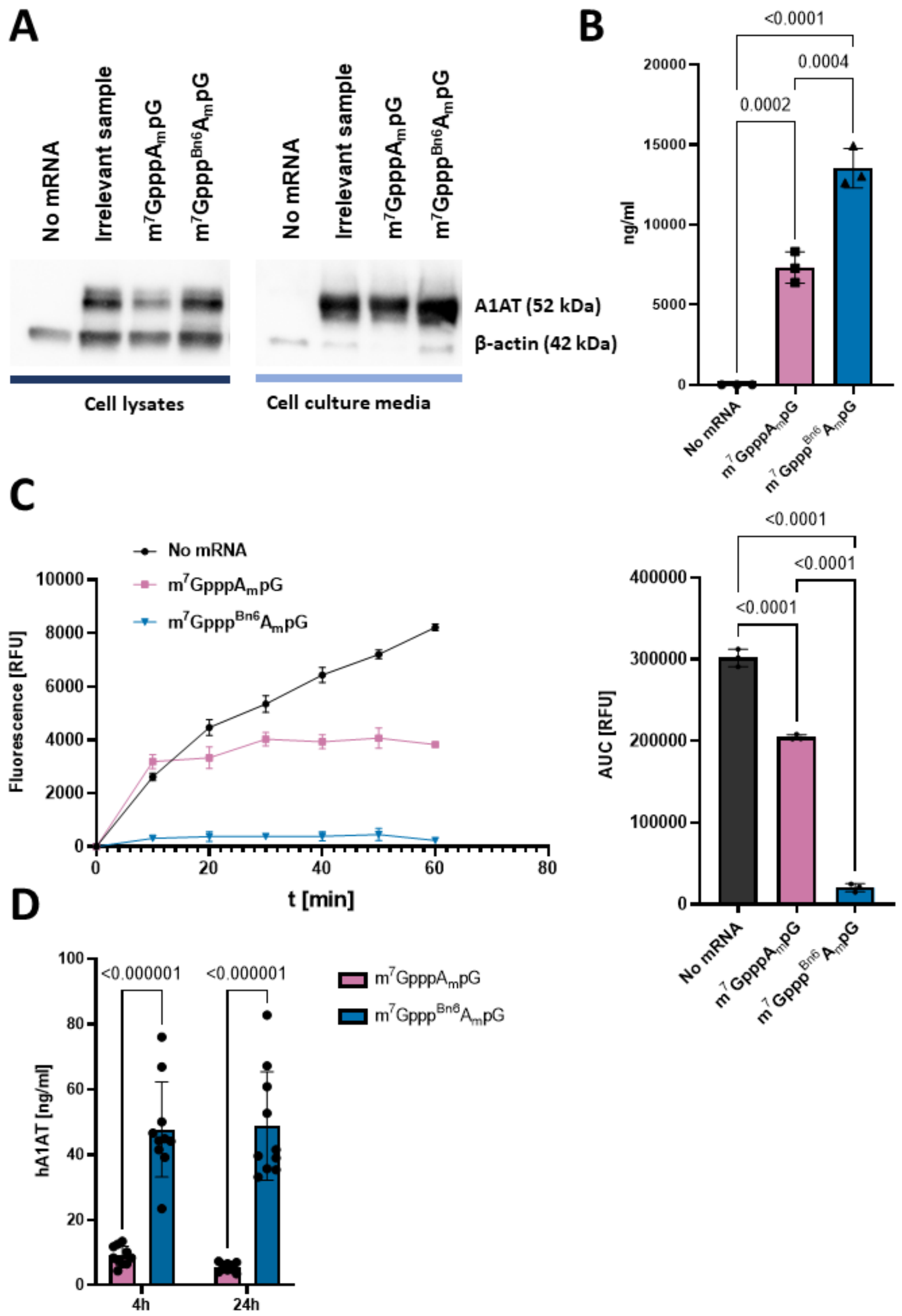

**Figure S7. In vitro and in vivo translational efficacy of differentially capped mRNA for human α1-antitrypsin (hA1AT).** (**A**) Amounts of human hA1AT protein in HEK293T cell lysates and culture media 24 h post transfection with differentially capped mRNA. Western blotting, β-actin levels served as loading control for the cell lysates; negative control - cells treated with LipofectAMINE Messenger Max only (no mRNA lanes). The amount of lysate applied in each well corresponded to the amount of culture medium used for this assay. (**B**) 1 × $10^4$ human lung cancer cells (A549) were transfected with 50 ng of hA1AT-encoding m$^7$GpppA$_m$pG or m$^7$Gppp$^{Bn6}$A$_m$pG-capped mRNA using LipofectAMINE Messenger Max. A1AT concentration in the culture medium 24 h post transfection measured with ELISA. Data show means±SD, n=3, one-way ANOVA with Tukey's multiple comparisons test. (**C**) Results of EnzChek Elastase Assay (ThermoFisher Scientific) run using culture medium from A549 cells 24 h post transfection with hA1AT mRNA or LipofectAMINE Messenger Max only (no mRNA). (**left**) Elastase activity in time measured as relative fluorescence units [RLU] of the fluorescent product of the enzymatic reaction. (**right**) Area under the curve [AUC] mean values ±SD, n=3, one-way ANOVA with Tukey's multiple comparisons test. The signal from samples incubated with culture medium from hA1AT mRNA-transfected cells with different 5'caps is reduced compared to non-transfected cells due to hA1AT-dependent inhibition of elastase enzymatic activity. (**D**) C57BL/6 mice were injected intravenously with 10 µg of hA1AT-encoding m$^7$GpppA$_m$pG or m$^7$Gppp$^{Bn6}$A$_m$pG-capped mRNA formulated in TransIT®. Serum hA1AT concentration at 4 and 24 h post injection measured with ELISA. Data show means±SD, n=10, multiple unpaired *t* tests.

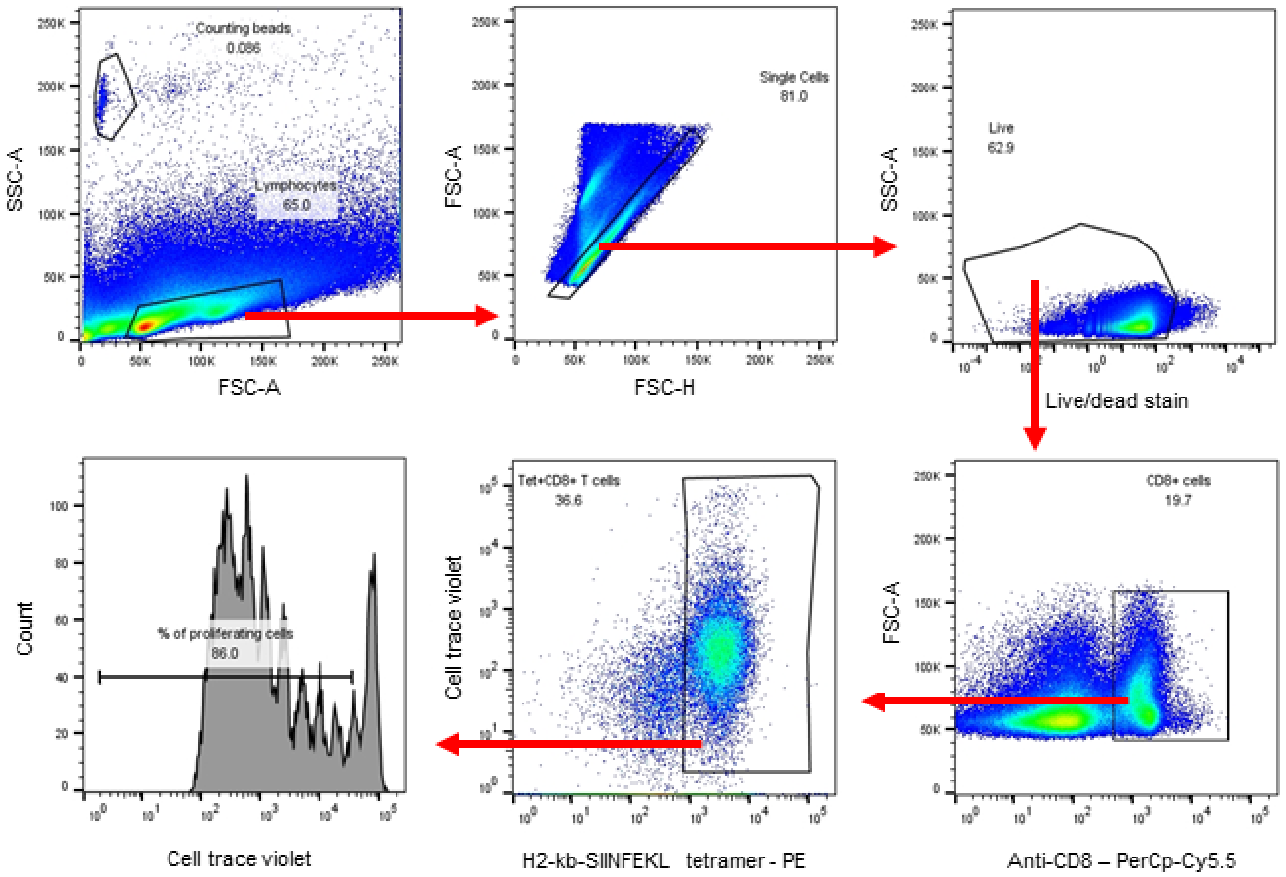

**Figure S8. Flow cytometry gating strategy for identification of proliferating OT-I T cells in the spleen.**

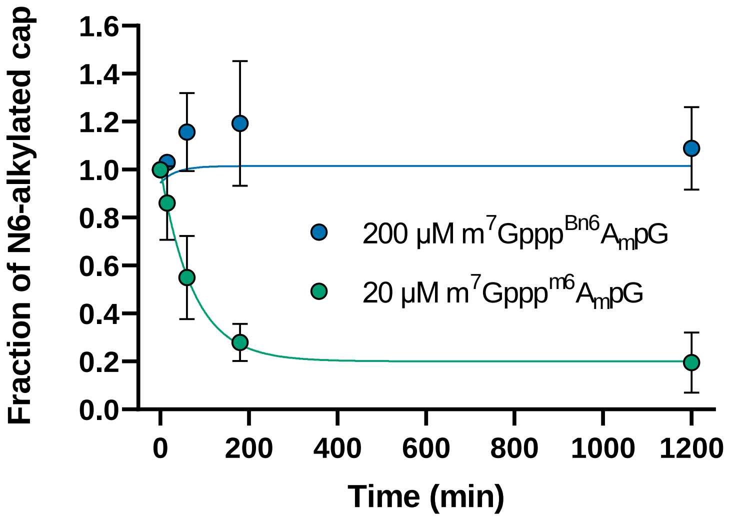

**Figure S9. FTO assay for 200 µM *AvantCap* (m$^7$Gppp$^{Bn6}$A$_m$pG).** $^{Bn6}$A$_m$- or $^{m6}$A$_m$-containing cap analogs were incubated with FTO and the amount of *N6*-alkylated cap remaining in the mixture was assessed by RP-HPLC-MS. Reaction conditions: 20 µM or 200 µM cap and 2 µM FTO in 50 mM HEPES pH 7.0, containing 150 mM KCl, 75 µM Fe(II), 300 µM 2-oxoglutarate, and 2 mM ascorbic acid.

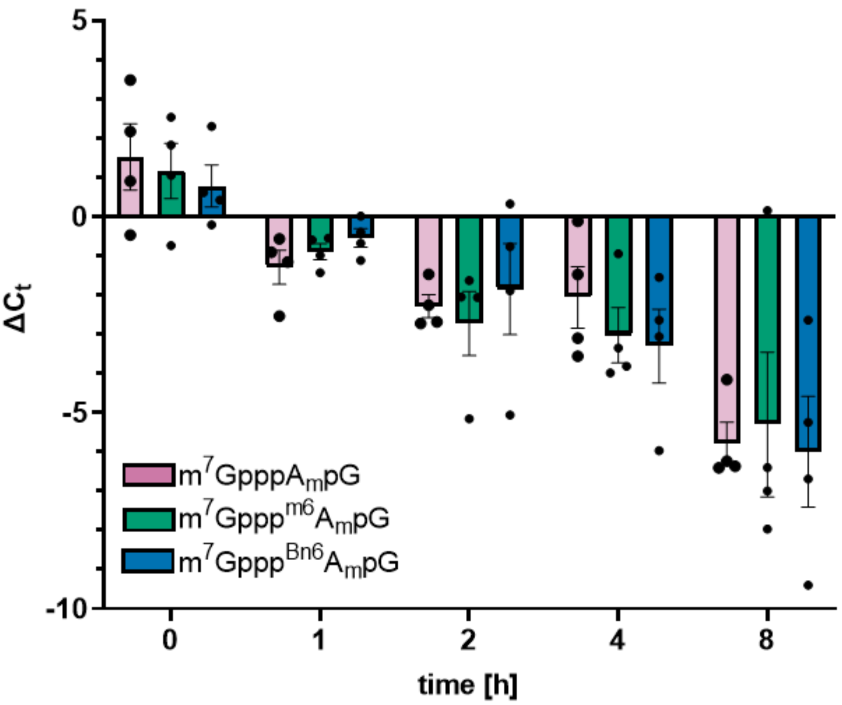

**Figure S10. *AvantCap* (m$^7$Gppp$^{Bn6}$A$_m$pG) does not stabilize mRNA in human cells.** HEK293T cells were electroporated at time 0 with 1 µg of m$^7$GpppA$_m$pG, m$^7$Gppp$^{m6}$A$_m$pG or m$^7$Gppp$^{Bn6}$A$_m$pG-capped mRNA for *Gaussia* luciferase (Gluc) per 1 mln of the cells. At indicated time points mRNA amount in the cells was evaluated with qRT-PCR. Gluc $\Delta c_t$ mean values ± SE, N=4, Kruskal-Wallis test with multiple comparisons showed no statistically significant differences between groups. Human β-actin was used as a housekeeping gene for normalization.

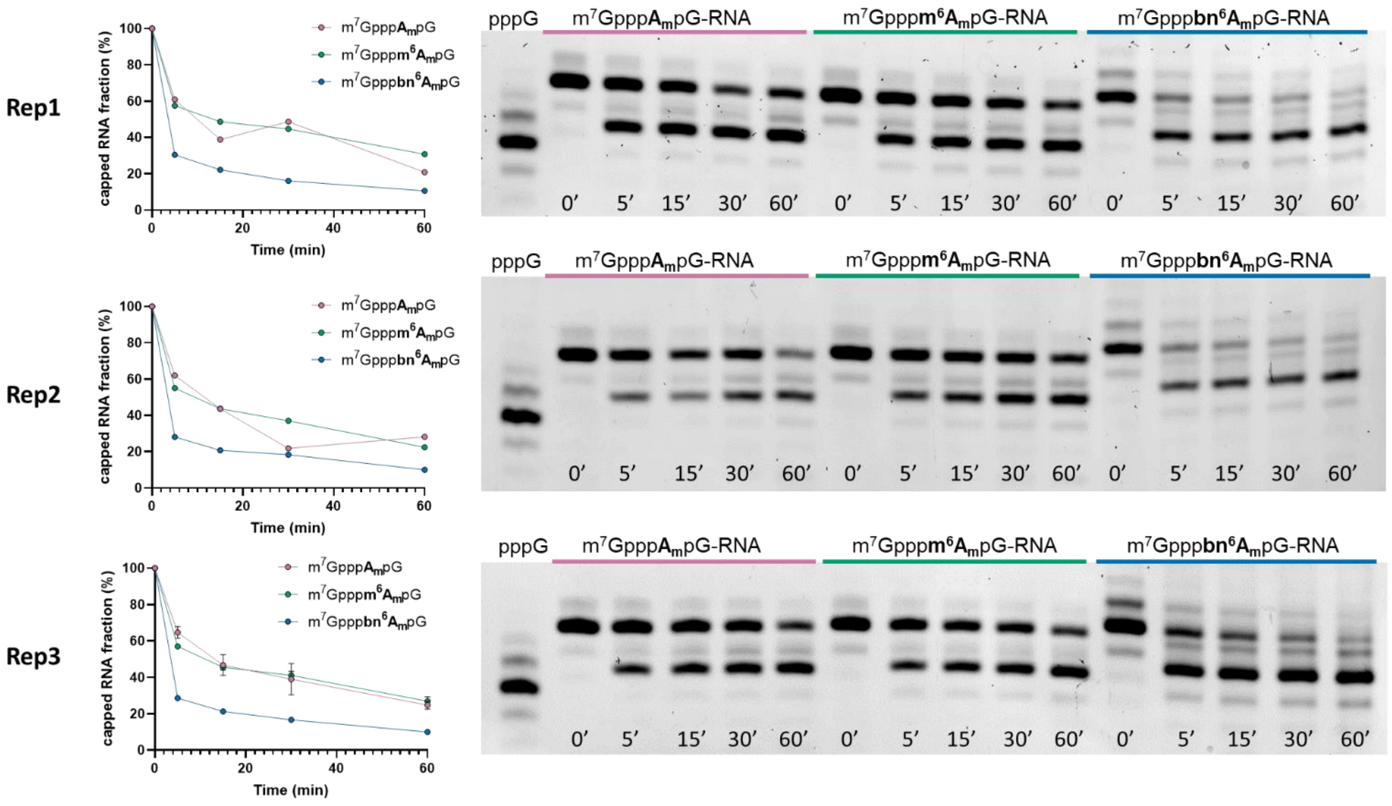

**Figure S11. Susceptibility of short RNAs to decapping by PNRC2-hDcp1/Dcp2 complex in vitro.** Individual data for all replicates.

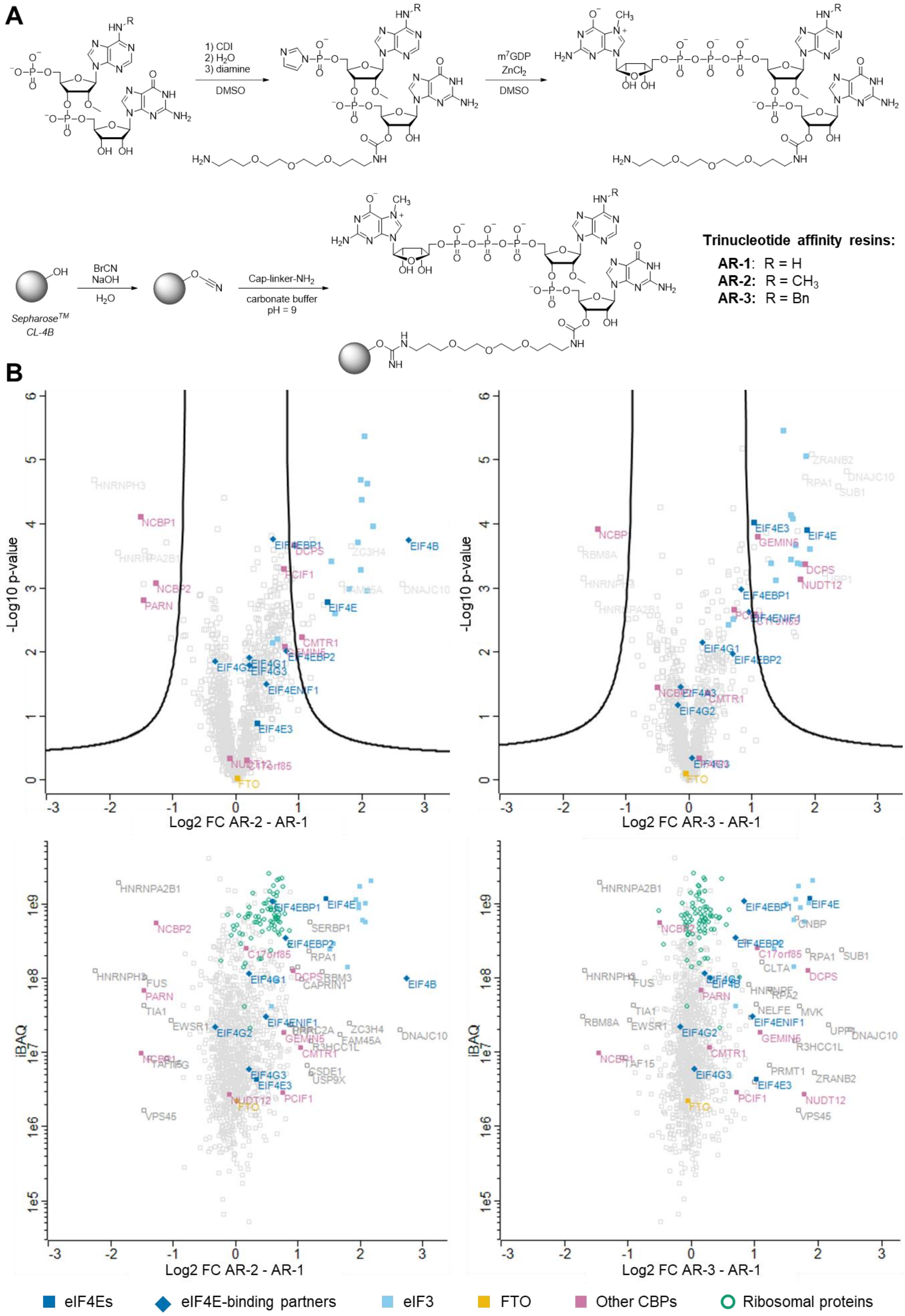

**Figure S12. Pull-down assay with protein extract from HEK293F cells. A)** Synthetic pathway to trinucleotide cap analogs immobilized on Sepharose beads. **B)** Volcano plots (**top**) displaying the log2 fold change (log2FC) against the t-test-derived −log10 statistical p-value (-Log p-value) for the protein groups in the eluates from trinucleotide cap affinity resins AR-2/AR-1 (n = 3) and AR-2/AR-1 (n = 3). Student's t-tests (two-sided, unpaired) were performed to assess binding preferences of protein groups to AR-2 and AR-3 resins relative to AR-1 resin. Dot plots (**bottom**) displaying the log2 fold change values against the mean iBAQ values (that provide a rough estimate of the relative abundance of proteins in the eluates).

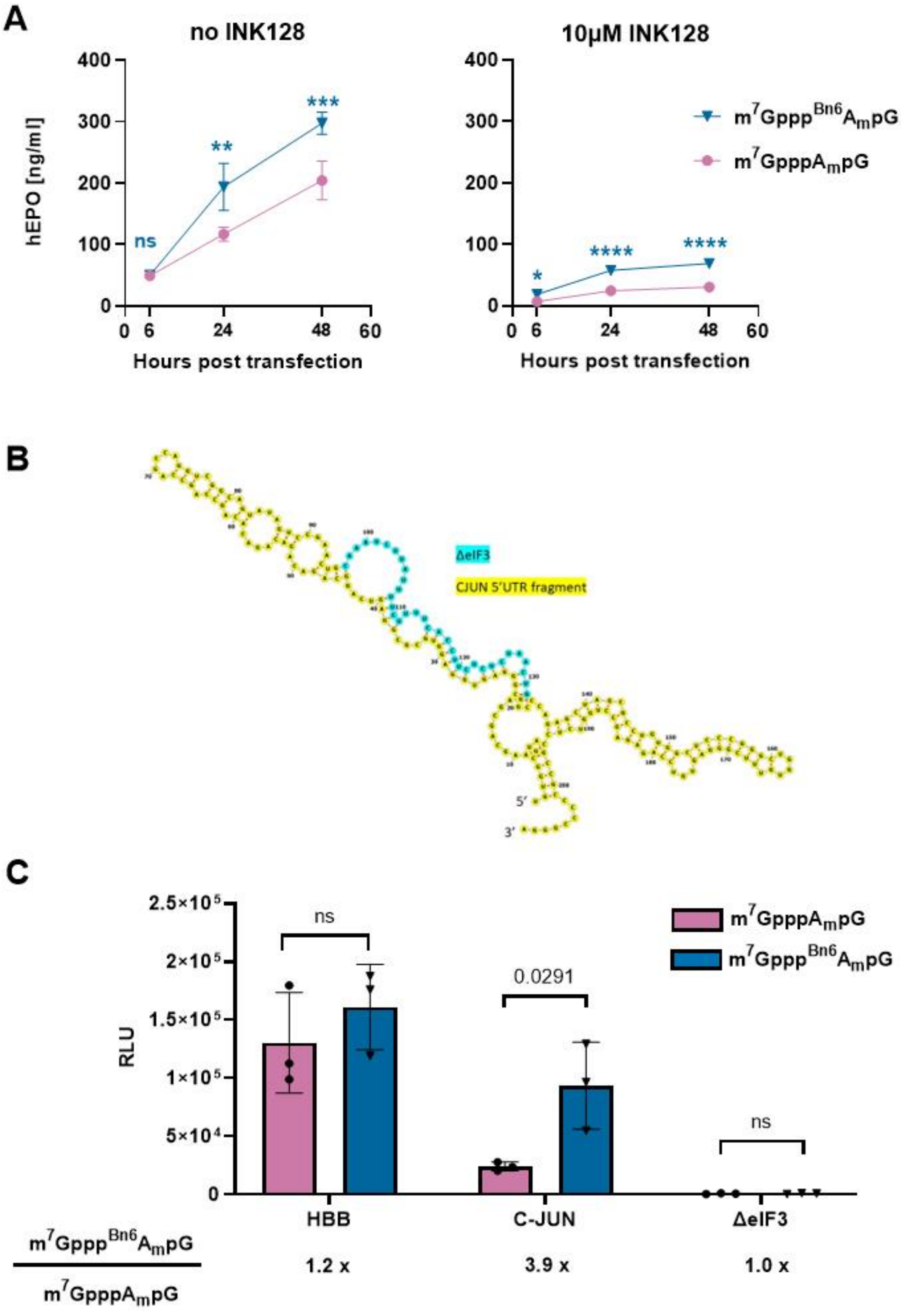

**Figure S13. Translation of mRNA with *AvantCap* ($m^7Gppp^{Bn6}A_mpG$) is partially initiated by eIF3d-dependent mechanism.** (**A**) Human monocyte-derived dendritic cells ($5 \times 10^4$) were pretreated for 4 h with 10 μM INK128 (mTOR inhibitor blocking eIF4E-dependent mRNA translation initiation) or DMSO at concentration corresponding to 10 μM INK128 (no INK128). Next, the cells were transfected with 50 ng of $m^7GpppA_mpG$ or $m^7Gppp^{Bn6}A_mpG$-capped mRNA for hEPO using LipotectAMINE Messenger Max. hEPO concentration in culture medium 6, 24 and 48 h post transfection measured with ELISA, mean ± SD, n-3; ns-not significant, ** $P<0.01$, *** $P<0.001$, ****$P<0.0001$, one-way ANOVA with Šidák's multiple comparisons test. (**B**) 2D structure of human CJUN 5'UTR fragment showing eIF3 binding site (cyan). Structure created using RNAfold WebServer (http://rna.tbi.univie.ac.at/; minimum free energy prediction) and Forna visualization tool (http://rna.tbi.univie.ac.at/forna/). (**C**) Human monocyte-derived dendritic cells ($5 \times 10^4$) were transfected with 50 ng of $m^7GpppA_mpG$ or $m^7Gppp^{Bn6}A_mpG$-capped mRNA for Fluc with 5'UTR from HBB, c-JUN or truncated c-JUN lacking eIF3 binding fragment (ΔeIF3) using LipotectAMINE Messenger Max. Relative luminescence units [RLU] 18 h post transfection; mean ± SD, n=3, ns – not significant, one-way ANOVA with Tukey's multiple comparisons test. $m^7Gppp^{Bn6}A_mpG$-capped mRNA to $m^7GpppA_mpG$ -capped mRNA ratio for each of the tested 5'UTR sequences is shown below the graph.
