## Supplementary figures and images for "Trinucleotide mRNA cap analog N6-benzylated at the site of posttranscriptional ^m6^Am mark facilitates mRNA purification and confers superior translational properties in vitro and in vivo"

### Graphical summary

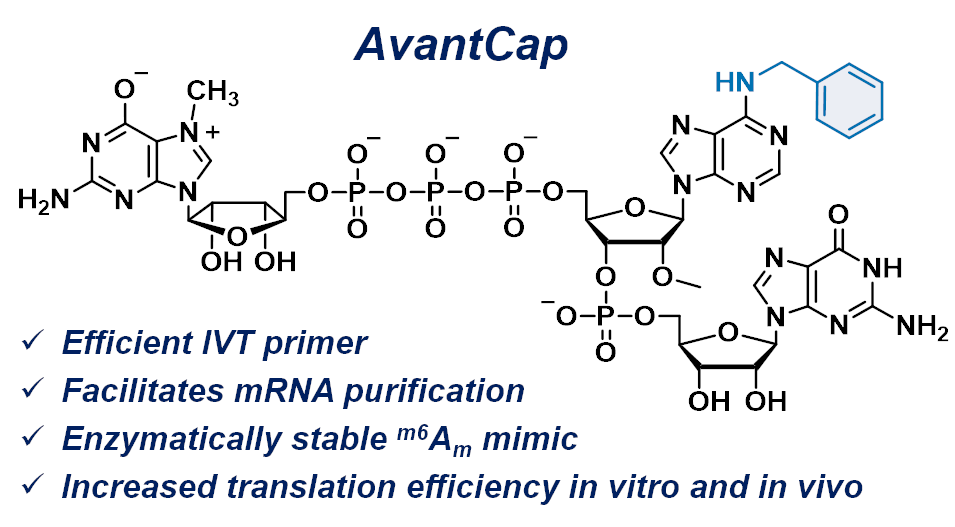
